# Biomechanical control of meiotic chromosomal bouquet and germ cell morphogenesis by the zygotene cilium

**DOI:** 10.1101/2021.02.08.430249

**Authors:** Avishag Mytlis, Vineet Kumar, Qiu Tao, Rachael Deis, Neta Hart, Karine Levy, Markus Masek, Amal Shawahny, Adam Ahmad, Hagai Eitan, Farouq Nather, Shai Adar-Levor, Ramon Y. Birnbaum, Natalie Elia, Ruxandra Bachmann-Gagescu, Sudipto Roy, Yaniv M. Elkouby

## Abstract

The hallmark of meiosis is chromosomal pairing and synapsis via synaptonemal complexes, but chromosomal pairing also depends on cytoplasmic counterparts that tether and rotate telomeres on the nuclear envelope. Telomeres slide on perinuclear microtubules, shuffling chromosomes and mechanically driving their homology searches. Pull of telomeres towards the centrosome drives formation of the “zygotene chromosomal bouquet”. These telomere dynamics are essential for pairing and fertility, and the bouquet, discovered in 1900, is universally conserved. Nevertheless, how cytoplasmic counterparts of bouquet formation are mechanically regulated has remained enigmatic. Here, we report the “zygotene cilium” - a previously unrecognized cilium, in oocytes. We show in zebrafish that this cilium specifically connects to the bouquet centrosome, constituting a cable system of the cytoplasmic bouquet machinery. Furthermore, zygotene cilia extend throughout the germline cyst, a conserved germ cell organization. Using multiple ciliary mutants and laser-induced excision, we demonstrate that the zygotene cilium is essential for chromosomal bouquet and synaptonemal complex formation, germ cell morphogenesis, ovarian development and fertility. Mechanistically, we provide evidence that the cilium functions at least partly via anchoring the bouquet centrosome in order to counterbalance telomere rotation and pulling. We also show that the zygotene cilium is conserved in both male and female meiosis in zebrafish, as well as in mammals. Our work uncovers the novel concept of a cilium as a critical player in meiosis and sheds new light on reproduction phenotypes in ciliopathies. We propose a cellular paradigm that cilia can control chromosomal dynamics.

## Introduction

Meiosis is a cellular program essential for the production of haploid gametes that is conserved in sexually reproducing organisms. During prophase stages of meiosis, induced double strand breaks must be repaired, and their repair via homologous recombination requires the pairing of homologous chromosomes and formation of the synaptonemal complex. The synaptonemal complex is comprised of synaptonemal complex proteins (Sycp1-3) that assemble between chromosomes and along their axes in a stereotypic manner (*1*). Together with meiotic cohesins, these connect homologous chromosomes and hold them together for the execution of recombination (*1, 2*).

In females, meiosis begins in the developing ovary. In mammals, these prophase events are executed in the fetal ovary and females are born with a set pool of primordial and primary follicles. In zebrafish, prophase is initiated during post-embryonic development, but the program of meiosis and generally oogenesis are highly conserved between zebrafish and mouse (*3*). In both species and most vertebrates, oogenesis begins after germline stem cells give rise to oocyte mitotic precursors, called oogonia (*3*). Oogonia undergo several rounds of incomplete divisions, where cytoplasmic bridges are retained between sister cells, forming the germline cyst, a conserved organization of germ cells (*4–8*). In the cyst, oogonia are surrounded by somatic pre-granulosa cells (*4, 6, 9*).

The induction of meiosis transforms oogonia into differentiating oocytes, which undergo the leptotene and zygotene prophase stages synchronously within the cyst (*5–7, 10*). Oocytes then switch from the cyst organization and form the primordial follicle by the pachytene stage (*3, 4, 11*). During prophase, double-strand breaks, homology searches and initial pairing and formation of the synaptonemal complex are executes at leptotene-zygotene stages, and homologous recombination completes in pachytene (*3, 4, 12-15*). Oocytes then continue to grow in the developing follicle, while arrested at the diplotene stage of meiosis, which will later resume during ovulation (*3*).

Recent breakthroughs have dissected the structure of synaptonemal complexes and how synaptonemal complex proteins and meiosis-specific cohesins assemble to connect homologous chromosomes (*1, 2, 16-21*). However, the nuclear events of meiosis occur in the cellular context of a differentiating gamete, where chromosomal pairing depends on cytoplasmic counterparts. In pairing, telomeres perform a unique function: At the leptotene stage, telomeres tether to Sun/Kash complexes on the nuclear envelope (NE), which associate with perinuclear microtubules (MTs) via dynein (*22–29*). This facilitates telomere rotations around the NE during leptotene-zygotene stages (Fig. S1A), which in turn shuffle chromosomes, driving their search for homologs. The perinuclear MTs at these stages emanate from the centrosome MT organizing center (MTOC) (*4, 28, 30*) (Fig. S1A). Ultimately, rotating telomeres are pulled towards the centrosome and cluster on the centrosome side of the NE, while looping their chromosomes to the other side - a configuration called the zygotene chromosomal bouquet (*3, 31*) (Fig. S1A). These telomere dynamics are essential for synapsis, homologous recombination, and fertility (*22–29, 32*). Telomere clustering is thought to stabilize initial pairing between homologous chromosomes, and in zebrafish the bouquet was proposed a hub for chromosomal pairing (*33*). It was shown that double strand breaks cluster at sub-telomeric regions and the synaptonemal complex proteins Sycp3 and Sycp1 first load adjacent to clustered telomeres specifically at the bouquet stages, from which they extend along chromosomal axes (*33, 34*) (Fig. S1A).

The bouquet configuration was discovered in 1900 (*31*), and is conserved from yeast to mammals (*22–29*). However, how the cytoplasmic counterparts of bouquet formation are mechanically regulated in rotating telomeres is unknown. It is unclear how rotation forces are generated, counterbalanced to allow movements, and then halted at the correct stage.

## Results

We previously characterized the nuclear and cytoplasmic components of the bouquet and its formation in zebrafish oogenesis (*4, 35*) (see ‘Staging prophase oocytes’ in the Methods section, and Fig. S1B). To obtain a better temporal resolution of bouquet formation, we performed live time-lapse analysis of bouquet chromosomal rotation in cultured ovaries (*35*) (Movie S1, Fig. S2). This analysis revealed rapid rotational movements, specifically in leptotene-zygotene oocytes, in contrast with oogonia and somatic pre-granulosa follicle cell nuclei, which do not execute meiotic chromosomal pairing (Fig. S2, Movie S1). These rapid rotational movements suggest the underlying action of substantial mechanical forces. Rotating telomeres then cluster, strictly juxtaposing the centrosome (*4, 30, 34*) (Fig. S1A). Live recording of the centrosome during bouquet chromosomal rotations showed that it vibrates in place, remaining fixed at the same position in the cytoplasm (Movie S2). Thus, like in mice (*30*), telomeres rotate and are pulled by MTs toward the centrosome to cluster. These observations suggest that ***1)*** MTs and the centrosome are subjected to mechanical strain during rotations, and ***2)*** the centrosome is likely to be physically anchored.

### The zygotene cilium

We detected elongated fiber-like structures of acetylated tubulin that emanated from the centrosome in bouquet stage oocytes with clustered telomeres (*4*). To our knowledge, such structures, which are highly reminiscent of cilia, have not been described before in oocytes of any species. A cilium is a MT-based organelle, comprised of an axoneme with stereotypically organized acetylated microtubule doublets that grow from the mother centriole (basal body) and extend extracellularly (*36, 37*). We reasoned that a stage-specific cilium-like structure is an intriguing candidate regulator of bouquet mechanics.

Several lines of evidence demonstrated that the bouquet acetylated tubulin structures are cilia. ***1)*** The specific ciliary membrane marker Arl13b (*38, 39*) colocalized with the acetylated tubulin (AcTub)-positive structures (Fig. 1A, C). ***2)*** Tubulin glutamylation (GluTub), another post-translational MT modification of ciliary axonemes, showed similar structures (Fig. 1B), that co-localized with the AcTub signal (Fig. 1C). ***3)*** Transgenic Arl13b-GFP in zygotene oocytes confirmed consistent ciliary structures in live ovaries (Fig. S3B). ***4)*** Transmission electron microscopy (TEM) of ovaries identified structures that correspond to ciliary axonemes with typical MT doublets that are found in spaces between cell membranes of adjacent zygotene oocytes (Fig. 1D, bottom panels, Fig. S3A). Within oocytes, we identified the basal body, transition zone, and axoneme extending extracellularly through the ciliary pocket (Fig. 1D, top panels, Fig. S3A). Serial Block-Face Scanning Electron Microscopy (SBF-SEM, discussed below) further confirmed axonemal structures that emanated from basal bodies in prophase oocytes (Movies S3-5).

**Figure 1.**
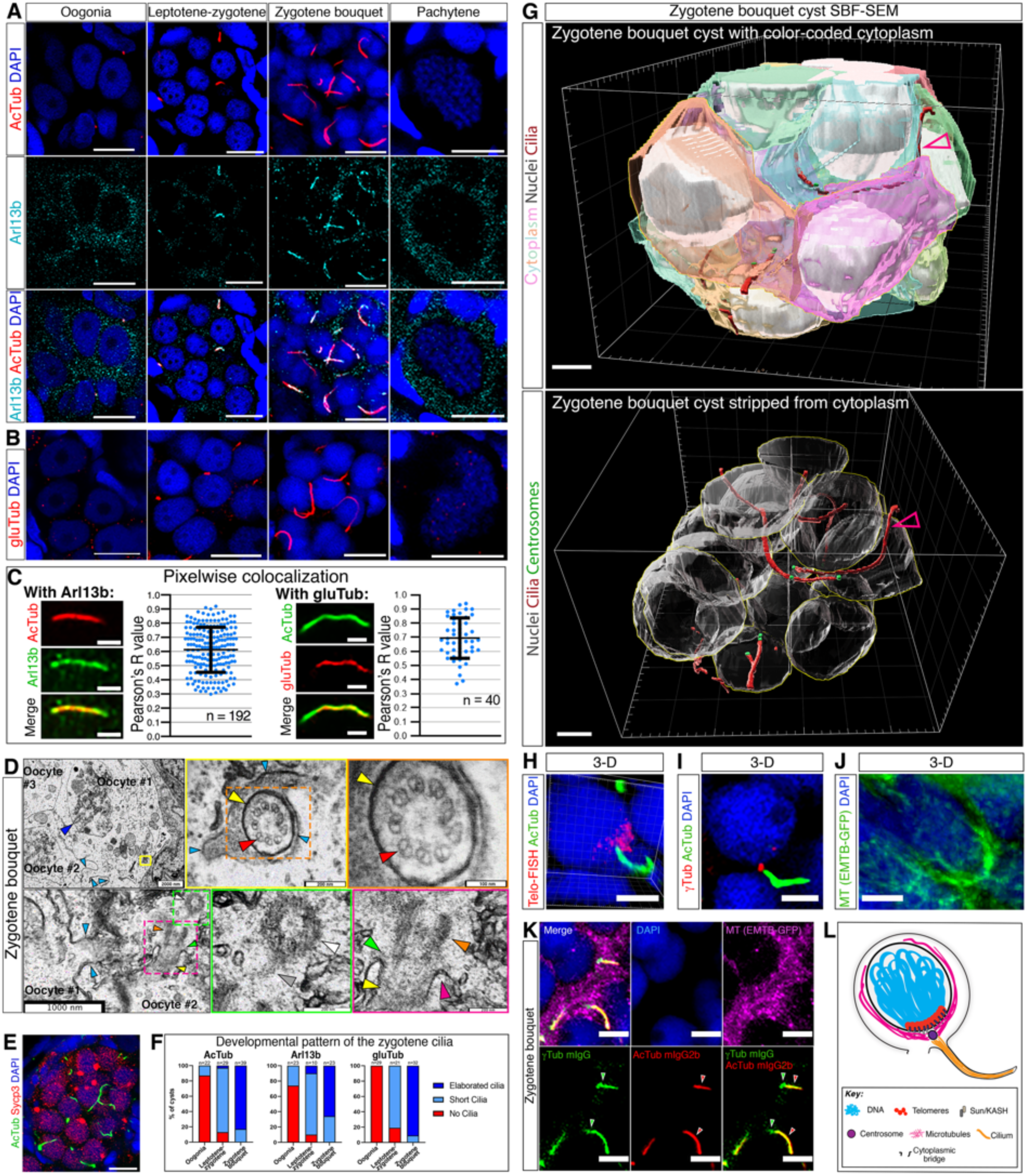
A zygotene specific oocyte cilium. **A.** AcTub and Arl13b labeling shown in oogonia, leptotene-zygotene, and zygotene bouquet cysts, as well as pachytene oocytes. n=28 cysts in 8 ovaries. Scale bar for cyst panels is 5 μm and in pachytene panels 10 μm. **B.** GluTub labeling shown as in (A). n=82 cysts in 7 ovaries. Scale bars are as in (A). **C.** Pixelwise colocalization test and representative ROI images for AcTub and Arl13b, and for AcTub and gluTub. n=number of cilia. Bars are mean ± standard deviation (SD). **D.** TEM images of cilia in leptotene-zygotene oocytes. Top left panel: leptotene-zygotene oocytes [as indicated by presumptive synapsed chromosomes that interface the NE and likely represent NE-bound telomeres (dark blue arrow)] within a germline cyst [as indicated by cytoplasmic bridges that connect oocytes (light blue arrows)]. Yellow box indicates a cross-section of a ciliary axoneme, shown in the two right panels which are magnification of the color-coded boxed regions. An axonemal structure, including ciliary membrane (yellow arrow) and MT doublets (red arrow) are detected. Light blue arrows indicate oocyte membranes. Bottom panels: ciliary structures within a zygotene oocyte, adjacent to a cytoplasmic bridge (light blue arrows). Two right panels are magnification of the color-coded boxed regions in the left panel. In the three bottom panels, arrows indicate: centriole (white), linker (grey), basal body (orange) transition zone (green), ciliary pockets (pink), axoneme (yellow). Scale bars are indicated. n=23 oocytes in 5 ovaries. **E.** AcTub and the prophase marker Sycp3 labeling shown in a zygotene cyst. n=24 cysts in 2 ovaries. Scale bar is 10 μm. **F.** The developmental distribution of the zygotene cilia as detected by AcTub (left; n>100 ovaries), Arl13b (middle; n=18 ovaries), and gluTub (right; n=11 ovaries). n=number of cysts. **G.** 3-D renders of a zygotene bouquet cyst from SBF-SEM data, showing: cytoplasm of cyst oocytes (different transparent colors), nuclei (grey), cilia (maroon; maroon-grey arrowhead), and basal body centrosomes (green). Bottom panel: same cyst render stripped from cytoplasm to show the cilia. Scale bars are 5 μm. **H.** Telomeres (Telo-FISH) and AcTub labeling in a zygotene bouquet oocyte shows that the cilium emanates from the cytoplasm apposing the telomere cluster. n=3 ovaries. A snapshot from 3-D construction in Movie S7. Scale bar is 5 μm. **I.** γTub and AcTub labeling in a zygotene bouquet oocyte shows that the cilium emanates from the centrosome, which was shown to localize apposing the telomere cluster (10). n=5 ovaries. Scale bar is 5 μm. **J.** Microtubule labeling [*Tg(βact:EMTB-3XGFP)*] in zygotene bouquet oocytes show perinuclear cables (nucleus is counterstained with DAPI). n= 7 ovaries. A snapshot from 3-D construction in Movie S8. Another example is shown in Movie S9. Scale bar is 5 μm. **K.** Co-labeling of microtubules (EMTB-GFP), γTub, AcTub, and DAPI, reveals that the zygotene cilium emanate from the centrosome, which is embedded in a perinuclear microtubule network. A merged image and all individual channels are shown. n=5 ovaries. γTub (mIgG1) and AcTub (mIgG2b) antibodies are both isoforms of mIgG. We used mIgG secondary antibody that binds both primary γTub and AcTuB antibodies, and a mIgG2b secondary antibody that specifically binds to the AcTuB antibody and not γTub antibody (arrowheads in the three bottom panels). **L.** A schematic model showing the bouquet cytoplasmic cable system that spans from the zygotene cilium through the centrosome and perinuclear microtubules to telomeres on the NE.

Interestingly, the cilia were stage-specific: They were specifically present at the leptotene-zygotene stages as confirmed by the prophase synaptonemal complex marker Sycp3 (Fig. 1E). We previously established prophase staging criteria for zebrafish oocytes based on the characteristic telomere dynamics at each stage (*4, 35*)(Fig. S1B). We characterized cellular features, including the typical oocyte size and nuclear morphology that strictly identifies each stage even without telomere labeling, and developed a reliable method for measuring oocyte size by diameter in three dimensions (*4, 35*). For example, oogonia with dispersed intranuclear telomere distribution were always ∼10 μm with non-condensed DAPI-labeled DNA and central 1-3 nucleoli, leptotene-zygotene stages with telomere radially loaded on the nuclear envelope (NE), were always 7-9 μm with central few nucleoli, and zygotene bouquet with clustered telomeres on the NE were always 10-16 μm with condensed DAPI-labeled chromosomes and a peripheral single nucleolus (*4, 35*) (Fig. S1B). Sycp3 is an established stage-specific marker during prophase, including in zebrafish (*33, 34*). We labeled ovaries with Sycp3 and measured the size of oocytes with distinct leptotene, early- and late zygotene Sycp3 patterns, confirming our criteria and sizes above (Fig. S3C). In all our analyses we measured oocyte size (*35*) and determined its prophase stage based on these criteria. Cilia were absent from most pre-meiotic oogonial stages but were detected as a few short cilia at leptotene-zygotene, then fully elaborated at zygotene bouquet and were disassembled at the pachytene stage (Fig. 1A-B, F). This developmental pattern was consistently detected using either AcTub, Arl13b or GluTub labeling (Fig. 1A-B, F). Based on these molecular, live imaging, and ultrastructural analyses, we conclude that we have been able to identify a previously unknown cilium in early prophase oocytes, specific to the leptotene-zygotene and zygotene bouquet stages.

Early prophase, including leptotene and zygotene stages, are executed while oocytes develop within the germline cyst (*4, 35, 40*). The germline cyst is a conserved cellular organization of germ cells from insects to humans (*7, 41*). The germline cyst is generated during mitotic divisions of oocyte precursor cells called oogonia, whereby cytokinesis is incomplete, and cytoplasmic bridges (CBs) persist between sister-cells and connect them (*4, 8, 35, 42*)(Fig. S1B). In the cyst, germ cells are surrounded by somatic pre-granulosa cells (*4, 6, 7*) (Fig. S1B). The germline cyst is a compact and confined cellular organization, where cytoplasmic membranes of adjacent oocytes interface tightly (*4*).

Zygotene cilia extended throughout the volume of the cyst (Fig. S3B-D). To unequivocally characterize ciliary distribution in the cyst, we performed SBF-SEM, providing ultrastructural data of cysts at EM resolution in three-dimensions (3D; Methods; Movie S3). Tracking zygotene cilia in raw SBF-SEM data (Movie S4-5) and in rendered leptotene and zygotene cysts in 3D (Fig. 1H, Movie S3, 6) confirmed that these organelles tangle tightly between oocyte cytoplasmic membranes and extend throughout the cyst (Fig. 1G). Measurements of zygotene cilia diameters showed the expected diameter of ciliary axonemes (∼300nm (*43*); Fig. S5A-B).

In the germline cyst, cytoplasmic bridges (CBs) are also based on MT cables (*4, 8*). Several lines of evidence allowed us to distinguish cilia and CBs in our analysis. ***1)*** In our TEM and SBF-SEM dataset, ciliary cytoplasm always showed axonemal MT tracks (Fig. 1D, Fig. S5A, Movie S4-5), but CB cytoplasm was indistinguishable from the remaining oocyte cytoplasm and often contained vesicles (Fig. S4A). CB membranes were continuous with the cytoplasmic membranes of the two connected oocytes and was typically ruffled (Fig. S4A). ***2)*** Measurements of CB diameters (Fig. S5A-B), were consistent with those of CBs in *Xenopus* and mice (0.5-1 μm (*44*)), but not with the ciliary diameter. ***3)*** In contrast to cilia, CBs were adjacent to the centrosomes/basal bodies (*4*), but never associated with them. ***4)*** Cilia did not extend through or co-localize with CBs as labeled by the midbody marker Cep55 (*45*) (Fig. S4B, 5C-D). Thus, zygotene cilia and CBs are distinct and separate structures.

We confirmed the organization of the zygotene cilium with the bouquet cytoplasmic machinery. Labeling bouquet clustered telomeres, perinuclear MT, centrosome, and the zygotene cilia in different combinations (Fig. 1H-K, S3E, Movie S7-9) confirmed that our findings unravel a unique cytoskeletal arrangement of a cable system that extends from the zygotene cilium, through the centrosome and MT to telomeres on the nuclear envelope, as the potential machinery for chromosomal pairing movements (Fig. 1L). Furthermore, live time-lapse imaging confirmed that the zygotene cilium was present during bouquet chromosomal rotations (Movie S10).

### The zygotene cilium is required for telomere clustering in chromosomal bouquet formation

We next aimed to investigate the function of the zygotene cilium in oogenesis. Severe disruption to differentiation of motile as well as primary cilia causes embryonic lethality in zebrafish. However, mutations in certain cilia genes allow viability to the adult stage. We analyzed multiple such *loss-of-function* ciliary mutants that in humans cause ciliopathies - genetic disorders that are caused by ciliary defects. These included Cep290 (*cep290^fh297^* (*46*)), a basal body and transition zone protein that is required for cilia formation and transition zone assembly (*47–52*), Kif7 (*kif7^i271^* (*53*)) an atypical kinesin that is required for organizing the ciliary tip compartment and Hedgehog (Hh) signaling (*54–56*), Cc2d2a (*cc2d2a^w38^*), a transition zone protein and part of a multiprotein complex that acts as a gatekeeper to the ciliary compartment (*57, 58*).

We analyzed zygotene cilia in juvenile ovaries of *cep290^fh297^, kif7^i27^*, and *cc2d2a^w38^* mutants (hereafter referred to as *cep290^-/-^*, *kif7^-/-^,* and *cc2d2a^-/-^*, respectively; Fig. 2A-D). Based on wt ciliary length measurements in 3D (Fig. 2A, Fig. S6, Methods), we defined “normal cilia” (4-8.5 μm), “short cilia” (<4 μm) and absent cilia (N.D). Individual ciliary length (Fig. 2A), average ciliary length per cyst (Fig. 2B), distribution of scored cysts (Fig. 2C), and scoring of entire ovaries (Fig. 2D) established the shortening and loss of zygotene cilia in *cep290^-/-^* ovaries. Attempting to abolish cilia altogether, we generated *cep290; kif7* double mutants. *Kif7^-/-^* ovaries showed only mild partial ciliary shortening phenotypes (Fig. 2A-D, Fig. S6). We hypothesized that loss of Cep290 together with that of Kif7 would synergistically cause more severe ciliary defects. Indeed, in *cep290^-/-^; kif7^+/-^* and *cep290^-/-^; kif7^-/-^* ovaries, cilia were mostly or completely abolished (Fig. 2A-D, Fig. S6). In addition, *cc2d2a^-/-^* ovaries showed similar ciliary shortening and loss.

**Figure 2.**
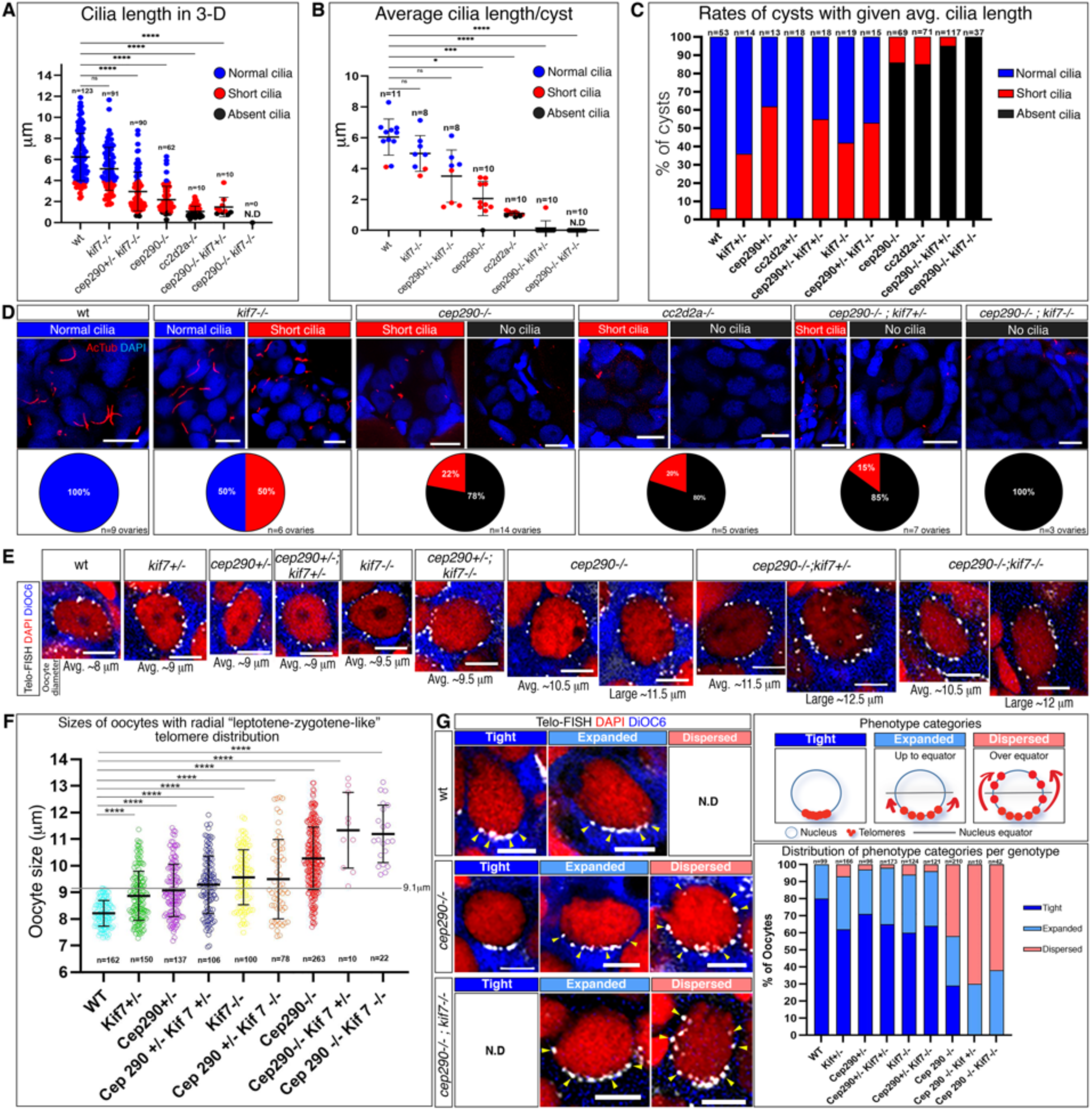
The zygotene cilium is required for bouquet formation. **A.** 3-D measurements of ciliary lengths in juvenile ovaries. n=number of cilia. N.D, not detected. Bars are mean ± SD. **B.** Average ciliary length per cyst. n=number of cysts. Bars are mean ± SD. **C.** Distribution of cysts with normal and short ciliary length averages, as well as with absent cilia. *Chi-square* test *p*-values are <0.0001 for all groups. **D.** Representative images (AcTub labeling) of cysts in juvenile ovaries with indicated ciliary length categories as in B-D per genotype. Percentage of ovaries with indicated predominant categories are shown below. n=number of ovaries. Scale bars are 10 μm. **E.** Representative images of “leptotene-zygotene-like” oocytes of average sizes per genotype. Large “leptotene-zygotene-like” *cep290^-/-^*, *cep290^-/-^;kif7^+/-^,* and *cep290^-/-^;kif7^-/-^* oocytes are shown. Scale bars are 5 μm. **F.** “Leptotene-zygotene-like” oocyte sizes per genotype. n=number of oocytes. Bars are mean ± SD. 9.1 μm represents the largest wt leptotene oocytes. **G.** Left panel: representative images of mid-bouquet stage oocytes with Tight, Expanded, and Dispersed telomeres (categories shown in top right panel). N.D=non-detected. Scale bars are 5 μm. Right bottom panel: the percentage of each category in all genotypes. n=number of oocytes. *Chi-square* test *p*-values n.s between wt and *cep290^+/-^,* or *cep290^+/-^;kif7^+/-^*, are <0.05 or <0.01 between wt and *kif7^+/-^, kif7^-/-^, cep290^+/-^;kif7^-/-^,* and <0.0001 between wt and *cep290^-/-^* or *cep290^-/-^;kif7^-/-^*.

Having established the loss of zygotene cilia in these ciliary mutants, we analyzed bouquet formation in developing ovaries from juvenile wt and mutant fish. We first noticed that oocytes with clear telomere clusters were fewer in *cep290^-/-^* ovaries, suggesting that prophase progression is defective. We therefore asked: 1) if oocytes at leptotene-zygotene stages are delayed in bouquet formation, 2) if oocytes at bouquet stages formed proper telomere clusters, and 3) if late bouquet oocytes eventually formed telomere clusters to distinguish between a delay versus failure in bouquet formation.

First, to address a delay in leptotene-zygotene stages, we asked whether some of the oocytes that showed radial telomere distribution on the NE, that is typical of the leptotene-zygotene stages, were abnormally larger as an indication for prophase progression delay. Wt leptotene-zygotene oocytes are 7-9 μm in diameter and zygotene bouquet oocytes are 10-16 μm in diameter (Fig. S1B, Fig. 2E-F) (*4, 35*). We measured the diameter of all oocytes with radial telomeres on the NE, based on their cytoplasmic staining by DiOC6 (*4, 35*).

In wt ovaries, these oocytes were 8.21 ±0.48 μm (Fig. 2E-F). In mutants, we identified oversized “leptotene-zygotene-like” oocytes, defined as oocytes with leptotene-zygotene-like radial telomere distribution, which were larger than the largest detected wt oocyte at these stages (>9.1 μm; Fig. 2E-F, S7B). In single and double heterozygous, *kif7^-/-^,* and *cep290^+/-^; kif7^-/-^* ovaries, ∼40-65% of oocytes with radial telomeres were mildly larger leptotene-zygotene-like oocytes (8.88 ±0.92 μm to 9.49 ±1.49 μm; Fig. 2E-F, Fig. S7A). In *cep290^-/-^* ovaries, 85% were much larger leptotene-zygotene-like oocytes (10.28 ±1.18 μm. ; Fig. 2E-F, Fig. S7A). In the *cep290^-/-^; kif7^-/-^* and *cep290^-/-^; kif7^+/-^* double mutants most ovaries were severely affected and contained low numbers of oocytes for analysis, and many ovaries of this genotype converted to testes. In zebrafish, ovary conversion to testis is indicative of severe oocyte damage and loss (*59–61*), which we address and discuss below and in Fig. 5B, S12. Nevertheless, we did capture some ovaries for analysis. In *cep290^-/-^; kif7^-/-^* and *cep290^-/-^; kif7^+/-^*ovaries, 100% of oocytes with radial telomeres were very large leptotene-zygotene-like oocytes (11.33 ±1.42 μm and 11.2 ±1.08 μm, respectively; Fig. 2E-F, Fig. S7A). Similar large leptotene-zygotene-like oocytes were detected in *cc2d2a^-/-^* ovaries (10.6 ±1.00 μm; Fig. S7B). This suggests that oocytes continued to grow in size, but bouquet formation was at least delayed.

In mice, activation of the zygotene meiotic checkpoint results in zygotene arrest in males, but oocytes with activated checkpoint continue to grow in size and are cleared later by apoptosis (*12, 62*). Consistent with this, in zebrafish mutant for *spo11,* encoding a DNA topoisomerase essential for meiotic double-strand break induction, males fail to produce sperm, while mutant females generate defective oocytes (*33*). The delay in progression from leptotene-zygotene to zygotene bouquet stages that we observed suggests that similar dynamics are likely with ciliary loss in zebrafish.

Second, we tested whether bouquet stage oocytes properly formed telomere clusters, and the bouquet configuration. We analyzed oocytes in sizes that were typical to mid-bouquet stages in the wt. We reasoned that the telomere cluster may be less compact during its early formation (oocyte sizes of 10-11.5 μm in diameter) and its later dissociation (oocyte sizes of 13-16 μm in diameter). We therefore focused on mid-bouquet stages (oocyte sizes of 11.5-13 μm in diameter), that showed clear clusters. We measured all oocyte sizes blind to the telomere labeling channel based on their cytoplasmic DiOC6 labeling, selected the ones within the 11.5-13 μm size range, and then scored their telomere cluster (Methods).

Telomere distribution in wt mid-bouquet stages (Fig. 2G, Methods) showed either tight telomere cluster or a slightly expanded cluster (“Tight” and “Expanded” in Fig. 2G). Ovaries of heterozygous, *kif7^-/-^,* and *cep290^+/-^; kif7^-/-^* fish were mostly similar to wt (Fig. 2G). However, in *cep290^-/-^* ovaries, in addition to the wt categories, we detected an abnormal “dispersed” category (Fig. 2G), where telomeres were largely expanded beyond the nuclear equator (Fig. 2G). Moreover, in double mutant *cep290^-/-^; kif7*^+/-^ and *cep290^-/-^; kif7*^-/-^ ovaries, we only detected “expanded” and “dispersed” category oocytes (Fig. 2G). Thus, telomere clustering is defective in bouquet stage mutant oocytes. Mid-bouquet stage oocytes in *cc2d2a^-/-^* ovaries exhibited consistent telomere clustering defects (Fig. S7C).

Thirdly, we examined whether in mutant ovaries, oocytes at late bouquet stages ultimately formed proper telomere clustering, which would suggest that clustering was delayed. We repeated the above analysis on oocytes at sizes of 13-15 μm in diameter, wherein the wt telomere clusters are “expanded” and “dispersed” (Fig. S7D-E) towards bouquet dissociation (*4, 35*). Heterozygous *cep290* and *kif7*, as well as *cep290^+/-^;kif7^+/-^* and *kif7^-/-^* ovaries, were similar to the wt (Fig. S7D). In *cep290^-/-^* ovaries, telomeres were “expanded” and most were “dispersed”, and in *cep290^-/-^;kif7^+/-^* and *cep290^-/-^;kif7^-/-^* ovaries all oocytes were “dispersed” (Fig. S7D-E). “Tight” telomere clusters were not detected at these stages, ruling out delayed clustering. Tight telomere clusters were similarly not detected in late bouquet stage oocytes in *cc2d2a^-/-^* ovaries as well (Fig. S7E). Thus, with loss of cilia, telomeres do not cluster properly, remain distributed on the nuclear envelope, and fail to form the bouquet.

These ciliary loss-associated defects are similar to bouquet disruption by dynein inhibition. Dynein carries Sun/KASH bound telomere complexes in mice (and the bouquet analogous pairing centers in *C. elegans*), sliding them on MTs and is essential for bouquet formation (*22–29*). We treated ovaries with the specific dynein inhibitor ciliobrevin and performed the same bouquet analysis as above (Fig. S8A). In contrast with control DMSO-treated ovaries, in ciliobrevin treated ovaries, dynein inhibition resulted in a dose dependent increase in zygotene bouquet oocytes (11.5- 13 μm in diameter) with “expanded” and “dispersed” telomeres and decrease in oocytes with “tight” telomere clusters (Fig. S8A). Thus, the loss of the zygotene cilia phenocopied dynein inhibition in bouquet formation.

We determined whether these bouquet phenotypes arose directly due to ciliary defects. Cep290 subcellular localizations include the ciliary basal body and transition zone, as well as the centrosome and centriolar satellites (*48, 52, 63-67*). Cep290 is essential for cilia biogenesis (*47–52*), but whether it is required for centrosome MTOC functions in oocytes is unknown. In *cep290^-/-^* oogonia and early prophase oocytes, γTub recruitment to the centrosome, which is the final step in MTOC maturation (*68, 69*), was intact even in the most severe cases where cilia were completely absent (Fig. S9B). In line with this, MT perinuclear organization was not disrupted (Fig. S8C). This demonstrates that the centrosome MTOC functions are unaltered, confirming that the observed bouquet defects arose directly from the loss of cilia.

### Ciliary anchoring of the bouquet centrosome is required for telomere clustering and proper synaptonemal complex formation

We hypothesized that mechanistically, the zygotene cilium could generate and transmit forces for telomere rotations or provide counterbalance for their movements by anchoring the bouquet centrosome. Since force generation by the cilium would require dynein and kinesin motor activity along the ciliary axoneme, and could involve cilia motility, and we addressed the motility of the zygotene cilium. Four lines of evidence suggest that the zygotene cilium in not a motile cilium: 1) Motile cilia axoneme is typically comprised of 9 MT doublets with dynein arms and a central pair of MT singlets in a “9+2” organization, and non-motile cilia usually exhibit a “9+0” MT doublet organization without a central pair and dynein arms (*70, 71*). Ultrastructure analysis of the zygotene cilium showed a 9+0 organization without obvious dynein arms (Fig. 1D). ***2)*** Live time-lapse imaging of cilia during chromosomal rotations did not reveal clear beating movement, which is a characteristic of motile cilia (Movie S10-11). ***3)*** Transgenic reporter lines of motile ciliary master transcriptional regulators of the Foxj1 family, [*Tg(foxj1a:GFP)* and *Tg (foxj1b:GFP)*] (*72, 73*), showed no or very weak expression in prophase oocytes within germline cysts where the zygotene cilium forms (Fig. S9). ***4)*** Ccdc103 is a dynein assembly factor specific to motile cilia and required for their motility (*74*). To test if cilia motility was essential for fertility, we were able to rescue the embryonic lethal *ccdc103* mutant to adult stages by wt sense mRNA injection. Rescued *ccdc103^-/-^* females were fertile (5/6 females mated spontaneously and each produced 50-300 eggs with 50-100% fertilization rate), leading us to conclude that motility is not essential for fertility, and that the zygotene cilium is not likely a motile cilium.

We crossed *Tg(h2a:H2A-GFP)* fish to *cep290* mutants and performed live time-lapse imaging of chromosomal rotations in *cep290^-/-^* ovaries (Fig. 3A, Movie S12). Chromosomal rotation in *wt* H2A-GFP siblings (Fig. 3A, Movie S12) were consistent with data shown in Fig. S2, showing similar gross track velocities (Fig. 3A, Movie S12). In *cep290^-/-^* oocytes, chromosomal rotations were still in place and we only detected a very small decrease in track velocities (from 3.56 ±0.67 in wt to 3.15 ±0.8; Fig. 3A, Movie S12). 20% of *cep290^-/-^* oocytes, including 13.6% outliers, were plotted lower than the lowest wt oocyte (Fig. 3A, Movie S12). However, these *cep290^-/-^* oocytes still exhibited slower but clear rotations (Fig. 3A left bottom panels, Movie S12). This analysis was limited to gross chromosomal rotations and could not resolve fine quantitative details of rotating individual telomeres. However, the persistence of gross chromosomal rotations in *cep290^-/-^* oocytes allow us to conclude that the zygotene cilium is not an essential generator of chromosomal rotation forces.

**Figure 3.**
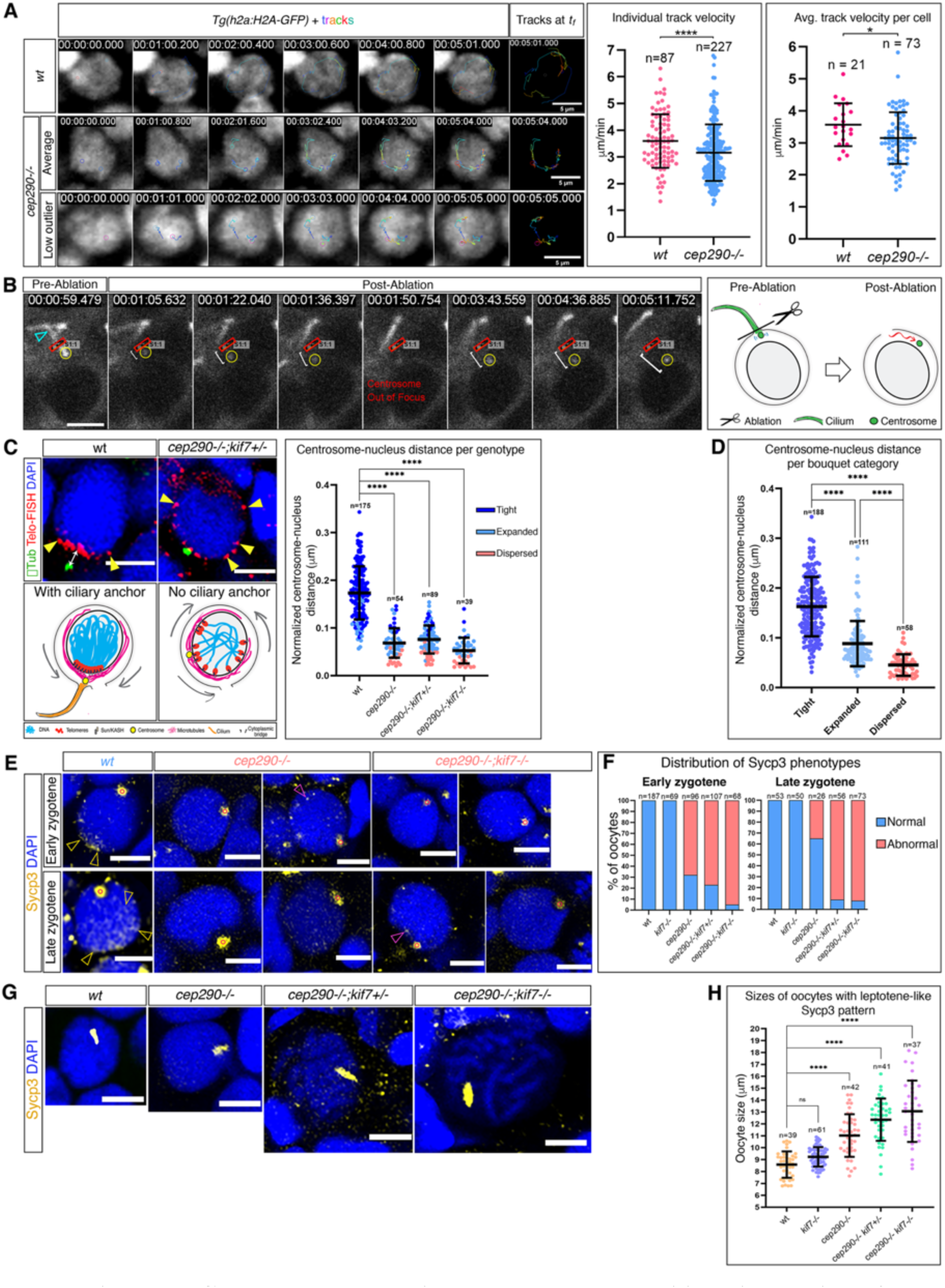
Centrosome anchoring by the zygotene cilium is required for telomere clustering in bouquet formation and for proper synaptonemal complex formation. **A**. Live time-lapse imaging of chromosomal bouquet rotations in *wt Tg(h2a:h2a-gfp);cep290^+/+^* (top) and sibling *Tg(h2a:h2a-gfp);cep290^-/-^* (middle and bottom) ovaries. Chromosomal tracks and their sum at *T_f_* are shown for each cell. Plots on the right are individual track velocities (n=number of tracks) and average track velocity per cell (n=number of cells) for each genotype. Two *cep290^-/-^* oocytes are shown: the middle panel shows a representative oocyte with mean values of average track velocities, and the bottom panel show an oocyte with low outlier values of average track velocities below the SD. Images are snapshots from Movies S12. Scale bars are 5 μm. **B.** Laser ablation of the zygotene cilium. The zygotene cilium that is targeted for ablation (teal arrowhead) is shown in the pre-ablation panel and the ablation region at its base (row labeled S1:1) is indicated throughout the time-lapse. Note that the ablated ciliary associated centrosome (yellow circle) dislocates upon ablation in both XY and Z (indicated as centrosome out of focus) axes. Time is indicated in mili-seconds. Scale bar is 5 μm. n=17 cilia (oocytes), from 11 ovaries. Images are snapshots from Movie S13 and another example is shown in Movie S14. Right panel shows a cartoon scheme of the experiment. **C.** Representative images of mid-bouquet stage oocytes from wt and *cep290^-/-^; kif7^+/-^* ovaries co-labeled for the centrosome (γTub) and telomeres (Telo-FISH). The distance between the centrosome and the nucleus (white arrow) in the wt is significantly decreased in mutants oocytes with defected telomere clustering (yellow arrowheads), as schematized in the cartoons in the bottom panels. The normalized distances for oocytes from wt, *cep290^-/^, cep290^-/-^; kif7^+/-^,* and *cep290^-/-^; kif7^-/-^* ovaries are plotted in the right panel. Note that bouquet phenotype categories are color-coded showing an almost complete elimination of the centrosome-nucleus distances in mutant dispersed telomere oocytes. Scale bar is 5 μm. n=number of oocytes. **D.** The normalized centrosome-nucleus distance in oocytes pooled from all genotypes in C, categorized based on their bouquet phenotype. n=number of oocytes. **E.** Representative images of Sycp3 localization patterns in early (top panels) and late (bottom panels) zygotene stages. Note normal Sycp3 patterns in wt oocytes (left panels, yellow arrowheads), and their absence in mutant oocytes, where only few scattered foci are detected (right four panels, magenta arrowheads). Empty red circle denote the nucleoli in which Sycp3 signal accumulate in all wt and mutant oocytes. Scale bars are 5 μm. **F.** The distribution of normal and abnormal Sycp3 oocytes in E is plotted for wt, *kif7^-/-^, cep290^-/^, cep290^-/-^; kif7^+/-^,* and *cep290^-/-^; kif7^-/-^* ovaries in early and late zygotene stages. n=number of oocytes. *Chi-square* test *p*-values for both plots are <0.0001 between wt and all genotypes, except for wt and *kif7^-/-^* is n.s. G. Representative images of “leptotene-like” oocytes from wt, *cep290^-/-^*, *cep290^-/-^;kif7^+/-^,* and *cep290^-/-^;kif7^-/-^* ovaries, shown in scale. Scale bars are 5 μm. **H.** “Leptotene -like” oocyte sizes per genotype. n=number of oocytes. Bars are mean ± SD.

We next addressed the potential ciliary anchoring of the bouquet centrosome. The observations that the centrosome is stationary during chromosomal rotations in zebrafish (Movie S2) and mice (*30*), while telomeres are being pulled on MT cables that are associated to it, suggests that the centrosome is under strain. It is plausible that the centrosome-end of the cables must be affixed in order for the telomeres to be pulled and clustered towards the centrosome, and not vice versa. If this is the case, without such counterbalance, telomeres will fail to cluster and form the bouquet. We hypothesized that the zygotene cilium could function as a physical anchor, that in turn would be required for counterbalancing telomere rotation forces, and for telomere pulling and clustering.

To experimentally eliminate potential ciliary anchoring, we performed laser excision experiments (see Methods) in ovaries of double transgenic fish fluorescently co-labeled for cilia and centrosomes [*Tg(βact:Cetn2-GFP); Tg(βact:Arl13b-GFP)*]; Fig. 3B). In consideration that the centrosome is stationary and likely under strain during telomere rotation, we reasoned that if the zygotene cilium is required to anchor the centrosome, then ciliary excision would result in dislocation of the centrosome. We therefore excised the ciliary base in zygotene bouquet stage oocytes (as determined by our staging criteria above, as well as by the presence of clear zygotene cilia), while tracing the centrosome by live time-lapse imaging. We carefully adjusted the excision laser power and ROI and all microscopy settings to conditions that consistently and precisely excise the ciliary base (Methods). These analyses showed that upon ciliary excision, the associated centrosome immediately dislocated away (Fig 3B yellow circle, Movie S13-14 red circles). In contrast, other centrosomes in the same cyst and frame that were associated with non-ablated cilia remained relatively stationary (Movie S13-14 green circles).

In control experiments, we showed that a significantly higher laser power and/or non-specific excisions were required for inducing cellular damage. Excising the cilia or regions in the cytoplasmic membrane with almost two-fold greater laser power in otherwise consistent experimental settings (Methods), immediately resulted in clear and stereotypic cell death (Movie S15-16). This was never observed in the above experimental groups, demonstrating the precise ciliary excision achieved in these experiments. Thus, when disconnected from the cilium, the centrosome does not remain stationary but moves around, likely by the rotational forces of hauled telomers that act on its associated microtubules.

We next addressed genetically the bouquet centrosome dynamics upon ciliary loss. We found that in the conditions were telomeres were dispersed upon ciliary loss in *cep290^-/-^, cep290^-/-^;kif7^+/-^,* and *cep290^-/-^;kif7^-/-^*, the centrosome was much more juxtaposed to the nucleus, compared to wt (Fig. 3C). We labeled ovaries for the centrosome (γTub) and telomeres (Telo-FISH) and measured the distance between the centrosome and the nucleus, normalized to the oocyte size in mid-zygotene bouquet stage oocytes (Methods; arrow in Fig. 3C). These normalized distances were significantly decreased from 0.174 ±0.056 in wt to 0.068 ±0.03 in *cep290^-/-^*, 0.076 ±0.03 in *cep290^-/-^;kif7^+/-^*, and 0.052 ±0.027 in *cep290^-/-^;kif7^-/-^* oocytes (Fig. 3C). In parallel, we independently scored telomere clustering in these oocytes as in Fig. 2G. The shortest centrosome-nucleus distances were measured in oocytes with dispersed telomeres (telomere clustering categories are color coded in Fig. 3C). These results demonstrate that in the absence of the ciliary anchor, telomeres are not normally pulled towards the centrosome; by contrast, the centrosome is pulled towards the nucleus, reminiscent of centrosome dislocation upon lase excision of the ciliary base.

Further, we reciprocally tested the centrosome-nucleus distances per telomere clustering category in pooled data from all genotypes above (Fig.3D). We found that the centrosome-nucleus distance associated with telomere clustering capability: sufficient distance (0.163 ±0.06) in “Tight” correlated with successful clustering, while oocytes with “dispersed” failed clustering always showed the shortest distances (0.045 ±0.021; Fig.3D). This provides further credence to the view that a counterbalance between the centrosome and rotating telomeres is an inherently required feature of bouquet formation. Together, these results suggest that the zygotene cilium is required for anchoring the centrosome as a counterbalance for telomere rotations, and that when lost, the centrosome is instead pulled towards the nucleus instead, telomere pooling is less efficient, and the bouquet formation fails.

We next finally tested whether bouquet failure affected the formation of synaptonemal complexes between pairing chromosomes. It has been shown in zebrafish that the bouquet telomere cluster is a hub for chromosomal pairing (*33*). Double-strand breaks cluster in sub- telomeric regions, and the loading of synaptonemal complex protein 3 (Sycp3) is initially seeded in these regions specifically at bouquet stages (*33, 61*). Sycp3 then spreads from this position along the axes of pairing chromosomes (*33, 61*). These findings suggest that telomere clustering in the bouquet configuration could be functionally required for synaptonemal complex formation. We examined the consequences of defected telomere clustering upon ciliary loss on Sycp3 patterns in wt and mutant ovaries (Fig. 3E-F).

We analyzed Sycp3 localization patterns in wholemount wt and mutant ovaries (Methods) in early zygotene bouquet (oocyte size 10-12 μm) and late zygotene bouquet (oocyte size 12-14 μm) stages as defined in Fig. S1B-C. Consistent with Fig. S1C, all wt and *kif7^-/-^* zygotene oocytes showed normal Sycp3 patterns, including initial polarized loading of Sycp3 at early zygotene bouquet stages (*33, 61*), and the beginning of its extension along chromosome axes at late zygotene bouquet stages (*33, 61*) (Fig. 3E, yellow arrowheads). In contrast, in most zygotene stage oocytes of *cep290^-/-^*, *cep290^-/-^;kif7^+/-^*, and *cep290^-/-^;kif7^-/-^* ovaries, these normal Sycp3 patterns were completely lost with only few scattered foci remaining in some oocytes (Fig. 3E-F, magenta arrowheads). All zygotene oocytes from all genotypes showed accumulation of Sycp3 signal in oocyte nucleoli (Fig. 3E, red empty circles). These results are consistent with the corresponding bouquet defects in these mutants (Fig. 2G).

Further, we noticed in mutants many oocytes with an Sycp3 oval patch pattern, which is a characteristic of the leptotene stage (*34*)(Fig. S1C), that were larger than the typical wt leptotene size (Fig. S1C). Measuring the diameter of all oocytes exhibiting a leptotene-like Sycp3 pattern, confirmed their typical sizes in wt, and similarly in *kif7^-/-^* ovaries, and that most of these oocytes in *cep290^-/-^, cep290^-/-^;kif7^+/-^,* and *cep290^-/-^;kif7^-/-^* ovaries were oversized (Fig. 3G-H). In line with the detected oversized leptotene-zygotene oocytes by our Telo-FISH analyses (Fig. 2E-F), these results confirm defects in prophase progression, where defected oocytes continue to grow while delayed in prophase, as discussed above.

Altogether, these findings demonstrate that upon ciliary loss, failed telomere clustering resulted in failed Sycp3 loading to- and along chromosome axes. Thus, the zygotene cilium is required for proper formation of synaptonemal complexes, at least partly via bouquet centrosome anchoring.

### Loss of the zygotene cilium perturbs germline cyst morphogenesis

We further identified a second phenotype in the ciliary mutant ovaries, where the morphology of germline cysts was abnormal. Wt cysts are compact, with no gaps between cytoplasmic membranes of adjacent oocytes (*4*), as detected by DiOC6 cytoplasmic staining (“Normal” in Fig. 4A). However, we observed an abnormal cyst integrity phenotype in *cep290^-/-^*, and *cep290^-/-^; kif7*^+/-^ ovaries (Fig. 4A, D), and categorized them based on their severity: “loose cysts” showed abnormal gaps between oocytes, while more severe “disintegrated cysts” showed larger gaps and exposed elongated cytoplasmic bridges (CBs; Methods section). In the compact wt cysts, CBs cannot be resolved without a specific marker, but CBs were clear in the increased spaces between oocytes in disintegrated cysts of the mutants. To quantify cyst disintegration, we measured the distance between the centers of all neighboring nuclei in 3D in wt, *cep290^-/-^,* and *cep290^-/-^; kif7*^+/-^ cysts (Fig. 4B, Movie S17), normalized to the average oocyte diameter per cyst (Fig. 4B, right panel). These measurements showed that the normalized distances between neighboring nuclei in *cep290^-/-^,* and *cep290^-/-^; kif7*^+/-^ cysts were significantly increased compared to the wt (Fig. 4B). A similar cyst integrity phenotype with greater normalized distances between cyst nuclei was observed in *cc2d2a* mutant ovaries (Fig. S10A-B).

**Figure 4.**
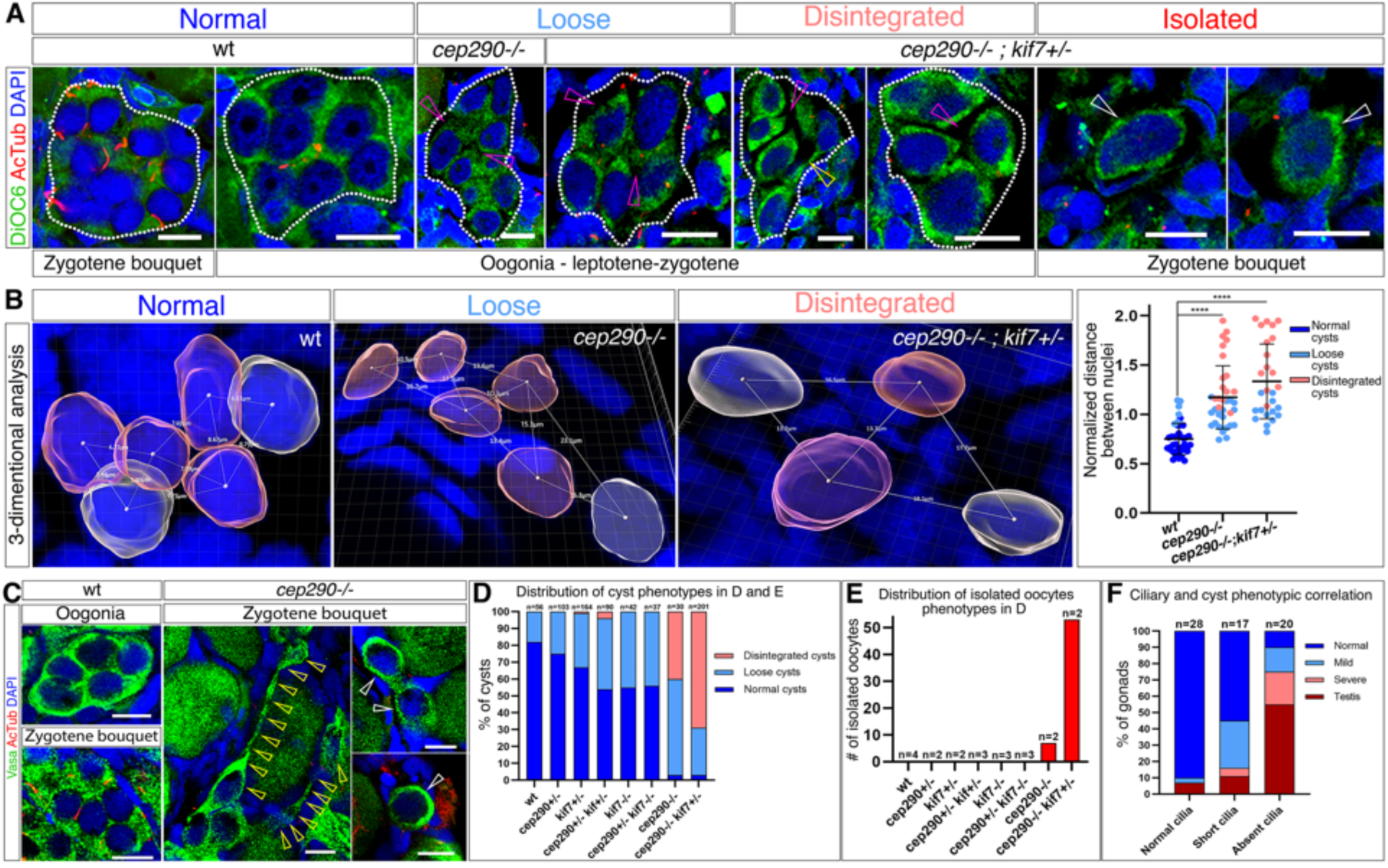
The zygotene cilium is required for germline cyst morphogenesis. **A.** Representative images of cyst (white outlines) phenotype categories in ovaries labeled for the cytoplasmic marker DiOC6 and AcTub, showing gaps between oocytes (magenta arrows), elongated cytoplasmic bridges (yellow arrows), and isolated oocytes (white arrows). Scale bars are 10 μm. The percentage of cyst phenotype categories and number of isolated oocytes in all genotypes from this experiment is shown in **D** (n=number of cysts) and **E** (n=number of ovaries), respectively. *Chi-square* test *p*-values in D, are n.s between wt and *cep290^+/-^,* and <0.05 between wt and *kif7^+/-^,* <0.001 between wt and *cep290^+/-^;kif7^+/-^, kif7^-/-^,* and *cep290^+/-^;kif7^-/-^,* and <0.0001 between wt and *cep290^-/-^* and *cep290^-/-^;kif7^+/-^.* **B.** Representative images of 3-D cyst morphology analysis per category, showing the distances between neighboring nuclei. Normalized distances are plotted in the right panel. n=5-7 cysts from 2-3 ovaries per genotype. Bars are mean ± SD. **C.** Vasa and AcTub labeling shows normal cysts in wt ovaries, and disintegrated cysts (elongated CBs, yellow arrows), as well as isolated zygotene oocytes (white arrows) in *cep290^-/-^* ovaries. Phenotype distribution in all genotypes is plotted in Fig. S11. Scale bars are 10 μm. **F.** Severity of cyst phenotypes and gonad conversions plotted for three categories of ciliary defects pooled from all genotypes. n=number of gonads. *Chi-square* test *p*-values are <0.0001 for all groups.

Strikingly, we observed a third and most severe category of “isolated oocytes” in *cep290^-/-^* and *cep290^-/-^; kif7*^+/-^ ovaries (Fig. 4A, E). Isolated oocytes are zygotene bouquet stage oocytes that are never seen outside the germline cyst in wt ovaries, but were found individually scattered in *cep290^-/-^* and *cep290^-/-^;kif7*^+/-^ ovaries (Fig. 4A), further indicating their disintegration from cysts. In our telomere labeling experiments (Telo-FISH), we were able to identify oocytes at zygotene bouquet stage with dispersed telomeres that were isolated from cysts (Fig. S11A), confirming that oocytes in the isolated category are abnormal zygotene bouquet oocytes. Overall, in contrast to wt, heterozygous, and *kif7^-/-^* ovaries, ovaries of *cep290^-/-^*, *cc2d2a^-/-^*, and *cep290^-/-^; kif7*^+/-^ fish barely contained normal cysts, but instead harbored mostly disintegrated cysts and isolated oocytes (Fig. 4D, S10A). *cep290^-/-^; kif7*^-/-^ gonads converted to testes in these experiments, preventing oocyte analysis, but indicating oocyte loss due to severe defects (see discussion below and Fig. 5, S12).

**Figure 5.**
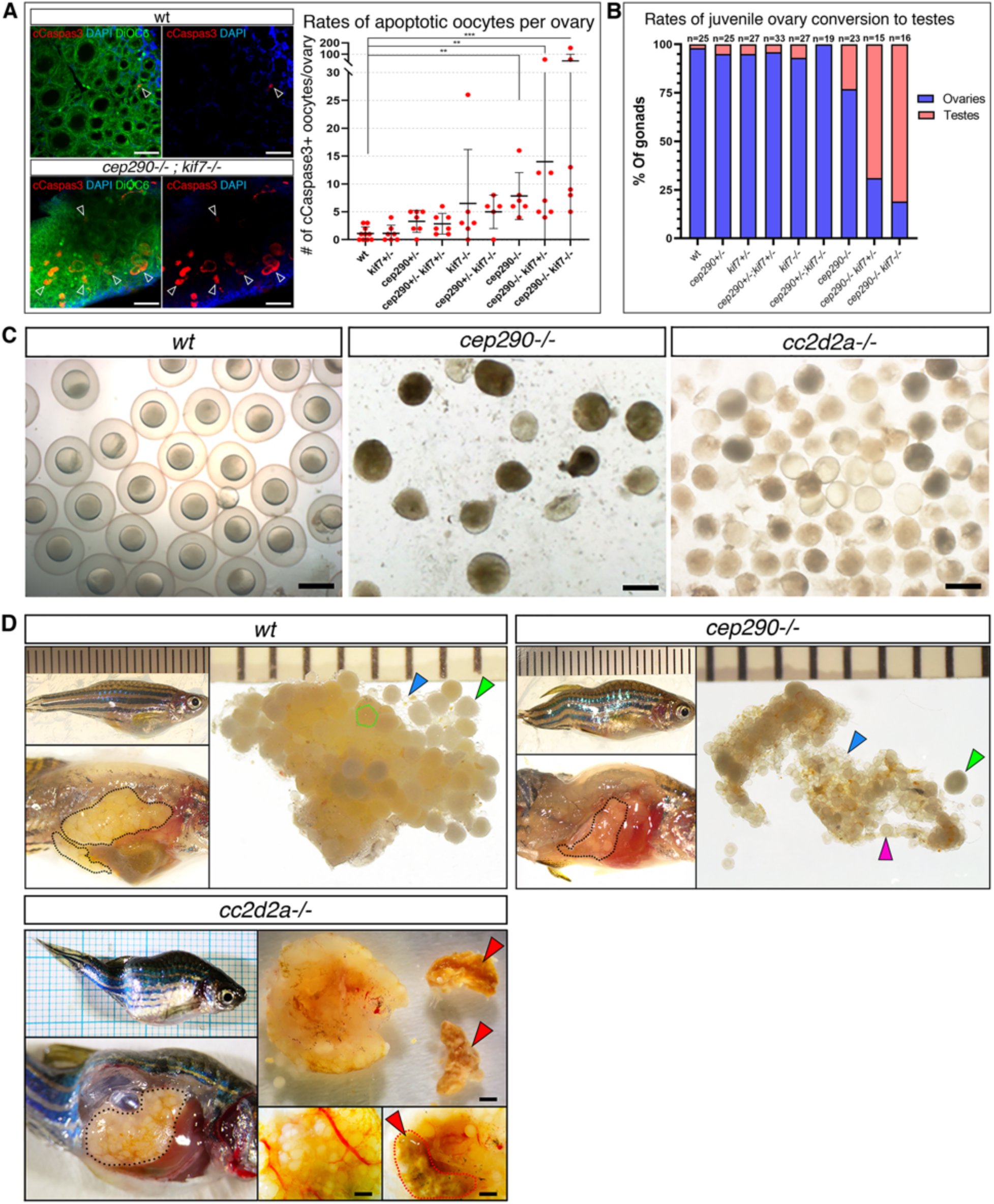
Ciliary mutants show defective ovarian development and fertility. **A.** cCaspase3 apoptosis labeling (white arrowheads) in juvenile ovaries. Left panels: representative images of wt (top) and *cep290^-/-^; kif7^-/-^* (bottom) ovaries. DiOC6 is a cytoplasmic marker. Scale bars are 50 μm. Right panel: number of cCaspase3-positive oocytes per gonad for all genotypes. Each dot represents a gonad, n=5-10 gonads per genotype. Bars are mean ± SD. **B.** Rates of juvenile gonad conversion per genotype. n=number of gonads. Images of normal and converted gonads are shown in Fig. S12. *Chi-square* test *p*-values are n.s between wt, *cep290^+/-^,* all *kif7* heterozygous combinations, and *kif7^-/-^,* and <0.0001 between wt and *cep290^-/-^, cep290^-/-^;kif7^+/-^,* and *cep290^-/-^;kif7^-/-^*. **C.** Representative images of eggs obtained by squeezing for IVF experiments from *wt*, *cep290^-/-^*, and *cc2d2a^-/-^* females. Scale bar=1mm. **D.** *Wt*, *cep290^-/-^*, and *cc2d2a^-/-^* adult females and ovaries showing: ovaries within the peritoneal cavity (black outline), st.III premature oocytes (green arrows and outlines), young transparent st.I oocytes (blue arrows), st.II oocytes (pink arrows), and degenerated tissue masses (red arrow and outline). Ruler marks and scale bars are 1 mm. Mutant females exhibit scoliosis, a typical ciliary phenotype.

Labeling ovaries with the germ cell-specific marker Vasa confirmed the detection of disintegrated cysts, where long cytoplasmic bridges extended between distant oocytes (Fig. 4C, Fig. S11B-C). These oocytes exhibited oval and irregular cellular morphology that coincided with their likely abnormal separation (Fig. 4C). Vasa labeling also detected isolated zygotene oocytes in doublets and singlets (Fig. 4C, S11B-C). Moreover, later pachytene and diplotene stage oocytes exhibited irregular mesenchymal-like cellular morphology in *cep290^-/-^* ovaries, instead of their smooth spherical morphology in the wt (Fig. S11C). Disintegration of cysts in our Vasa experiments was mostly apparent in *cep290^-/-^* mutants, while *cep290^-/-^; kif7*^+/-^ and *cep290^-/-^; kif7*^-/-^ were converted to testes (see discussion below) (Fig. S11B). These results show that the zygotene cilium is required for cyst integrity and morphogenesis.

In all the experiments above, we co-labeled cilia by AcTub staining and simultaneously analyzed cyst integrity and ciliary phenotypes. We performed correlation analysis between these phenotypes by categorizing gonads from all genotypes (n=43 gonads, 10-60 cysts were analyzed per gonad, see Methods) based on their ciliary phenotype (normal, short, absent, as in Fig. 2A-D), and separately their cyst integrity phenotypes (normal, mild, severe in Fig. 4D). Cross-referencing these data revealed a strong correlation between the severity of ciliary loss and severity of cyst phenotypes (Fig. 4F, S10C), and is in line with our findings that ciliary defects directly underlie bouquet phenotypes (Fig. S8B). Complete ciliary loss predicts conversion to testes, which represents oocyte loss in zebrafish (see below) and the most severe phenotype, and was observed in ∼60% of gonads (Fig. 4F). We found similar correlation between ciliary loss and cyst phenotypes in *cc2d2a* ovaries as well (Fig. S10C).

### Loss of the zygotene cilium is detrimental to oogenesis and fertility

Substantial oocyte loss by apoptosis during ovarian development in zebrafish often results in gonad conversion to testis (*59–61*). In the zebrafish, gonads initially develop as ovaries that execute normal oogenesis (*4, 35*). Sex determination then dictates continuation of ovarian development or their conversion to testes (*59, 60, 75*), resulting in ∼1:1 female:male ratio. Sex determination in laboratory zebrafish depends on environmental cues, and the mechanism is not completely understood (*59, 60, 75, 76*). However, the induction of oocyte loss by P53 dependent apoptosis results in conversion to testes (*59, 60*). Multiple oocyte defective mutants, including meiotic mutants, convert to testes and develop as sterile males (*60, 61*), likely activating meiotic checkpoint-induced apoptosis similar to mouse oogenesis (*12, 62*).

The above bouquet and germline cyst phenotypes represent severe oocyte defects, and we tested their consequences on oogenesis. We first tested whether these phenotypes result in oocyte loss by apoptosis. In wt juvenile ovaries, apoptotic oocytes are rare (*77*), as detected by cleaved Caspase3 in wt, *cep290,* and *kif7* heterozygous fish (Fig. 5A). We detected more apoptotic oocytes in *cep290^-/-^*, in *kif7^-/-^*, and in *cep290^+/-^; kif7^-/-^* ovaries, and this was significantly increased in *cep290^-/-^; kif7^+/-^* and *cep290^-/-^; kif7^-/-^* ovaries (Fig. 5A). These results show that ciliary loss results in severe oocyte defects and consequently oocyte loss by apoptosis.

In line with the increased apoptosis levels, we found that in juvenile *cep290^-/-^; kif7^+/-^* and *cep290^-/-^; kif7^-/-^* fish, most gonads converted to testes (Fig. 5B). In all our analyses above we have examined oocyte phenotypes in normal and affected ovaries across the reported genotypes prior to potential conversion. Gonads from these analyses, that were converted and did not include oocytes, were categorized separately as “converted to testis”, and are described here. In wt and heterozygous mutants, we detected up to ∼5% conversion (Fig. 5B), which represents occasional earlier sex determination. In contrast, ∼24% of *cep290^-/-^* ovaries were converted, and ∼60% of *cep290^-/-^;kif7^+/-^*, and ∼80% of *cep290^-/-^;kif7^-/-^* ovaries, where we detected the highest levels of apoptosis (Fig. 5A), were converted (Fig. 5B). Images of normal developing ovaries, normal later-stage developing testes, gonads that converted to testes early, and defective developing ovaries are shown in Fig. S12. Consistent with this, we observed a male bias in *cep290^-/-^* adults (Fig. S13). Most *cep290^-/-^;kif7^+/-^* and *cep290^-/-^;kif7^-/-^* animals do not survive to adulthood, precluding systematic adult analysis, but survivors consistently exhibited male bias as well (Fig. S13). These results, together with the increased level of oocyte apoptosis that we observed, demonstrate that ciliary loss is detrimental to oogenesis.

Ciliary loss could directly cause the above bouquet and germline cyst phenotypes that in turn induce apoptosis, or induce apoptosis which could indirectly cause these phenotypes. Throughout our analyses above, bouquet and cyst phenotypes were clearly detected in oocytes that otherwise appeared viable, suggesting that they preceded apoptosis. To confirm this, we crossed *cep290* fish to *tp53* mutants, which rescues apoptosis (*12, 59, 60*).

We first confirmed that loss of Tp53 completely rescued oocytes from apoptosis in *cep290^-/-^;tp53^-/-^* ovaries (Fig. S14A). The high number of apoptotic oocytes in *cep290^-/-^* ovaries was reduced back to *wt* levels in *cep290^-/-^;tp53^-/-^* ovaries (Fig. S14A). In keeping with this observation, loss of Tp53 function completely rescued juvenile ovary conversion to testes in *cep290^-/-^;tp53^-/-^* mutants (Fig. S14B), and these continued to develop (Fig. S14D). Consequently, the male bias in adult *cep290^-/-^* fish was also rescued in *cep290^-/-^;tp53^-/-^* fish (Fig. S14C). Strikingly, analysis of double mutant juvenile ovaries revealed that later stage oocytes exhibited an extremely abnormal morphology that is never detected in *wt* or *cep290^-/-^* ovaries (Fig.S14D). This demonstrates that in *cep290^-/-^;tp53^-/-^* ovaries, defected oocytes that in *cep290^-/-^* ovaries were otherwise cleared by apoptosis, survived but failed to develop normally, and that the loss of the zygotene cilium is detrimental to oocyte development. These dynamics are consistent with a likely activation of a meiotic checkpoint (which in turn induces tp53-dependent apoptosis) by the defects in bouquet and germline cyst morphology.

We next re-evaluated the ciliary bouquet and cyst phenotypes in *cep290^-/-^;tp53^-/-^* gonads. We reasoned that if the bouquet and cyst phenotypes are caused by oocyte apoptosis, then they should be rescued in *cep290^-/-^;tp53^-/-^* ovaries, whereas if they are caused directly by the loss of cilia, then they should persist in *cep290^-/-^;tp53^-/-^* ovaries.

We first examined the prophase delay from Fig. 2E-F. In *cep290^-/-^* ovaries, “leptotene-zygotene-like” oocytes were mostly oversized (10.4 ±0.917 μm; Fig. S15A-B), compared to the normal sized wt (8.2 ±0.713; Fig. S15A-B), and consistently with Fig. 2E-F. In *cep290^-/-^;tp53^-/-^* ovaries, “leptotene-zygotene-like” oocytes were still mostly similarly oversized (10.42 ±1.102 μm; Fig.S15A-B). We then analyzed telomere clustering in mid-zygotene bouquet oocytes as described in Fig. 2G. Consistent with the data shown in Fig. 2G, in *wt* ovaries most mid-bouquet oocytes showed tight telomere clusters, and in *cep290^-/-^* ovaries most mid-bouquet oocytes showed dispersed or expanded clusters (Fig. S15C-D). In *cep290^-/-^;tp53^-/-^* ovaries, most mid-bouquet oocytes similarly showed dispersed or expanded telomere clusters (Fig. S15C-D). These findings show that these bouquet phenotypes are caused independently of apoptosis.

In *tp53^-/-^* ovaries, we detected slightly over-sized “leptotene-zygotene-like” oocytes (8.945 ±0.933 μm; Fig. S15A-B), and ∼20% mid-bouquet oocytes with dispersed telomere clusters (Fig. S15C-D). We presume that in the absence of oocyte clearance by checkpoint-induced apoptosis, some defective oocytes that are otherwise cleared in wt ovaries, are retained in *tp53^-/-^* ovaries. This is consistent with our detection of extremely defective later-stage oocytes in *cep290^-/-^;tp53^-/-^* ovaries above (Fig. S15D), and with established data in mice showing that defective oocytes continue to grow when not cleared in *tp53^-/-^* or *chk2^-/-^* ovaries (*12*).

We then examined the cyst phenotypes. In *cep290^-/-^* ovaries, cysts were mostly loose or disintegrated (Fig. S16A, C), consistent with Fig. 3A, D. In *cep290^-/-^;tp53^-/-^* ovaries, cysts were similarly mostly loose or disintegrated (Fig. S16A, C). This persistence of cyst phenotypes between *cep290^-/-^;tp53^-/-^* and *cep290^-/-^* ovaries and compared to wt ovaries, suggests that apoptosis is not a primary cause for their induction. Consistent with this, 3D cyst analysis (from Fig. 3B), confirmed the significantly increased normalized distances between neighboring nuclei in cysts of *cep290^-/-^;tp53^-/-^* compared to wt ovaries (Fig. S16B, D), which were similar to those in *cep290^-/-^* ovaries (Fig. S16B, D). While we detected more normal cysts in *cep290^-/-^;tp53^-/-^* ovaries than in *cep290^-/-^* ovaries, >60% of cysts were defective in these ovaries compared to none in the wt and ∼2% in *tp53^-/-^*.

Altogether, all of the above analyses with *cep290^-/-^;tp53^-/-^* ovaries argue that bouquet and cyst phenotypes are not direct consequences of oocyte apoptosis. We conclude that ciliary loss causes bouquet and germline cyst defects that induce apoptosis which is detrimental to oogenesis.

We thus examined adult ovaries and female fertility as the ultimate readout of oogenesis. *kif7^-/-^* females, whose ovaries showed normal or only mildly shortened cilia (Fig. 2A-B) were normally fertile. In contrast, we found that *cep290^fh297/fh297^,* and *cc2d2a^w38/w38^* adult females did not mate spontaneously in either homozygous in-crosses or when homozygous females were crossed to wt males (Fig. S17A), indicating defective oogenesis, or abnormal spawning or mating behavior that might be secondary to their characteristic cilia loss-scoliosis phenotype (*46, 57, 78*). Scoliosis in ciliary mutants results from loss of motile cilia in ependymal cells in the central canal of the spinal cord, which are required for axial straightening (*46, 57, 78, 79*). However, as discussed in the earlier section, since rescued *ccdc103* mutants are fertile despite having profound scoliosis, it is unlikely that spine defects indirectly affect mating and spawning. To specifically test for abnormalities in oogenesis, we performed *in vitro* fertilization (IVF) assays between homozygous mutant females and wt males. Most *cep290^fh297/fh297^,* and *cc2d2a^w38/w38^* females produced degenerated eggs or no eggs at all (Fig. 5C, S17A). In line with their deficient fertility, *cep290^fh297/fh297^* and *cc2d2a^w38/w38^* females exhibited striking ovarian dysgenesis (Fig. 5D, S17C). In comparison to the wt, mutant ovaries were smaller, underdeveloped, and exhibited degenerated masses of tissue (Fig. 5D, S17C). We were able to further confirm these phenotypes in a fifth ciliary mutant that is viable in zebrafish and causes human ciliopathies, *armc9^zh505^*. Armc9 is a ciliary base and tip protein that is required for ciliary stability and involved in Hh signaling (*78, 80, 81*). *armc9^zh505/zh505^* females exhibited similar fertility deficiency in both spontaneous mating and IVF experiments, as well as ovarian dysgenesis (Fig. S17A-C). These data show that the zygotene cilium is required directly in oogenesis and is essential for ovarian development and fertility.

### The zygotene cilium is conserved in male and female meiosis in zebrafish and mouse

Considering the importance of zygotene cilia in oogenesis, we examined whether it also forms during spermatogenesis. In juvenile testes, co-labeling of AcTub with the cytoplasmic membrane marker β-catenin, and with the prophase marker Sycp3, detected cilia-like structures within developing seminiferous tubules, specifically in prophase spermatocytes (Fig. S18A). Sperm flagella are a specialized form of cilia that are composed of similar acetylated tubulin-positive axonemes (*82*). Flagella formation begins after prophase, at mid-round spermatid stages, and completes by the mature spermatozoa stage (*82*). We analyzed developed seminiferous tubules in adult testes, where all spermatogenesis stages can be detected, to distinguish potential zygotene cilia from flagella. We co-labeled AcTub with three prophase spermatocyte markers separately: telomere dynamics as visualized by Telo-FISH (*34*), Sycp3 (*61*), and γH2Ax (*61*) (Fig. 6A-C). Round spermatids had AcTub-positive flagella (E2 in Fig. 6A-C; see staging criteria in the Methods section), and flagella of spermatozoa were very long and swayed within the lumen of the tubules (E3 in Fig. 6A-C). Strikingly, zygotene stage spermatocytes with the characteristic bouquet patterns of telomeres (Telo-FISH), Sycp3, and γH2Ax (*34, 61*), showed clear AcTub-labeled cilia (Fig. 6A-C).

**Figure 6.**
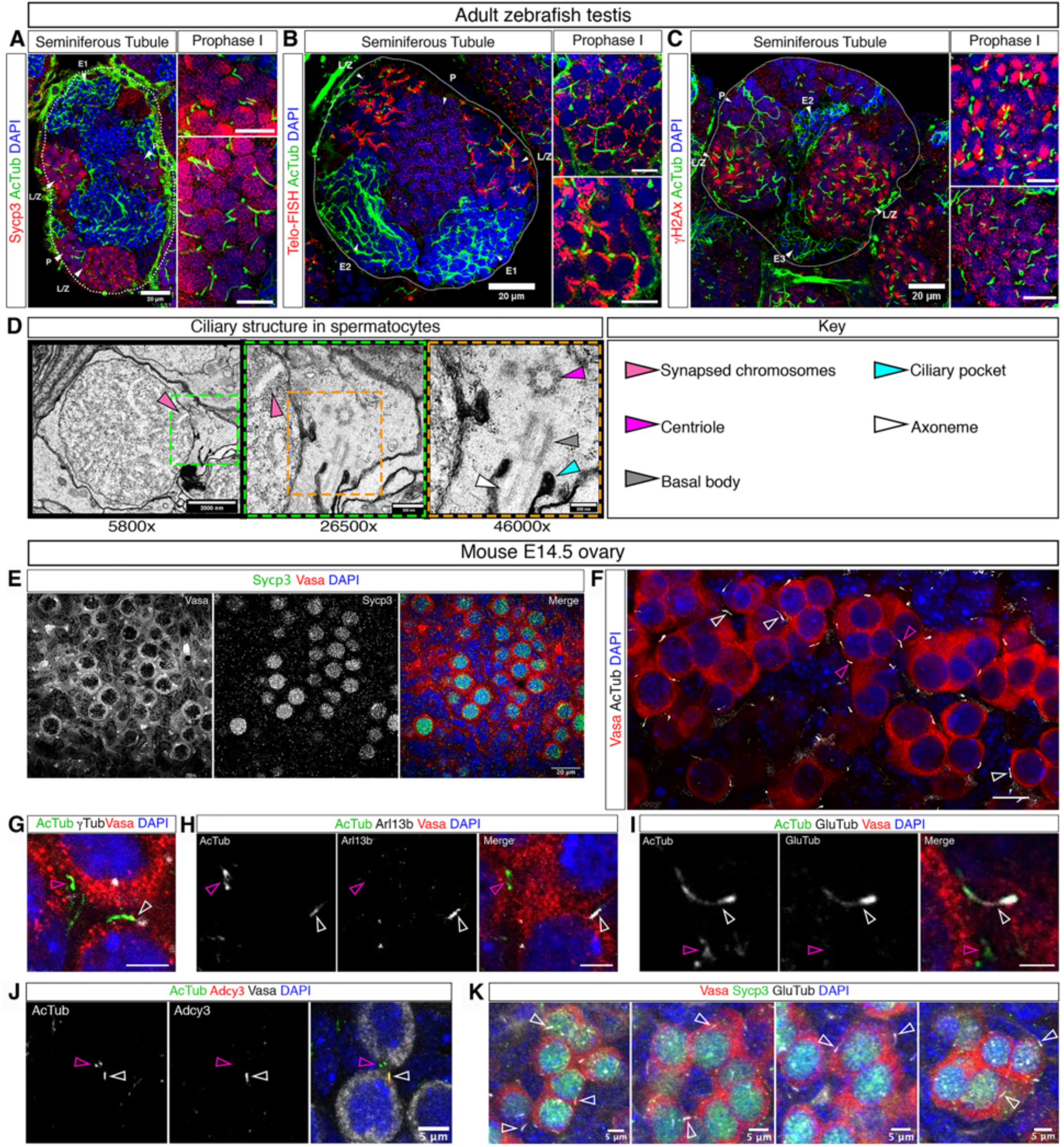
The zygotene cilium is conserved in zebrafish male meiosis and mouse oogenesis. **A-C.** Seminiferous tubules of adult zebrafish testes labeled with AcTub and Sycp3 (**A**, n=35 tubules in 3 testes) or Telo-FISH (**B**, n=29 tubules in 2 testes) or γH2Ax (**C**, n=34 tubules in 3 testes). In **A-C**, left panels are optical sections of entire tubules (white outline, scale bars are 20 μm). The right panels are zoom-in images of prophase spermatocytes (scale bars are 10 μm). L/Z - Leptotene/Zygotene Primary Spermatocytes, P - Pachytene Spermatocytes, E1 - Initial Spermatids, E2 - Intermediate Spermatids, E3 - Final (Mature) Spermatids. **D.** TEM images of adult testis showing ciliary structures (as indicated by arrowheads and the key in the right panel) within (left panels) prophase spermatocytes (identified by their synapsed chromosomes, pink arrowheads). Color-framed panels are larger magnification images of color-matched boxes in left panels. Magnifications and scale bars are as indicated. n=25 spermatocytes in 2 testes. **E.** Mouse E14.5 ovaries labeled with Vasa and Sycp3. Scale bar is 20 μm. **F.** Mouse E14.5 ovaries labeled with Vasa and AcTub, showing AcTub-positive CBs (pink arrows) and cilia (white arrows) in Vasa labeled cysts. n=4 ovaries. Scale bar is 15 mm. **G-J.** E14.5 ovaries labeled with AcTub, Vasa, and γTub (**G**, n=2 ovaries), or Arl13b (**H**, n=2 ovaries), or GluTub (**I**, n=2 ovaries), or Adcy3 (**J**, n=2 ovaries). Scale bars are 5 μm. **K.** E14.5 ovaries co-labeled with Vasa, Sycp3, GluTub, and DAPI. Representative cyst merged images are shown, taken from larger frames, whose individual channels are shown in Fig. S18 (n=2 ovaries). Scale bars are 5 μm. In E-J, cilia (white arrows), and CB-like structures (pink arrows) are indicated.

In pachytene spermatocytes and E1 early round spermatid stages, some showed AcTub-positive cilia while others did not (Fig. 6A, B), suggesting an intermediate time window during/between zygotene cilia removal and flagella formation. TEM analysis confirmed ciliary structures in zygotene spermatocytes (Fig. 6D). These clearly emanated from densely packed spermatocytes (Fig. 6D), distinct from flagella structures in the lumen of seminiferous tubules (*82*) (Fig. S18C). Thus, the zygotene cilium is a separate structure from the flagellum and occurs in the same meiotic stages in spermatogenesis as in oogenesis.

Bouquet mechanisms of meiotic chromosomal pairing are universal in sexually reproducing species (*22-30, 83-86*). Considering this, we examined whether the zygotene cilium is conserved in mammals. In the mouse fetal ovary, oocyte development is synchronized (*87–90*). Sycp3 labeling showed that E14.5 ovaries contained mostly early prophase oocytes (Fig. 6E), as expected (*87–90*). AcTub labeling of E14.5 ovaries identified two structures in Vasa-positive germline cysts (Fig. 6F): one resembled a cytoplasmic bridge (CB), and the other a cilium. We confirmed the ciliary structures by co-labeling AcTub and Vasa with ciliary markers. γTub (Fig 6G), Arl13b (Fig. 6H), GluTub (Fig. 6I), and Adcy3 (Fig. 6J), all confirmed AcTub-positive ciliary structures adjacent to negatively stained AcTub-positive CBs. Co-labeling for Vasa, Sycp3, and GluTub further confirmed that the cilia form in prophase oocytes, and similar to zebrafish, within the germline cyst (Fig. 6K, Fig. S18D).

In addition, we identified ciliary structures in spermatocytes in P12 and P16 mouse testes, using multiple ciliary markers including Arl13b and gluTub co-labeled with Sycp3 (Fig. S19). Furthermore, the ciliary protein ADCY3 co-localized to the centrosome basal body as labeled with γTub (Fig. S19A bottom panels). These ciliary structures in mouse spermatocytes were very short and included mostly a shortly extended basal body (arrowheads in Fig. S19 right panels). Consistent with this, highly similar atypical ciliary structures with yet unknown functions have been previously reported in Drosophila spermatocytes (*91*).

These results demonstrate that the zygotene cilium is conserved between sexes and from zebrafish to mouse, raising the possibility that it is very likely to be an inherent and fundamental part of the meiosis program.

## Discussion

Our work uncovers the concept of a cilium as a newly identified player in meiosis that regulates chromosomal bouquet formation and germ cell morphogenesis. To our knowledge, ciliary involvement in meiosis has not been previously reported in any system. In the zebrafish, analyses of embryonic lethal ciliary mutants, like in the *ift88* gene, have been performed by germline replacement, where chimeric animals with wild type soma but mutant germline were generated (*92*). However, defects in oogenesis and/or ovarian phenotypes were not investigated, although some females with mutant germline were found to be infertile. This suggests partially penetrant defects in oogenesis, which are consistent with the findings reported here.

The bouquet telomere dynamics facilitates the mechanics of chromosomal homology searches for pairing and was first described in 1900 (*31*), but fundamental principles of its activity and regulation are still being uncovered. The proteins that tether the telomeric Shelterin complex to Sun proteins on the inner NE were only recently discovered in animals (*27, 28*), and their functional characterization is under pioneering investigation (*85, 93–95*). Our work now describes the complete framework of the cytoplasmic bouquet machinery as a cable system assembled by the zygotene cilium, centrosome and microtubules. This machinery biomechanically facilitates telomere dynamics for chromosomal bouquet formation in meiosis. We showed that as part of this machinery, the zygotene cilium anchors the centrosome to counterbalance telomere rotation and pulling to form the bouquet, and that bouquet telomere clustering is required for proper synaptonemal complex formation. In this machinery, the cilium could also contribute to meiotic progression and cyst organization by transducing a yet unknown signal.

Supporting evidence for centrosome anchoring strategies during chromosomal rotations exist in other systems. In the fission yeast *S. pombe* bouquet, telomeres associate with the centrosome, called spindle pole body (SPB), which drives back and forth nuclear movements in the cell while associated with telomeres, in an analogous manner to bouquet telomere rotations (*96*). It was shown that astral MT that grow from the SPB are anchored to the cell cortex through Num1p and Mcp5, and that this anchoring is required for nuclear movements and *S. pombe* meiosis (*96–98*). In Drosophila, an analogous bouquet mechanism is executed in premeiotic stages where centromeres (instead of telomeres) are associated with Sun/KASH proteins on the NE and with perinuclear MT (*13, 99, 100*). Centromere rotations drive homology searches very similarly to bouquet rotations in vertebrates (*99, 100*). However, these MTs are not organized from the centrosome, but from the fusome MTOC (*99*).

The fusome is a proteinaceous structure that is specific to Drosophila and connects sister oocytes through CBs like a scaffolding network in the cyst (*101–104*), and thus can potentially provide an efficient anchor. Indeed, the centrosome in this case is not fixed in position, but moves around the nucleus (*99*), providing a case-study that shows the results of its non-anchoring during bouquet movements, consistent with our data in zebrafish showing centrosome dislocation upon ciliary laser excision and in ciliary mutants.

Our findings suggest that telomere clustering in the bouquet is required for proper synaptonemal complex formation. Extensive investigation in mice has established that bouquet telomere rotation provides the mechanism for homology searches and are clearly essential for pairing and fertility (*23-25, 27, 28*). However, mutants in these studies prevented proper loading of telomeres onto the NE, and/or their association with MT, precluding the experimental uncoupling between telomere rotations and their clustering per se (*23-25, 27, 28*). We now show that upon ciliary loss, telomeres still rotate on the NE, but do not cluster, and that in these conditions, Sycp3 fails to load on chromosomes, and defected oocytes are cleared by a P53-dependent apoptosis. On a *tp53* mutant background, ciliary mutants possess defective oocytes that survive but exhibit bouquet and morphological aberrations and fail to develop normally. These data strongly suggest the activation of a meiotic checkpoint, which is expected with aberrant Sycp3 loading and synaptonemal complex formation (*12, 62*).

Our data provide a functional insight into the bouquet configuration. In many species, synaptonemal complex proteins load on chromosomes at sub-telomeric regions. It is plausible that in the bouquet, telomere clustering could contribute to their seeding in assembling the synaptonemal complex. Consistent with this, a recent yeast-based mathematical modelling of chromosomal dynamics and pairing showed that pairing is significantly faster if a bouquet configuration is specifically added to the modelling simulation (*105*). Indeed, in zebrafish, the bouquet telomere cluster was proposed as a hub for chromosomal pairing (*33*). DSB cluster in sub-telomeric regions, and Sycp3 and Sycp1 initially load adjacent to clustered telomeres, specifically at the bouquet stage, from which they extend along chromosomal axes (*33, 61*).

Our findings also shed new light on reproduction phenotypes in ciliopathies, which so far have been explained by sperm flagella defects and defective motile cilia lining the fallopian tubes and efferent ducts (*106, 107*). Our study reveals ciliary functions in early differentiating gametes and in fetal ovaries. Severe pediatric ciliopathic syndromes are lethal or lead to severe developmental defects in young children prior to puberty (*49-52, 108-112*), likely masking reproduction phenotypes. In patients with milder ciliopathies, phenotypes could be variable. In our analyses, truncated cilia produced milder phenotypes than complete ciliary loss. Further, *cep290*, *cc2d2a*, and *armc9* females had deficient fertility, but some were able to produce fertilizable eggs. This is in line with reported pleiotropic and variable ciliopathy phenotypes in humans (*49-52, 108, 109*). Thus, our findings uncover a previously unrecognized mechanism that could additionally contribute to reproduction phenotypes in ciliopathies.

Finally, our work offers the unexpected paradigm that ciliary structures can control nuclear chromosomal dynamics. Most cells in metazoans are ciliated and many exhibit cell type-specific nuclear and/or chromosomal dynamics, implying that ciliary regulation of nuclear events may be widely conserved.

## Acknowledgments

We thank M.C. Mullins, C. Moens, P. Ingham, S. Farber, Z. Sun, and B. Perkins, for generously sharing fish lines, and Z. Sun for the zebrafish Arl13b antibody. We thank Y. Buganim for his help with our mouse ovary work. We thank S. Burgess and M Kluetstein for their help with Sycp3 labeling. TEM analysis was performed with the help of Y. Friedmann, The Bio-Imaging Unit, at The Alexander Silberman Institute of Life Science of The Hebrew University of Jerusalem. The SBF-SEM analysis was performed with the help of A. Beckett, Biomedical EM Unit, at The University of Liverpool. We thank Z. Manevich from the Faulty of Medicine Microscopy core and Y. Addadi from the Weizmann Institute core facility for their help with laser ablation experiments.

## Author contributions

Conceptualization: AM, VK, QT, RBG, SR, YME

Methodology: AM, VK, QT, RD, NH, AS, AA, KL, SAL, RYB, NE, RBG, SR, YME

Investigation: AM, VK, QT, RD, NA, KL, MM, FN, HE, SR, AS

Visualization: AM, VK, QT, RD, KL, MM, RBG, SR, YME

Funding acquisition: SR, YME Supervision: RGB, SR, YME

Writing – original draft: AM, VK, QT, MM, RBG, SR, YME

## Supplementary information

[Materials and Methods, Supplementary figures, Captions for supplementary videos]

## Materials and Methods

### Fish lines and gonad collections

Juvenile ovaries were collected from 5-7 week post-fertilization (wpf) juvenile fish. Fish had a standard length (SL) measured according to (*113*), and were consistently ∼10-15 mm. Ovary collection was done as in (*4, 35*). Briefly, to fix the ovaries for immunostaining, and DNA-FISH, fish were cut along the ventral midline and the lateral body wall was removed. The head and tail were removed and the trunk pieces, with the exposed abdomen containing the ovaries, were fixed in 4% PFA at 4°C overnight with nutation. Trunks were then washed in PBS and ovaries were finely dissected in cold PBS. Ovaries were washed in PBS and then either stored in PBS at 4°C in the dark, or dehydrated and stored in 100% MeOH at -20°C in the dark. Testes were collected similarly from 7 wpf juvenile fish and from adult fish and processed similarly. Adult ovaries were collected form wt and mutant fish using similar microdissection and were imaged using a Leica S9i stereomicroscope and camera.

Fish lines used in this research are: TU wild type, *cep290^fh297^* (*46*), *kif7^i271^* (*53*), *cc2d2a^w38^* (*57*)*, tp53^M214K^*(*114*)*, Tg(β−act:Arl13b-GFP)* (*115*), *Tg(h2a:H2A-GFP)* (*116*), *Tg(β−act:Cetn2-GFP*(*117*), *Tg(foxj1a:GFP)*(*72, 118*)*, Tg (foxj1b:GFP)*(*72, 118*), *Tg(β−act: mCherry-Cep55l), armc9^zh505^* (*81*). The *Tg(β−act: mCherry-Cep55l)* line was generated using the *tol2* system. *mCherry-cep55l* was PCR amplified from a *mCherry-cep55l C1* vector (*119*) and cloned into *pDest-Tol2* plasmid under *β-actin* promoter, using Gibson assembly (E5510S, New England Biolabs, Ipswich, MA). Other fish lines analyzed for fertility included.

### Staging of prophase oocytes

Oocyte prophase stages were determined based on established staging criteria (*4, 35*) (Fig. S1B). We established prophase staging criteria for zebrafish oocytes based on the characteristic telomere dynamics at each stage (*4, 35*). Briefly, telomeres are distributed intranuclearly in mitotic oogonia, then load on the nuclear envelope (NE) at the onset of meiosis, and remain radially distributed on the NE through leptotene-zygotene stages (including during chromosomal rotations). Telomeres are clustered at one pole of the nucleus during zygotene bouquet stages. Later they disperse radially and dissociate back into the nuclear vicinity (*4, 35*). We characterized cellular features, including the typical oocyte size and nuclear morphology that strictly and distinctively accompanied each telomere distribution, such that they identify each stage even without telomere labeling (*4, 35*) (Fig. S1B). We developed a reliable method for measuring oocyte size in diameter in three dimensions (*35*), which we routinely use for determining the oocyte size as part of its staging criteria. For example, oogonia with dispersed intranuclear telomere distribution was always ∼10 μm with non-condensed DAPI-labeled DNA and central 1-3 nucleoli, leptotene-zygotene stages with telomere radially loaded on the nuclear envelope (NE), were always 7-9 μm with central few nucleoli, and zygotene bouquet with clustered telomeres on the NE were always 10-16 μm with condensed DAPI-labeled chromosomes and a peripheral single nucleolus (*4, 35*) (Fig. S1B).

In this study, to further precisely determine the specific leptotene-zygotene stages of prophase oocytes we used the established nuclear patterns of the synaptonemal complex protein Sycp3. We identified the characteristic patterns to each prophase stage as established in (*33, 34*). The leptotene stage is characterized by an elongated patch of Sycp3 signal in the oocyte nucleus (*33, 34*) (Fig S1C). Early Zygotene is characterized by a polarized Sycp3 signal that corresponds to the bouquet telomere cluster where it begins to load on chromosomes in zebrafish (*33, 34*) (Fig S1C). During late Zygotene, the Sycp3 signal extends from its polarized location along chromosomal axes (*33, 34*) (Fig S1C). We measured oocyte size per each Sycp3 pattern in three dimensions in wholemount ovaries, as described in (*35*). Fig. S1C shows Sycp3 patterns and their corresponding oocyte sizes. Note that this Sycp3-based oocyte sizes are fully consistent with these based on Telo-FISH and morphological features above, further confirming our staging criteria. In all our analyses we always include cytoplasmic labeling (like DiOC6), or use the background of other labeling to visualize oocyte cytoplasm, measure oocyte size in three dimensions (*4, 35*), and determine its prophase stage based on these size and morphological criteria.

### Fluorescence immunohistochemistry (IHC), and DNA-Telo-FISH

IHC was performed as in (*4, 35*). Briefly, ovaries were washed 2 times for 5 minutes (2×5min) in PBT (0.3% Triton X-100 in 1xPBS; if stored in MeOH, ovaries were gradually rehydrated first), then washed 4×20min in PBT. Ovaries were blocked for 1.5-2 hours (hr) in blocking solution (10% FBS in PBT) at room temperature, and then incubated with primary antibodies in blocking solution at 4°C overnight. Ovaries were washed 4×20min in PBT and incubated with secondary antibodies in fresh blocking solution for 1.75 hr, and were light protected from this step onward. Ovaries were washed 4×20min in PBT and then incubated in PBT containing DAPI (1:1000, Molecular Probes), with or without DiOC6 (1:5000, Molecular Probes) for 50 min and washed 2×5min in PBT and 2×5min in PBS. All steps were carried out with nutation. Ovaries were transferred into Vectashield (with DAPI, Vector labs). Ovaries were finally mounted between two #1.5 coverslips using a 120 μm spacer (Molecular Probes).

Primary antibodies used were γTubulin (1:400, Sigma-Aldrich), GFP (1:400; Molecular Probes), Vasa (1:5000) (*120*), Acetylated tubulin (1:200; Sigma-Aldrich), β-Catenin (1:1000; Sigma-Aldrich), Arl13b (*121*), glutamylated tubulin (1:400; Adipogen), Sycp3 (1:200, Abcam), γH2Ax (1:400,GeneTex), cCaspase3 (1:300, Abcam). Secondary antibodies were used at 1:500 (Alexa-flour, Molecular Probes).

DNA-FISH for telomeric repeats (Telo-FISH) was performed using the PNA technique (PNA-Bio) following the company protocol. Hybridization buffer was 70% Formamide, 1 mM Tris pH 7.2, 8.5% MgCl2 buffer (25 mM magnesium chloride, 9 mM citric acid, 82 mM sodium hydrogen phosphate, pH7), 1x Blocking reagent in Maleic acid buffer (100 mM Maleic acid, pH7.5), 0.1% Tween20, 88 nM probe (*5’-CCCTAACCCTAACCCTAA-3’,* Cy3- conjugated).

For a combination of IHC with Telo-FISH, IHC was performed first. At the end of the IHC procedure, ovaries were washed an extra time for 30 min in PBT and fixed quickly in 4% PFA for 15-20 min at room temperature. After staining was complete, DAPI (+/- DiOC6) staining and mounting was performed as described above.

### Confocal microscopy, image acquisition and processing

Images were acquired on a Zeiss LSM 880 confocal microscope using a 40X lens. The acquisition setting was set between samples and experiments to: XY resolution=1104×1104 pixels, 12-bit, 2x sampling averaging, pixel dwell time=0.59sec, zoom=0.8X, pinhole adjusted to 1.1μm of Z thickness, increments between images in stacks were 0.53μm, laser power and gain were set in an antibody-dependent manner to 7-11% and 400-650, respectively, and below saturation condition. Unless otherwise noted, shown images are partial Sum Z-projection. Acquired images were not manipulated in a non-linear manner, and only contrast/brightness were adjusted. All figures were made using Adobe Photoshop CC 2014.

### Confocal live and time-lapse imaging of cultured ovaries

Live imaging was performed as in (*35*). Briefly, ovaries were dissected from juvenile fish (5-7wpf, SL∼10-15mm) into fresh warm HL-15 media (Hanks solution containing 60% L-15 (no phenol red), Sigma-Aldrich, and 1:100 Glutamax, Invitrogen, warmed to 28°C). Ovaries were then embedded in 0.5% low-melt agarose in HL-15 on a glass-bottom dish, and covered with warm HL-15. After the agarose polymerized, ovaries were incubated in HL-15 at 28°C. Time-lapse images of live ovaries were acquired using either a Zeiss LSM 880 or Nikon Ti2E spinning disc confocal microscopes, both equipped with an environmental chamber set to 28°C, and using a 40X and 60X lenses, respectively. Images were acquired every 6 seconds for single Z section, or 18-20 seconds for Z stacks, over 5-10 minutes.

### Chromosome rotation tracking in live time-lapse images

Live time-lapse imaging of *Tg(h2a:h2a-gfp)* cultured ovaries were performed as above, recording large frames of ovaries, that included multiple zygotene oocytes, oogonia, and somatic pre-granulosa cells side by side, and under identical conditions. Chromosomal dynamics in these cells were tracked based on H2A-GFP intensities, using the TrackMate plugin on ImageJ (*122*) with the following setting parameters: Blob Diameter was set to 3 microns with an appropriate threshold, and Linking distance was set to 4 microns with maximum permissibility of 2 frame gap for Gap-closing and Gap-closing maximum distance of 1 micron. Identified tracks were confirmed manually. Selected tracks were identified over 20-30 consecutive time points and at least 6. consecutive time points were selected, and images with at least 2 appropriately identified tracks were analysed. For track visualization, spot sizes were set to 0.2, Tracks were set to show tracks backward, and were color-coded for time. Tracks were then recorded as an overlay on the original image using the “Capture Overlay” function. Track parameters, including velocity, displacement, and track duration were exported to Excel and GraphPad Prism for statistical analysis.

### Coloc2 analysis

A pixel-wise quantitative test for colocalization of AcTub with Arl13b and with GluTub signals was performed using the Coloc 2 plug-in on Fiji as in (*4*). Cilia regions of interest (ROI) were taken along individual cilia from representative images. The shape of the ROI varies according to the cilia orientation: ROI of straight cilia are narrower, while ROI of bent or curled cilia are wider, however the spaces taken around the cilia were consistent in all ROI. We calculated a point spread function (PF) of ∼1.6 pixels on average for each channel and image, based on PSF= [0.8xExitation wavelength (nm)/objective’s NA]/pixel size (nm). Min and max thresholds were automatically set by the plug-in for each image. Pearson correlation coefficient (R) was calculated. R values represent: -1<R<0, anti-correlation; R=0, no correlation; 0<R<1, correlation. The plug- in then scrambles the pixels to generate a random image and test for the probability to yield the same Pearson coefficient from a random image. Scrambling was set to 100 iterations, generating 100 different random images from the tested image’s pixels. The probability of receiving the calculated Pearson coefficient values for both AcTub and Arl13b and AcTub and gluTub were *p*=1. A pixel-wise quantitative test for colocalization of GluTub with Cep55 was performed similarly.

### Image 3D reconstructions in IMARIS

ROIs of whole cysts were extracted from confocal raw data and used to reconstruct nuclei, cilia and other cellular features with respect to each other using blend volume rendering mode. Signal brightness and object transparency were edited in all channels to optimize signal and reduce background. Animation frames were made using the Key Frame Animation Tool.

### Serial Block Face Scanning Electron Microscopy (SBF-SEM)

Resin embedded ovaries were mounted onto a cryo pins using conductive silver epoxy and targeted trimming was performed using an ultra-microtome (leica, Milton Keynes, UK) to expose specific area of the ovaries. The trimmed block was painted with Electrodag silver paint and coated with 10 nm AuPd using a Q150T sputter coater (Quorum Technologies).

Cysts were imaged using a Gatan 3View (Gatan, Pleasanton, USA) mounted on a Quanta 250 SEM (FEI, Hillsboro, Oregon, USA). Two areas containing cysts were imaged using the multiple regions of interest (ROI) function, ROI-00, ROI-01. Imaging conditions were as follows; ROI 00, magnification 3445x, pixel size 5.9 nm in x and y, 75nm in z, image dimensions 8189 × 5932 pixels. ROI 01, magnification 3480x, pixel size 5.9 nm in x and y, 75nm in z, frame width 8192 × 5932. Chamber pressure of 70 Pa. 2.8kV, dwell time per pixel 68μs for both RIO’s. Before converting the images into Tiff format with Digital Micrograph, they were binned by two giving a final pixel size in x and y of 12nm.

### 3-D rendering of SBF-SEM images

Renders were made from over 330 75nm sections for each cyst (Movie S3). The image sequence generated from the Serial Block Facing Scanning Electron Microscope (SBF-SEM) was imported into ImageJ. The imported image was then adjusted with the correct voxel size, brightness, contrast, and alignment. The stack was imported into TrakEM2 canvas for segmentation. In TrakEM2, a project tree was made by adding area lists. Once the Area list was made cell were manually segmented at every 7^th^ to 8^th^ slice using the brush tool of ImageJ and the gap is filled by using the “Interpolation function” of TrakEM2. The segmented structure of the cyst was then exported as “TIFF” file to IMARIS software, and Image properties were set to the acquisition voxel size. Surfaces were created using the surface creation wizard and colour and transparency were set for individual cells. The segmented cells were then exported as “Scenes”. This pipeline was repeated for different structures of the cyst and exported as different scenes from the Area lists. Then the SBF-SEM image stack was imported to the “Arena” area of IMARIS. The voxel size was corrected, and Scenes were imported. Once all the scenes were imported, colour and transparency were set for individual elements for optimal representation. The reconstruction was finally saved using the “Animation” function of IMARIS (Movies S3, 6).

### Rescue of *ccd103* mutant embryonic lethality

The mRNA of zebrafish ccdc103 was transcribed from template with the mMessage mMachine SP6 transcription kit (Invitrogen, AM1340). To rescue the mutants, total 0.5 nl of the mRNA with final concentration of 150 ng/ul was injected into the animal pole of one-cell stage embryos. Rescued *ccd103^-/-^* females were confirmed by genotyping, and exhibited the *ccd103* characteristic scoliosis phenotype, which results from loss of cilia in ependymal cells in the spine central canal.

### Measurements of cilia and CBs in SBF-SEM data

The image sequence was imported as a stack in Fiji. The Image voxel size was set as per the acquisition metadata by selecting image properties. Slices where cytoplasmic bridge or cilia were widest were selected. Using the line tool from the Fiji Menu bar, measuring lines were drawn across the width of the cilia or CBs. The lines drawn were color-coded Red from overlay properties and added to the ROI manager. Three measurement lines were drawn across cilia and CBs longitudinal sections. Two perpendicular lines were drawn across cilia cross sections (along perpendicular diameters). All lines in ROI manager were measured and the result table is saved as a CSV file, and exported to Excel for calculating average width of Cytoplasmic Bridge and Cilia and graphical representation is produced using Graphpad Prism 8.

### Measurements of cilia length in confocal data and defining ciliary length categories in wt and mutant ovaries

10 representative cyst ROIs from each genotype were cropped in FIJI and used for image processing in the IMARIS 3D software. The filaments IMARIS feature was used to mark the cilia pathways, as follows. To create the cilium filament, the “Auto-depth” semi-automatic IMARIS algorithm was used, combining user’s manual tracking and local signal intensity (Fig. S6). After creating filaments for all cilia in a cyst, the filament lengths were extracted to EXCEL and GraphPad Prism for statistical analysis, and the images of cilia tracks were extracted as JPEG (Fig. S6).

We first measured ciliary lengths in wt cysts in 3D as described above and defined ciliary length categories. Wt cilia mostly ranged between 4-8.5 μm, with the longest cilia reaching 12 mm (Fig. 2A). We defined ciliary lengths at 4-8.5 μm as “normal cilia”, and cilia with outlier values <4 μm as “short cilia”. Cilia that were only detected as an extremely short dot-like signal that was <1 μm, or not detected at all, were defined as “absent cilia” (N.D in Fig. 2A-B). Individual cilia were measured as described above in cysts from all genotypes (Fig. 2A), and the average ciliary length per cyst was calculated (Fig. 2B). Cysts with averages >4 mm were scored cysts with “normal cilia”, cysts with averages <4 μm, were scored as cysts with “short cilia” (Fig. 2B). Distribution of scored cysts were plotted (Fig. 2C) and entire ovaries were scored and their distribution for the same categories was plotted (Fig. 2D).

### Bouquet analysis

Zygotene bouquet stages span oocytes sizes of 10-16 μm in diameter (*4, 35*), and the telomere cluster might spread wider during early bouquet formation (∼10 μm), and later deformation (∼16 μm)(*35*). We focused on oocytes with clear telomere clusters at mid-bouquet stages (11.5-13 μm)(*4, 35*). We co-labeled telomeres by Telo-FISH and the cytoplasm with DiOC6 (Fig. 2G). We scored all oocytes blind to the Telo-FISH channel, measured their size in diameter based on the DiOC6 cytoplasmic labeling according to (*35*), and selected oocytes at mid-bouquet sizes (11.5-13 μm). We then used the Telo-FISH channel to analyze the distribution of telomeres in all Z sections spanning these entire oocytes. Categories were defined as “Tight” for telomere clusters at one pole of the nucleus that span up to approximately a quarter of the nuclear circumference, “Expanded” for wider clusters that spread beyond a quarter of the nuclear circumference and up to the nuclear equator, and “Dispersed” for unclustered telomeres that spread over the nuclear equator as shown in Fig. 2G. The analysis in Fig. S7D, which addresses later bouquet stages, was performed identically, but selected oocytes were at a size range of 13-15 μm.

### Ciliobrevin treatments

Ovaries were dissected from juvenile wt fish and cultured in HL-15 as described in the “Confocal live and time-lapse imaging of cultured ovaries” section above. The HL-15 media was then replaced with HL-15 media containing either DMSO, or 15, or 50 μM ciliobrevin. Ovaries were incubated for 80 minutes at 28°C, and were fixed for Telo-FISH, DiOC6, and DAPI labeling. Bouquet analysis was performed on 11.5-13 μm oocytes as described above.

### Ciliary laser excision

Ovaries were dissected from 6 wpf *Tg(βact:Arl13b-GFP);Tg(βact:Cetn2-GFP)* fish and mounted as described for live imaging above. Excisions were performed using a Leica TCS SP8 MP two-photon microscope with a 25X objective and equipped with an incubation chamber set to 28°C. A region of interest with good number of cilia in the field of view was selected. The laser stimulation region for excision was drawn manually using the “Draw ROI” function. Time-lapse was recorded for 60 seconds pre-ablation, followed by a laser stimulation (900nm, 2.79 watt) for 20 seconds, and post-ablation time-lapse recording for 8 minutes (Movies S13-14). Control experiments (Movies S15-16) were performed side-by-side with the experimental ablations, under identical conditions and settings, except laser power was set to 4 watt for 20 seconds.

### Centrosome-nucleus distance measurements

The distance between the centrosome and the nucleus was measured in wt and mutant ovaries co-labeled for centrosome (γ-Tub antibody) and telomeres (Telo-FISH), and counter-stained with DAPI. Oocyte at mid-zygotene bouquet stage were selected as in the bouquet analysis above (oocyte size 11-13 μm). For centrosome-nucleus distance measurements, the optical section that includes the center of the centrosome was selected, and the distance between the centrosome and the closest point on the nucleus surface was measured in FIJI. Measurements were tracked using the FIJI ROI manager. We then independently scored the telomere clusters of measured oocytes to the “Tight”, “Expanded” and “Dispersed” categories according to the bouquet analysis above. Centrosome-nucleus distances were normalized to oocyte size (diameter) and plotted per genotype, color-coded for telomere cluster categories (Fig. 3C). Pooled oocytes from all genotypes in this analysis were also plotted per telomere cluster category (Fig. 3D).

### Sycp3 phenotype analysis

To analyze effects on Sycp3 in wt and mutant wholemount ovaries labeled for Sycp3 and DAPI, we scored oocytes at three stages and sizes based on our wt prophase staging in Fig. S1B-C: early zygotene (10-12 μm) and late zygotene (12-14 μm). For each stage, we selected oocytes of the appropriate size range, and then scored the Sycp3 signal as either normal or defected (Fig. 3E), compared to wt and in Fig. S1B-C.

### Quantitative analysis of cyst morphology and integrity

We defined phenotype categories for germline cyst morphology, as follows. “loose cysts” category showed abnormal gaps between sister oocytes in cysts (Fig. 4A). “disintegrated cysts” category phenotype was more severe, showing larger abnormal gaps between cyst oocytes (Fig. 4A), and clear elongated cytoplasmic bridges extending between distant oocytes (Fig. 4A). Normally, at later stages when oocytes leave the cyst, their CBs are detected by MT labeling and appear elongated as they disjoin from their sister oocytes (*4*). Within normal cysts, CBs are very challenging to detect by either MT(*4*) or cytoplasmic labeling without a specific marker (Fig. 4A). This is likely because the cytoplasmic membranes of oocytes interface tightly, masking the fine and short structure of CBs (Fig. S4). In “disintegrated cysts” the elongated CBs are visible without labeling of MT or specific markers, and are similar to the MT-detected CBs of oocytes that naturally leave the cyst later (*4*). They likely represent CBs of oocytes that are separating apart.

For quantitative analysis of category phenotypes, and because borders between tight cells in wt cysts are difficult to detect, we measured the distances between the centers of all the neighboring nuclei in 3D in wt, *cep290^-/-^,* and *cep290^-/-^; kif7*^+/-^ cysts (Fig. 4B, Fig. S10B, Movie S17). All distances were normalized to the average oocyte diameter per cyst (Fig. 4B, Fig. S10B, right panels). 5-8 representative cysts ROIs from each phenotypic category were cropped in FIJI and used for image processing in the IMARIS 3D software. We used semi-automatic segmentation approach to create nuclei surfaces. Each nucleus was segmented in each Z-layer to create a full 3D surface, either by the software automatic segmentation (intensity based) or by marking the nucleus contour manually. After creating surfaces for all cyst nuclei, we used the measurement point feature to calculate the distances between the center of mass of all neighboring nuclei. Distance measurements were exported to an excel sheet and normalized to the average oocyte size (diameter in μm; measured as in (*35*)) per given cyst. Normalized distances were exported to Graphpad Prism for statistical analysis between groups and generation of dot plots. Images of processed cysts were exported from IMARIS.

### Correlation analysis of cyst integrity and ciliary phenotypes

In all cyst morphology experiments we co-labeled cilia by AcTub staining and simultaneously analyzed cyst and cilia phenotypes. To analyze the correlation between cyst and ciliary phenotypes, we pooled gonads from across all examined genotypes and categorized them by the effect on cilia, i.e., normal, short (truncated), or absent (as in Fig. 2A-D), regardless of their genotype. We then asked how many cysts per ciliary phenotypic category were observed in each ovary, and based on this distribution, labeled ovaries as normal, mildly effected, and severely affected (n=43 gonads, 10-60 cysts were analyzed per gonad). We next categorized cyst phenotypes in these ovaries and considered “disintegrated” and “isolated oocytes” categories as severe effects. In most ovaries, a certain cyst phenotype category was predominant in at least 50% of cysts. The ovary category was termed according to this cyst category, or to the most frequently detected one in few cases where no category was frequent in 50% of cysts. The frequency of cyst severity phenotypes were plotted for each ciliary phenotype (Fig. 4F). *cc2d2a* mutants were analyzed identically (Fig. S10C).

### Determining stages of spermatogenesis

We identified all spermatogenesis stages in tubules based on their characteristic nuclear morphology and size, using DAPI counter-stain according to(*123*), as well as the prophase markers telo-FISH, Sycp3 and gH2Ax, as described in the text. Briefly, round spermatids had small round nuclei (2.5 ± 0.1 μm in diameter) and were found towards the center of the tubule as expected. Spermatozoa had compact round nuclei (2.1 ± 0.1 μm in diameter) and were found in the lumen of the tubules as expected. Early round spermatids had a nucleus diameter of 3± 0.1 μm. Round spermatids and spermatozoa were negative to all three prophase markers. Nuclei diameter of Leptotene-Zygotene spermatocytes (L/Z) and pachytene spermatocytes (P) were 5.1+/-0.1 mm and 6+/-0.1 mm, respectively, and both were positive for all prophase markers, showing the expected pattern of each stage (*34, 61*).

### Transmission Electron microscopy (TEM)

Gonads were dissected from juvenile fish (5-7 wpf, SL∼10-15mm) and fixed as described above in PBS containing 4% PFA and 2.5% glutaraldehyde for overnight at 4°C. Gonads were then rinsed 4 times, 10 minutes each, in 0.1M cacodylate buffer (pH 7.4) and post fixed and stained with 1% osmium tetroxide, 1.5% potassium ferricyanide in 0.1M cacodylate buffer for 1 hour. Ovaries were washed 4 times in cacodylate buffer followed by dehydration in increasing concentrations of ethanol consisting of 30%, 50%, 70%, 80%, 90%, 95%, for 10 minutes each step followed by 100% anhydrous ethanol 3 times, 20 minutes each, and propylene oxide 2 times, 10 minutes each. Following dehydration, the ovaries were infiltrated with increasing concentrations of Agar 100 resin in propylene oxide, consisting of 25, 50, 75, and 100% resin for 16 hours each step. Gonads were embedded in fresh resin and let polymerize in an oven at 60^0^C for 48 hours.

Embedded gonads in blocks were sectioned with a diamond knife on a Leica Reichert Ultracut S microtome and ultrathin sections (80nm) were collected onto 200 Mesh, thin bar copper grids. The sections on grids were sequentially stained with Uranyl acetate for 10 minutes and Lead citrate for 10 minutes and imaged with Tecnai 12 TEM 100kV (Phillips) equipped with MegaView II CCD camera and Analysis**®** version 3.0 software (SoftImaging System GmbH). For TEM images, only the “brightness/contrast” functions on Adobe Photoshop, were mildly adjusted. These adjustments did not affect the biological properties of the imaged cellular features.

### In vitro fertilization (IVF)

IVF was performed as in (*124*). Briefly, sperm was collected from anesthetized males into Hank’s solution and stored on ice until eggs are collected. Sperm from two males of the same genotype was collected into a single tube and was used to fertilize eggs collected from control wt, or *cep290^fh297^, or cep290^fh378^*, or *cc2d2a^w38^*, or *armc9^zh505^* homozygous females. Anesthetized wt or mutant females were placed in a dish and squeezed for egg collection. 100-150ul sperm solution was added to the collected eggs, and incubated for 20 seconds. 1ml of E3 containing 0.5% fructose was added to activate sperm, gently mixed and incubated for 2 minutes. 2 ml of E3 was added followed by 5 minutes incubation. The dish was then flooded with E3 and placed in a 28C incubator until examination of fertilization rates.

### Mouse ovaries and IHC

Ovaries from fetal mice at day E14.5 were first fixed in trunk with 4% paraformaldehyde in PBS at room temperature for 2 hours, before dissected out and washed briefly in PBS. The ovaries were further permeabilized with 0.3% TritonX-100 in PBS for 30 min at room temperature, followed by blocking with 3% BSA in 0.3% TritonX-100 in PBS for 1 hour at room temperature. Primary antibodies incubation was performed at 4 ℃ for overnight. After 6×20 min washing with 0.3% TritonX-100 in PBS, the ovaries were incubated with secondary antibodies at room temperature for 3 hours. The stained ovaries were finally washed 6×20 min with 0.3% TritonX-100 in PBS, and mounted in 70% glycerol.

Testes from P12 and P16 mice were fixed in trunk with 4% paraformaldehyde in PBS at room temperature for 2 hours, followed by dissection to isolate the testes. The whole testes were then proceeded with cryo-embedding. Representative cross-sections of the testes were acquired at anterior, posterior and middle region, with thickness of 20 μm. To start the testes staining, cryo-sections on the slides were first thawed and dried before drawing borders around the sections with a PAP pen (Abcam, ab2601). The slides were then fixed with 4% PFA for 15 min at room temperature in Coplin jars (all subsequent steps were performed in Coplin jars unless stated otherwise), rinsed twice with ice-cold PBS followed by permeabilization with 0.2% Triton (in PBS) for 15 min. Next, slides were washed 3 times, 5 min each in PBS and blocked with 2% BSA in PBS for 2 h at room temperature. The slides were then transferred to a humidified box.

Primary antibodies in PBS (with 0.1% Tween20 and 1% BSA) were pipetted onto the sections and incubated overnight at 4°C. **S**lides were washed with PBS on a shaker, six times for 10 min each at room temperature. Secondary antibodies in PBS (with 0.1% Tween20 and 1% BSA) were then added, and the slides incubated in the humidified box for 5 h at room temperature. Finally, the slides were washed six times (10 min each) with PBS at room temperature, dried and mounted before imaging.

Primary antibodies used to stain mice ovaries or testes cryo-sections included: AcTub (1:500, Sigma-Aldrich), Vasa (1:500, R&D systems), γTub (1:500, Adipogen), Arl13b (1:500, Proteintech), Adcy3 (1:500, LSBio). Alexa-flour fluorescent secondary antibodies (Invitrogen) were used at 1:500.

### Statistical analysis

All statistical analysis and data plotting was performed using the GraphPad Prism 7 software. Data sets were tested with two-tailed unpaired *t*-test, except with *Chi-square* Fisher’s exact test when indicated. *p*-values were: *<0.05, **<0.01, ***<0.001, ****<0.0001, ns=not significant (>0.05).

---

**Supplementary Figure 1.**
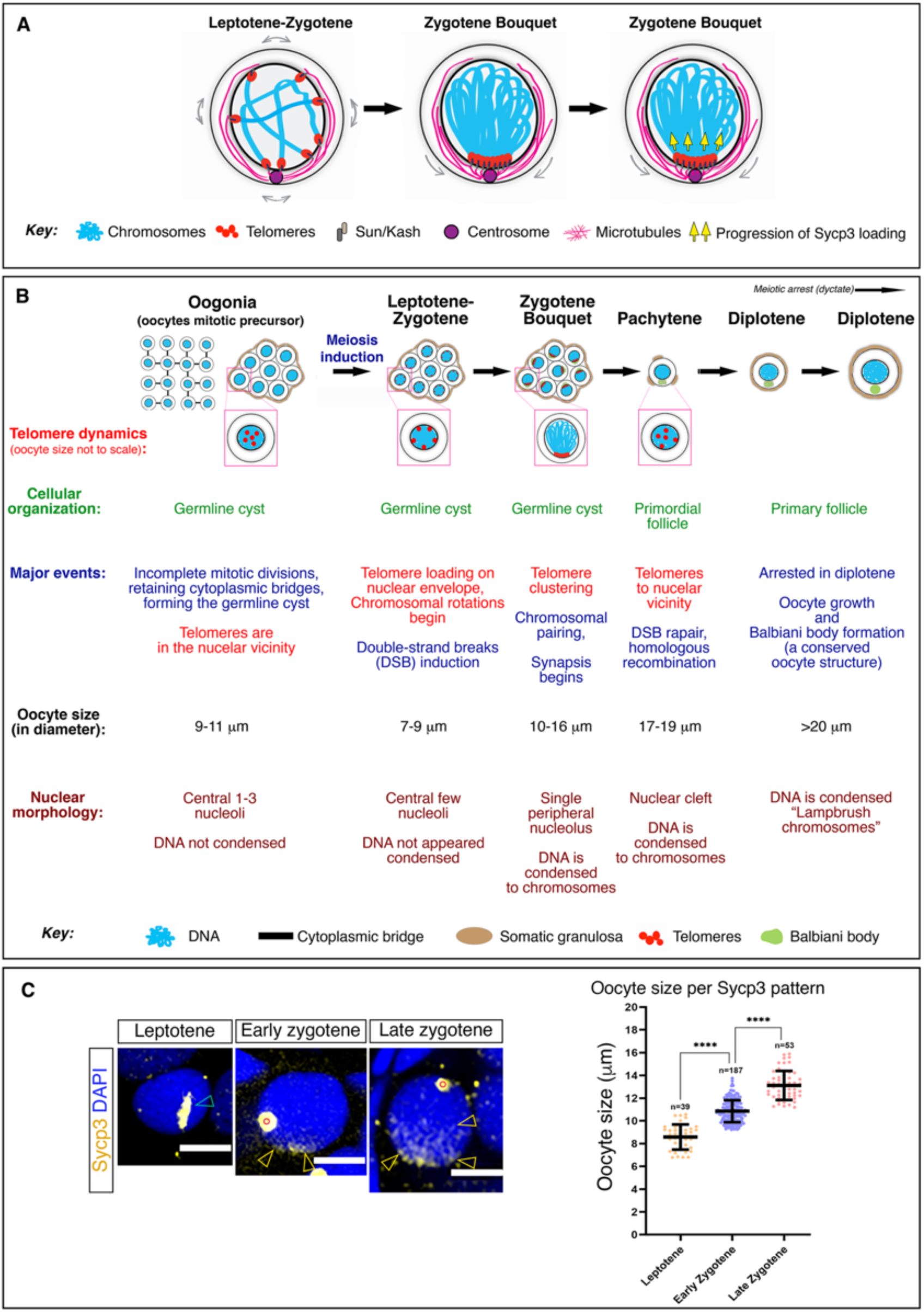
The zygotene chromosomal bouquet and early oogenesis in zebrafish. **A.** A schematic representation of the chromosomal bouquet configuration and its formation. **B.** A layout of early oogenesis in zebrafish, showing the telomere dynamics, cellular organization and major evens for key prophase stages, as well as characteristic features like oocyte size and nuclear morphology that are established as staging criteria. Both panels are modified from (12). **C.** Sycp3 labeling in early prophase. Left panels are representative images of Sycp3 patterns as established in leptotene (oval puncta), early zygotene (polarized as appears to load on sub- telomeric regions), and late zygotene (polarized with apparent extension of loading along chromosomal axes). Right panel shows oocyte size in diameter corresponding to each Sycp3 pattern, and consistent with our Telo-FISH based criteria in B. n=314 oocytes, from 10 ovaries.

**Supplementary Figure 2.**
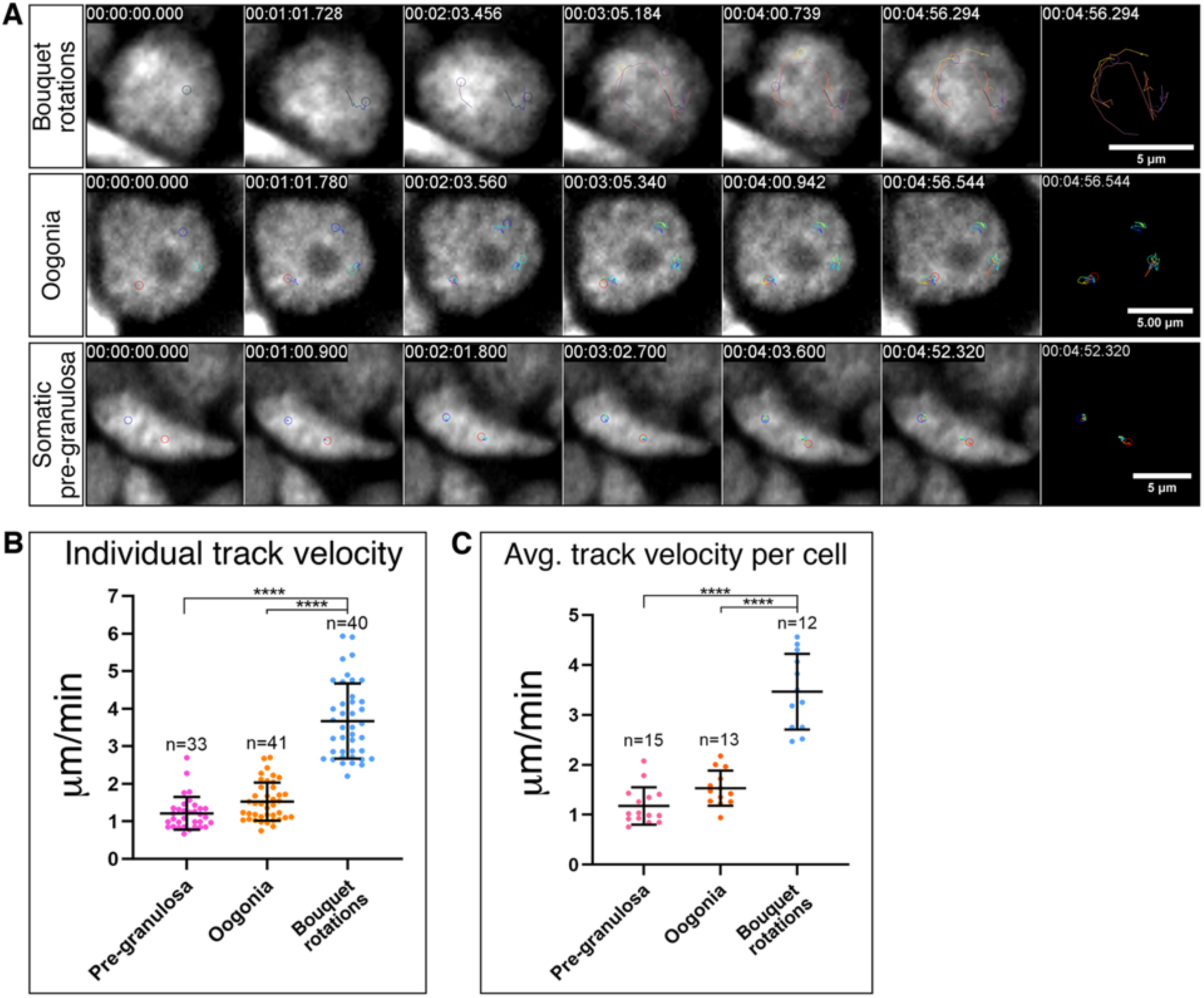
Live imaging analysis of bouquet chromosomal rotations. **A.** Live time-lapse imaging of *Tg(h2a:h2a-gfp)* (grey) ovaries, recording chromosomal dynamics during bouquet rotations, and their absence in control oogonia and somatic pre-granulosa follicle cells. Sum of chromosomal tracks at *T_f_* are shown for each cell. Images are snapshots from Movie S1. Scale bars are 5 μm. n=12 bouquet rotation oocytes, 13 oogonia, and 15 follicle cells, from 4 ovaries. **B-C.** Individual track velocity (B) and average track velocity per cell (C) are plotted from a representative experiment. n=number of tracks (B) and cells (C).

**Supplementary Figure 3.**
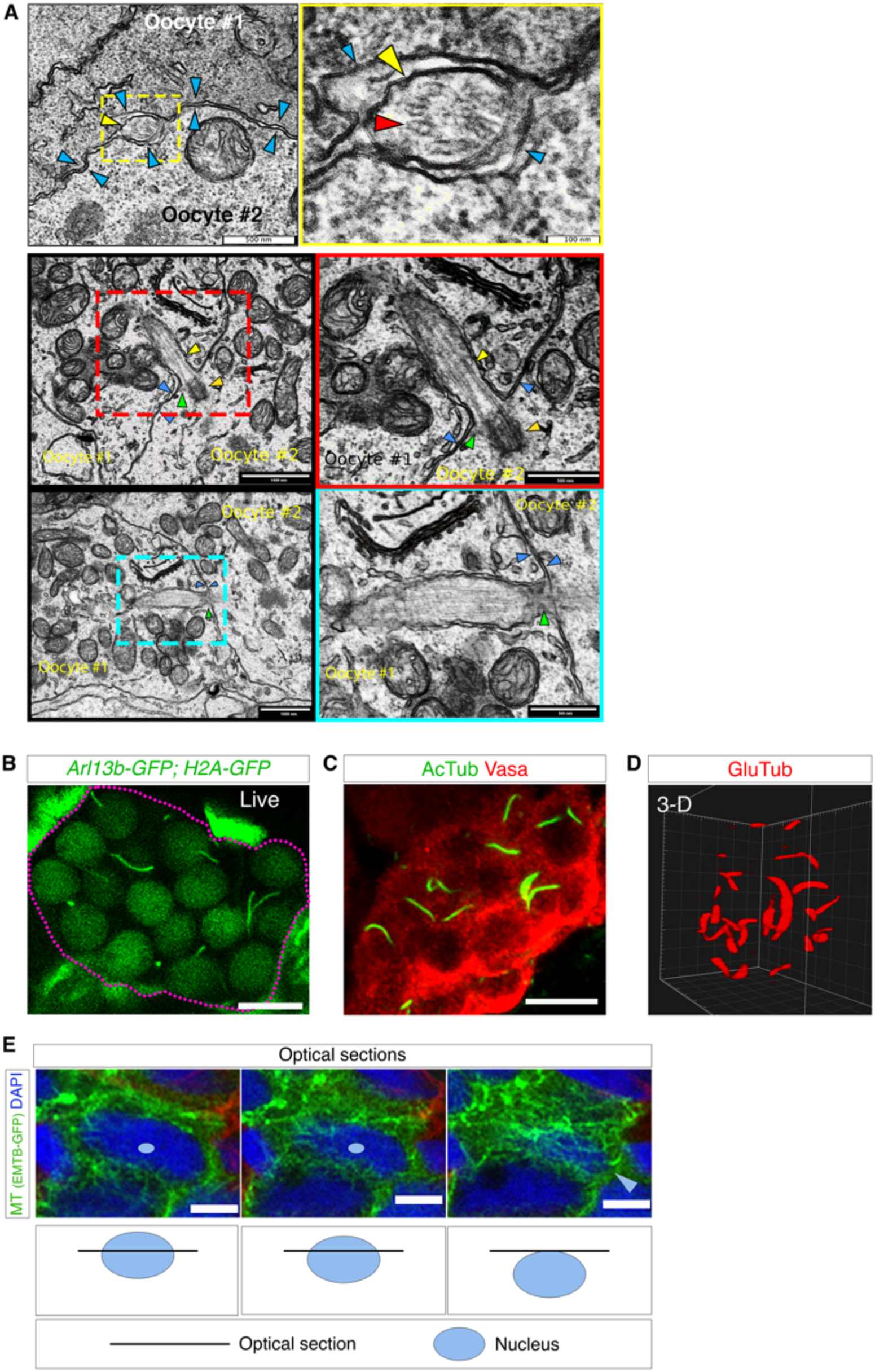
**A.** Additional examples of TEM images of the zygotene cilium. A cross section of a cilium found between two oocytes is shown in the top pair panels, and longitudinal sections of two cilia that emanate from oocytes are shown in the middle and bottom pair panels. The right panels are magnifications of the color-coded boxed regions in the corresponding left panels. Arrowheads indicate: oocyte membrane (light blue), axoneme (yellow), MT doublets (red), basal body (orange), transition zone (green). Scale bars are indicated. n=23 oocytes in 5 ovaries. **B.** Detection of the zygotene cilia in live *Tg(bact:arl13b-gfp); Tg(h2a:h2a- gfp)* ovaries. Magenta dashed line outlines the cyst. Image is a snapshot from Movie S11. n=9 ovaries. Scale bar is 10 μm. **C.** Sum-projection of AcTub and the germ cell marker Vasa labeling in the germline cyst. n=5 ovaries. Scale bar is 10 μm. **D.** 3-D reconstruction of gluTub labeling in a zygotene cyst. n=7 ovaries. Grid is 1 μm. **E.** Microtubule labeling [*Tg(βact:EMTB-3XGFP)*] in zygotene bouquet oocytes show perinuclear cables (nucleus is counterstained with DAPI). Optical sections from different Z positions, shown from left to right (indicated by the boxes below each panel), reveal perinuclear microtubule cables that surround the nucleus. The nucleus of interested is indicated with a light blue circle in the two left panels, and an arrowhead indicate its X,Y location in the new Z position. n= 7 ovaries.

**Supplementary Figure 4.**
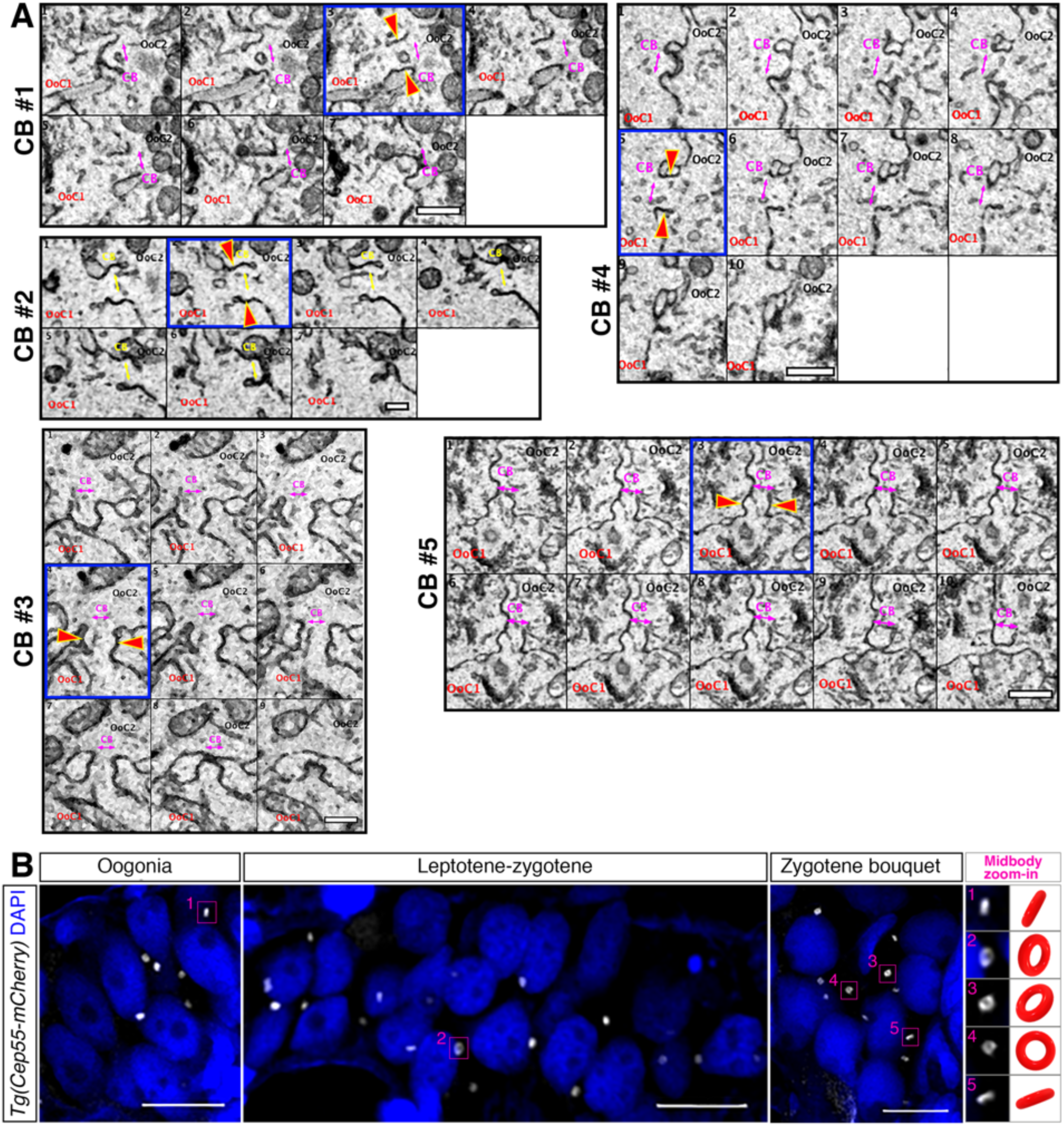
Identification of cytoplasmic bridges (CBs) in the germline cyst. **A.** Montage images through z-sections from SBF-SEM data showing several examples of CBs. In all panels, the two oocytes connected by the CB are marked as “Ooc1”, “Ooc2” and the CB opening is labeled “CB” and with magenta arrows. The blue framed panels mark a section through approximately the center of the CB, in which red arrows indicate the CB cytoplasmic membranes. CBs are short and often exhibit a typical ruffled membrane shape. The CB cytoplasm is indistinguishable from the oocyte cytoplasm and often contain vesicles (see for example CB #1 and #4). Scale bars are 500 nm, except CB #2 =200 nm. **B.** CBs are marked by a transgenic midbody marker (*Tg(bact:Cep55-mCherry)*, grey), found between oocyte (nuclei are shown by DAPI, blue), in oogonia, leptotene-zygotene, and zygotene bouquet cysts. Cep55-mCherry shows a typical ring shape of the midbody as shown in the numbered zoom-in images on the right that correspond to the numbered boxes in the images. The cartoon ring in each panel shows the orientation of the Cep55-mCherry ring. Scale bars are 10 μm.

**Supplementary Figure 5.**
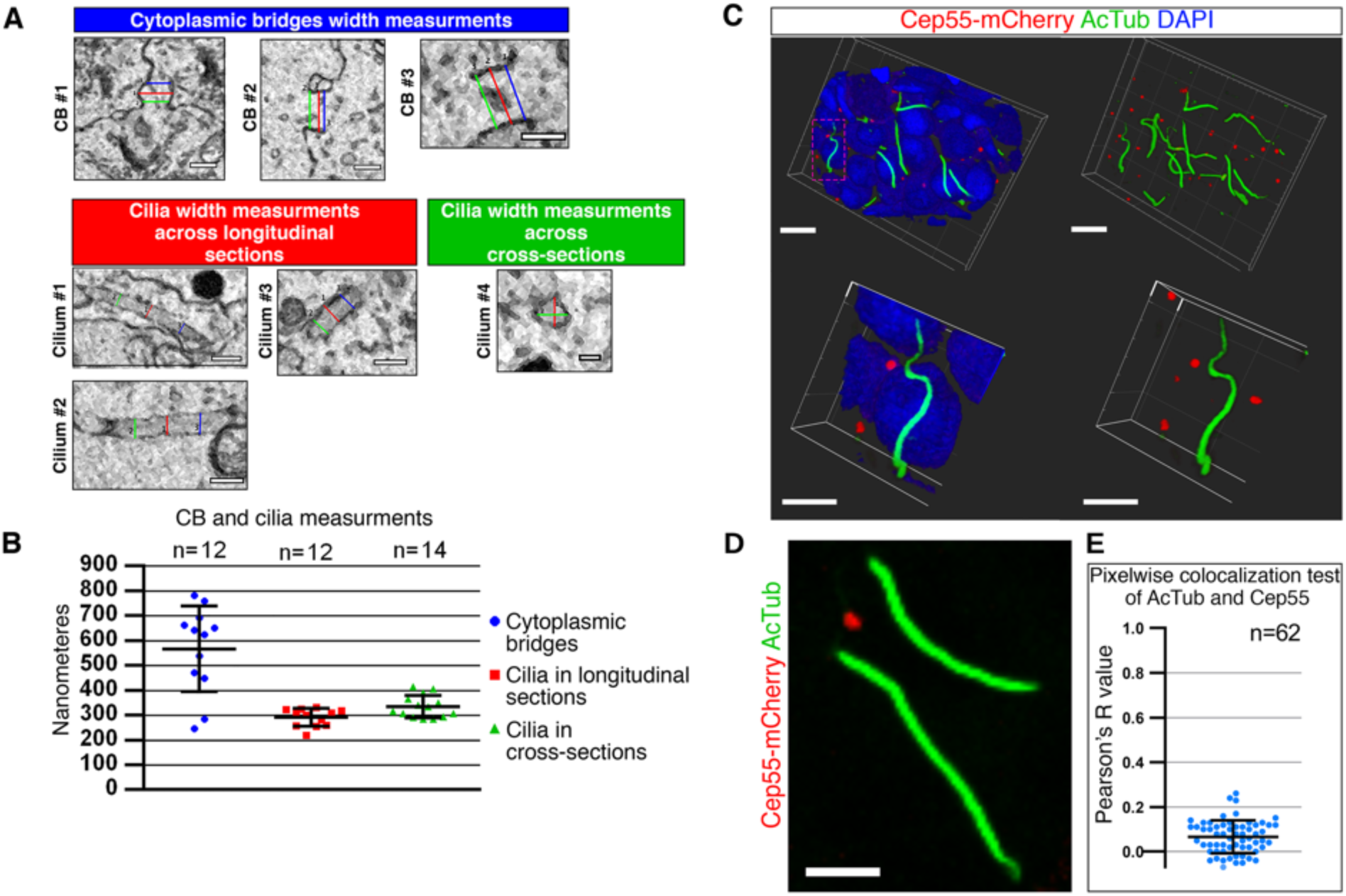
The zygotene cilia are distinct from CBs. **A.** Measurement of cilia and CBs from the SBF-SEM data. CB opening diameter (width) was measured by three cross sections (red, green, blue) along the length of the CB. The average of the three lines was determined as the width of the CB. Three examples are shown. Cilia were measured in two ways. 1. Across ciliary longitudinal sections (left), three cross sectioning lines (blue, red, green) along the length of the cilia were measured and their average was determined as the cilia width (291 ± 36 nm). 2. Across ciliary cross sections, two perpendicular lines along the diameter of the cross section (red, green) were measured and their average was determined as the ciliary width (324 ± 59 nm). Scale bars are 500 nm, except Cilium #4 – 200nm. **B.** Measurements of cilia and CBs from A are plotted (CBs are 567 ±172 nm wide). Bars are mean ± SD. n = number of CBs and cilia measured in each method. **C.** 3-D images from ovaries labeled with AcTub (green), and transgenic Cep55-mCherry (red), with (left) or without (right) DAPI. Bottom images are zoom-in from the magenta boxed region in the top left image, with (left) or without (right) DAPI. n=6 ovaries. Scale bars in top and bottom images are 10 μm and 5 μm, respectively. **D-E.** Pixelwise colocalization test of AcTub and Cep55 shows no or random co-localization. n=number of cilia. **D** is a representative ROI used in E. Scale bar is 5 μm.

**Supplementary Figure 6.**
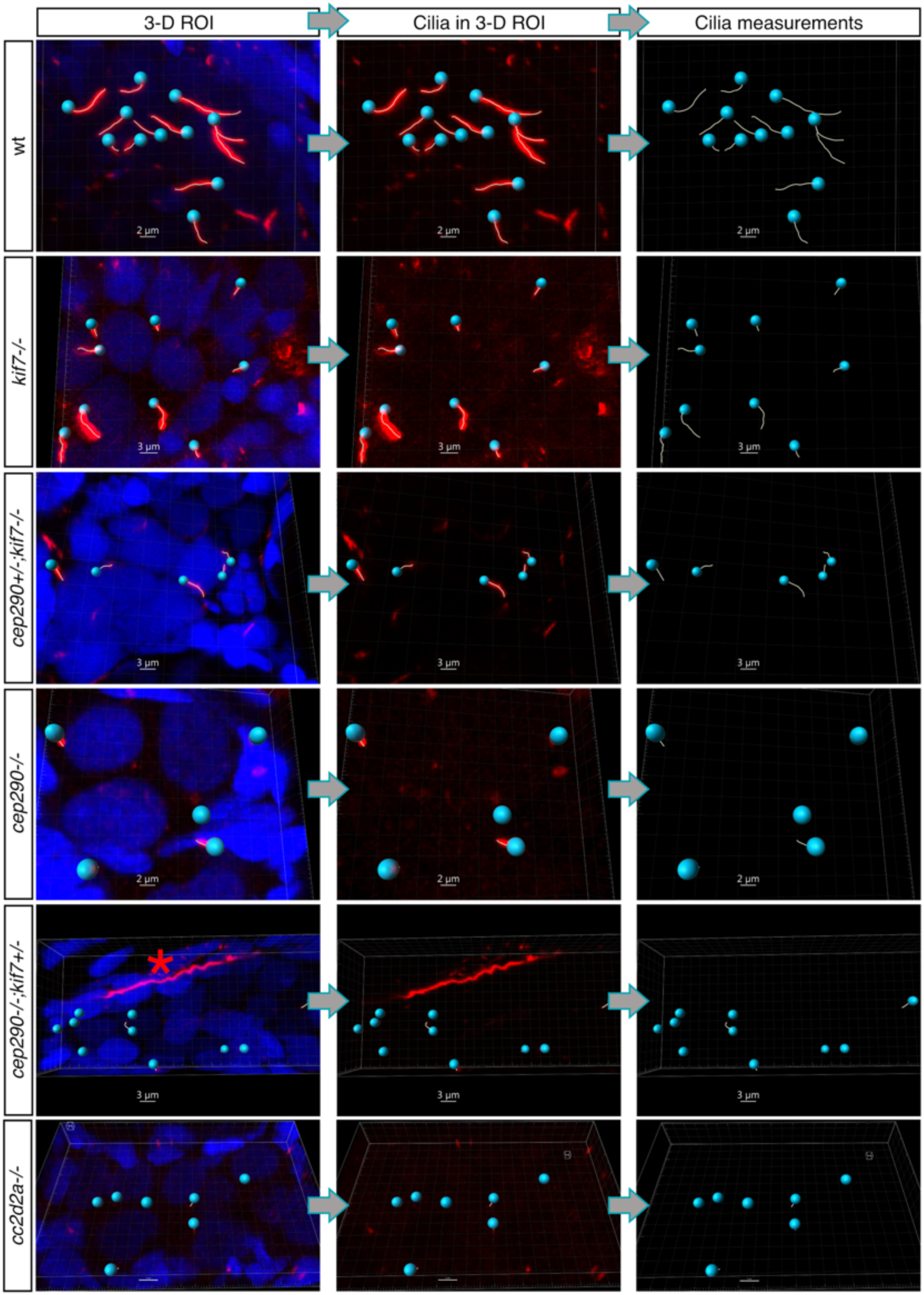
Measurement of ciliary length. Ovaries labeled for AcTub (red), DiOC6 (green, used for visualizing oocyte cytoplasm, not shown), and DAPI (blue) were used for analysis. In entire cyst 3D images (left panels), cilia were tracked and measured (cyan balls and lines, middle panels), as described in the Methods section. Measured lengths (right panels) were exported for statistical analysis. Representative images of analyzed genotypes are shown. Red Asterix in the left bottom panel indicate a nearby AcTub-labeled ovarian axon. Scale bars are indicated.

**Supplementary Figure 7.**
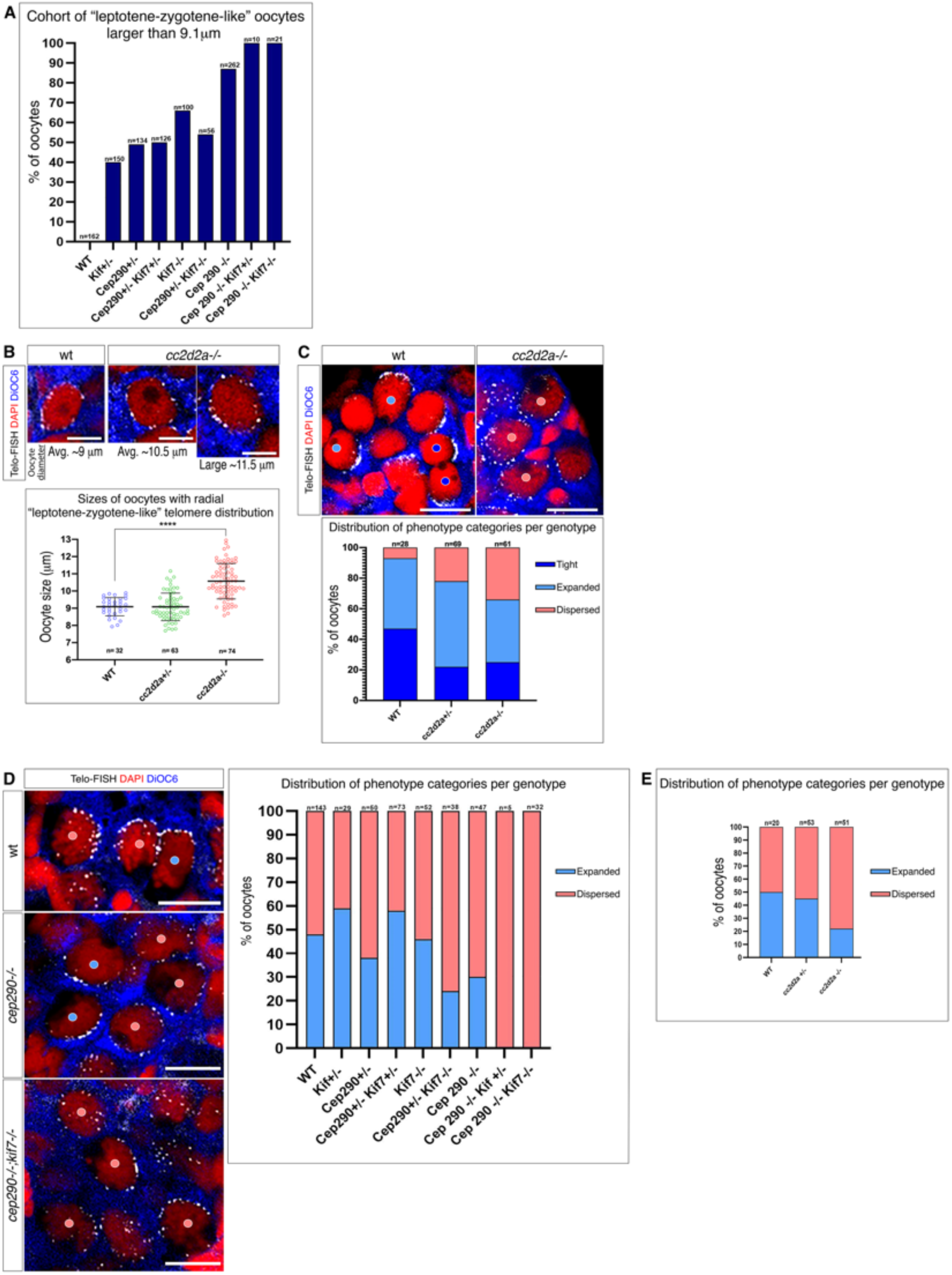
Supporting information for Figure 2. **A.** The percentage in each genotype of leptotene-zygotene-like oocytes that were larger than 9.1 μm (the largest wt leptotene-zygotene oocyte size). n=total number of leptotene-zygotene-like oocytes. **B.** Representative images of “leptotene-zygotene-like” oocytes of average sizes in wt (n=3 ovaries), *cc2d2a^+/-^* (n=3 ovaries), and *cc2d2a^-/-^* (n=5 ovaries). Large “leptotene-zygotene-like” *cc2d2a^-/-^* oocytes are shown. Scale bars are 5 μm. Bottom panel: sizes of “Leptotene-like“ oocyte per genotype. n=number of oocytes. Bars are mean ± SD. **C.** Representative images of mid-bouquet stage oocytes with Tight (blue dot), Expanded (light blue dot), and Dispersed telomeres (red dots; categories as in Fig. 2G). Scale bars are 10 μm. Bottom panel: the percentage of each category in wt (n=3 ovaries), *cc2d2a^+/-^* (n=3 ovaries), and *cc2d2a^-/-^* (n=5 ovaries). n=number of oocytes. **D.** Late bouquet analysis, showing representative images (left panels) of oocytes with Expanded (light blue dot), and Dispersed telomeres (red dots; categories as in Fig. 2G), and their distribution per genotype (right panel). n=number of oocytes. Scale bar is 10 μm. **E.** Same analysis as in **D**, showing the category distribution in wt, *cc2d2a^+/-^* and *cc2d2a^-/-^* ovaries. n=number of oocytes.

**Supplementary Figure 8.**
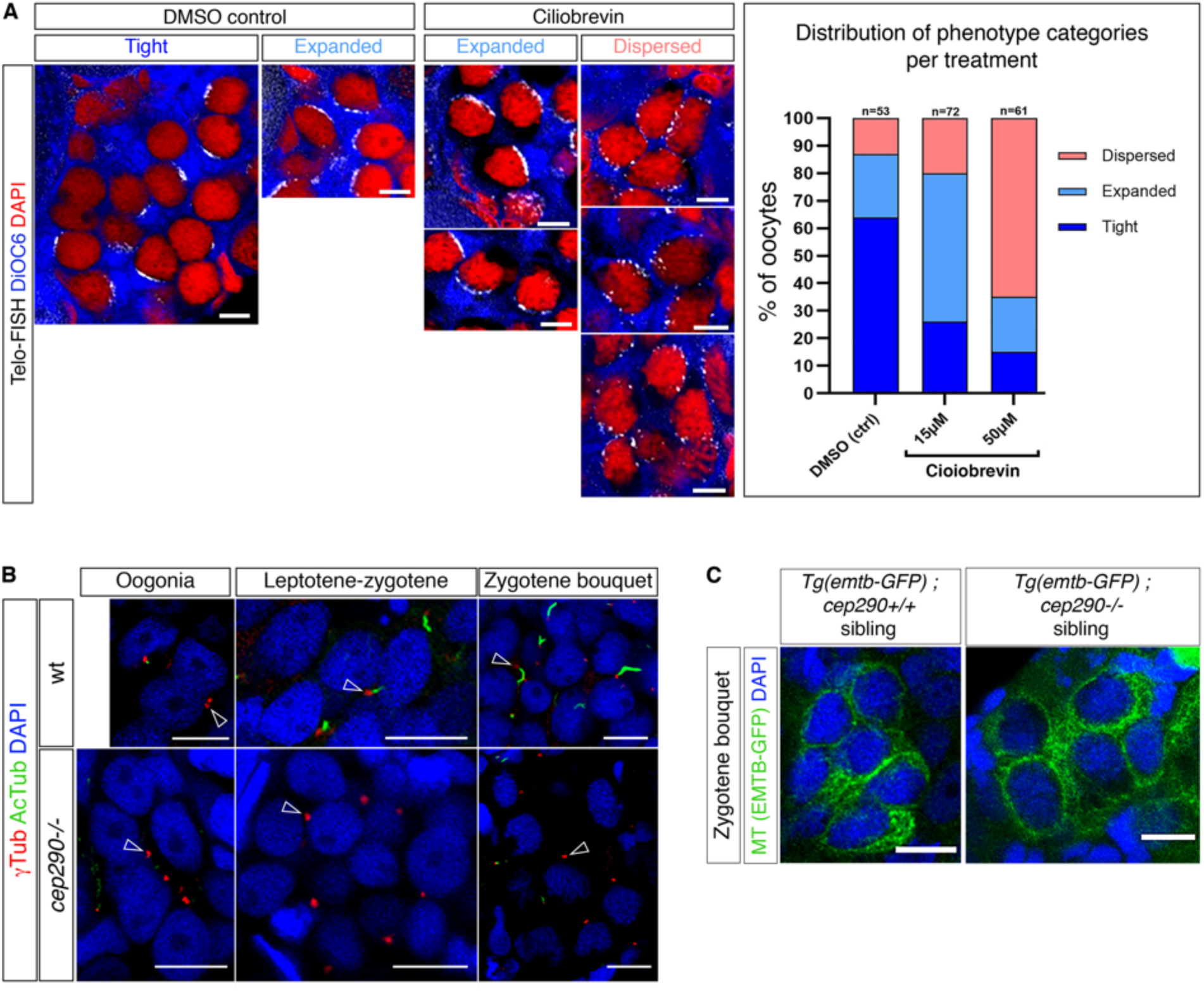
Ciliary loss phenocopies bouquet perturbation by dynein inhibition, and centrosome MTOC functions are normal in *cep290^-/-^* ovaries. **A.** Ovaries were treated with DMSO control or the dynein inhibitor ciliobrevin (15μM or 50 μM, for 80 minutes), and then fixed for analysis. Bouquet analysis was performed as in Fig. 2G. Representative images of mid-bouquet stage oocytes with Tight, Expanded, and Dispersed telomeres (categories as in Fig. 2G) are shown in the left panels. The distribution of categories per treatment is plotted in the right panel. n=number of oocytes. 3-5 ovaries were analyzed per treatment. Scale bars are 5 μm. **B.** Ovaries labeled for AcTub (green), γTub (red), and DAPI (blue), showing normal localization of γTub to centrosomes (white arrows) in both wt and *cep290^-/-^* oogonial, leptotene-zygotene, and zygotene bouquet germline cysts. Note the absence of cilia in *cep290^-/-^* cysts. Scale bars are 10 μm. **C.** *cep290^+/-^* fish were crossed to *Tg(bact:emtb-3gfp)* fish and in-crossed to produce wt, and *cep290^-/-^* fish that express transgenic *emtb-3gfp* to label MTs. Ovaries of siblings were labeled for GFP (MT, green) and DAPI, showing normal MT organization in zygotene bouquet cysts in *cep290^-/-^* (n=7 ovaries), similar to wt (n=7 ovaries) ovaries. Scale bars are 10 μm.

**Supplementary Figure 9.**
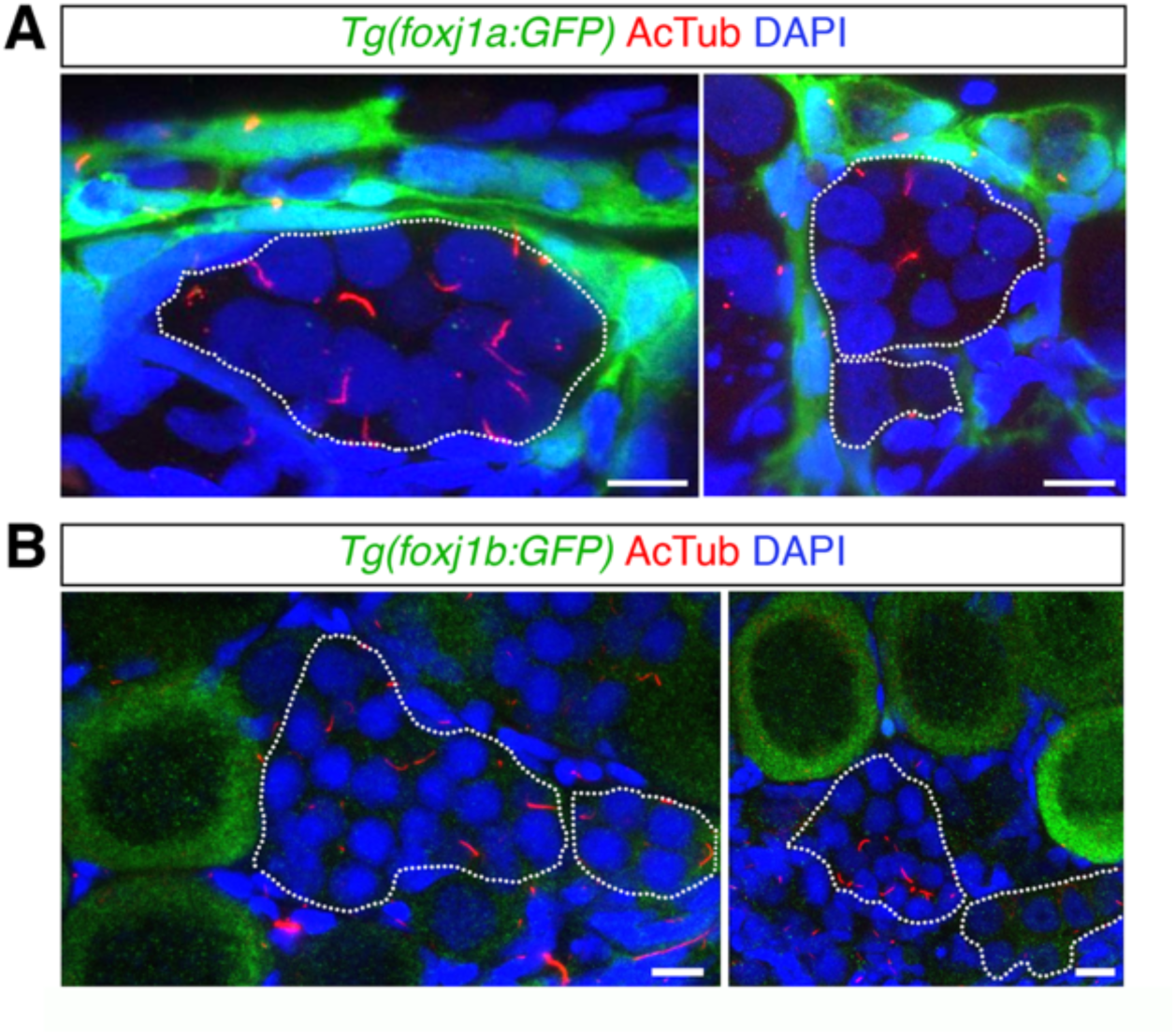
*foxj1a* and *foxj1b* reporter line expression is not detected in the prophase stage germline cyst (white outline), where the zygotene cilium forms (AcTub), but only in later stage oocytes and somatic cells. n=6 ovaries for *foxj1a* and 4 ovaries for *foxj1b*. Scale bar is 10 μm.

**Supplementary Figure 10.**
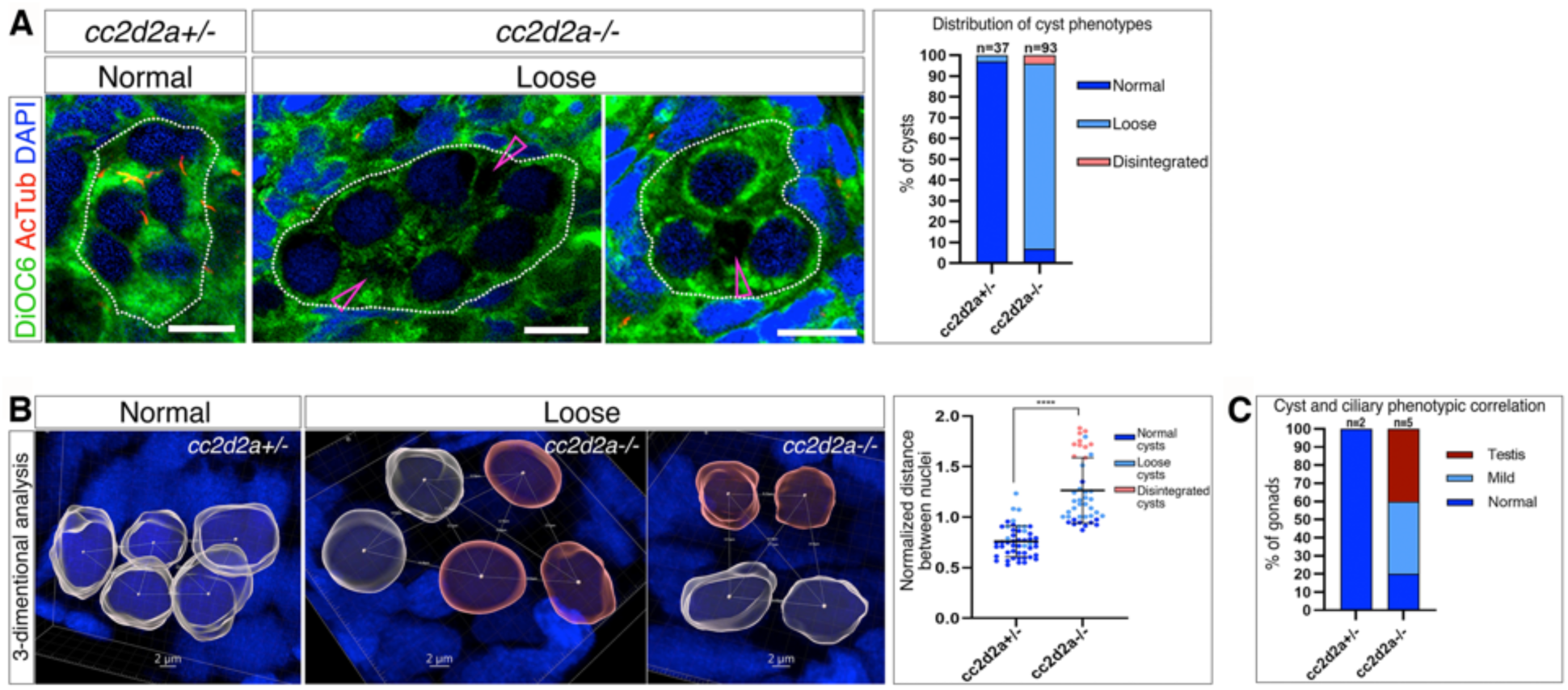
Germline cyst disintegration in *cc2d2a* mutant ovaries. **A.** Representative images of cyst (white outlines) phenotype categories in *cc2d2a^+/-^* (n=2 ovaries), and *cc2d2a^-/-^* (n=5 ovaries) ovaries labeled for the cytoplasmic marker DiOC6 and AcTub, showing gaps between oocytes (magenta arrows). Scale bars are 10 μm. Right panel: the percentage of cyst phenotype categories in all genotypes (n=number of cysts). **B.** Representative images of 3-D cyst morphology analysis per category, showing the distances between neighboring nuclei. Normalized distances are plotted in the right panel. n=7 cysts from 2 ovaries per genotype. Bars are mean ± SD. **C.** Severity of cyst phenotypes and gonad conversions plotted for *cc2d2a^+/-^* and *cc2d2a^-/-^*. n=number of gonads.

**Supplementary Figure 11.**
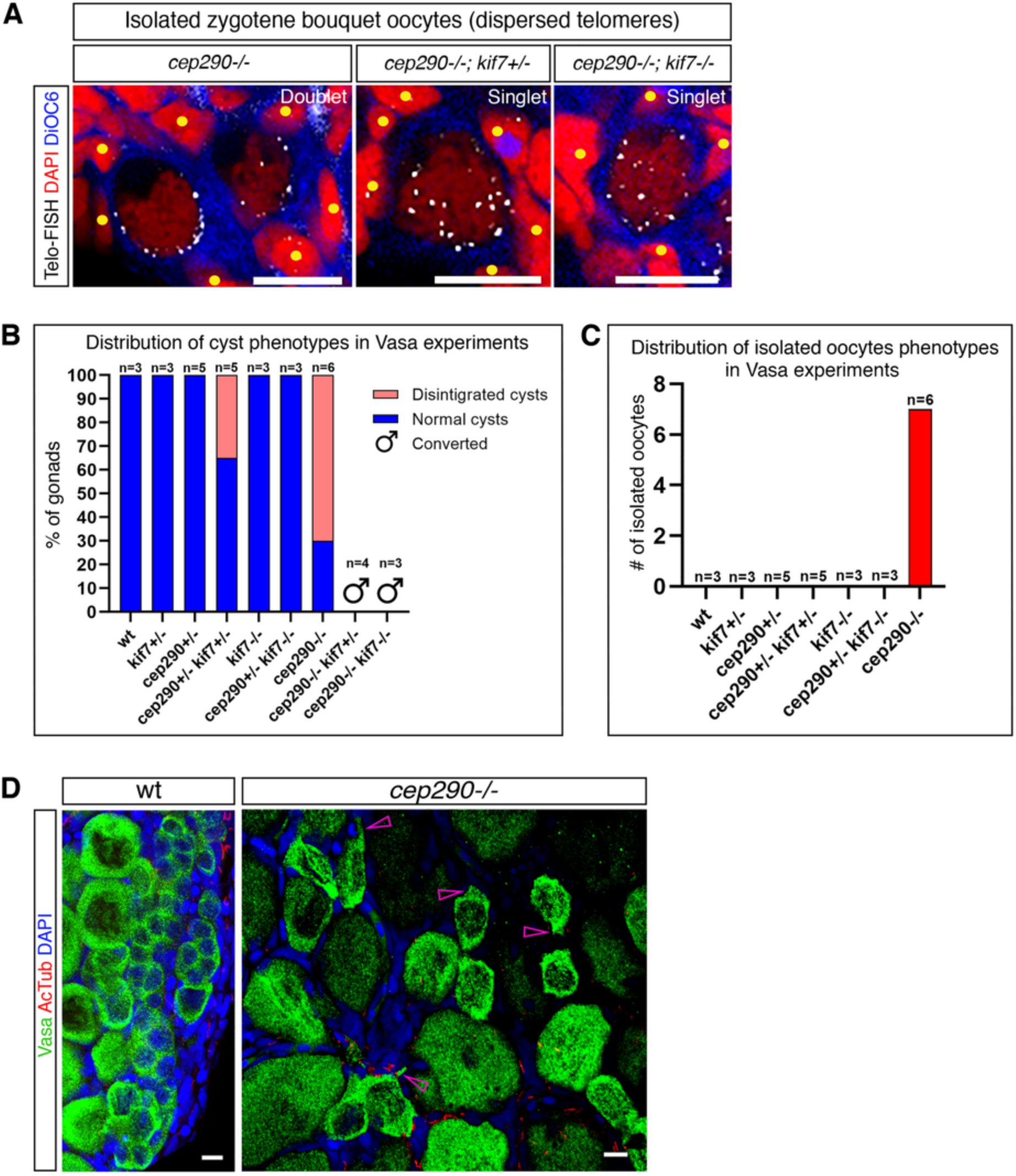
Supporting information for Figure 3. **A.** Isolated oocytes are confirmed as zygotene bouquet stage oocytes by size (according to DiOC6 cytoplasmic labeling, blue), and by telomere labeling (Telo-FISH, greyscale), which shows dispersed telomere on the NE. DAPI labeling (red) show follicle cell nuclei (yellow circles) that surround the isolated oocyte, confirming that it is not part of a cyst. We identified 43 isolated oocytes in 7 ovaries of *cep290^-/-^*, *cep290^-/-^;kif7^+/-^* and *cep290^-/-^;kif7^-/-^*, and 0 isolated oocytes in 10 wt ovaries. Scale bars are 5 μm. **B.** The percentage of cyst phototype categories in all genotypes from Vasa experiments in Fig. 4C. n=number of ovaries. *cep290^-/-^;kif7^+/-^* and *cep290^-/-^;kif7^-/-^* ovaries were converted to testes (discussed in Fig. 5B, and S12). **C.** The number of isolated oocytes detected per genotype in the same experiments. n=number of ovaries. **D.** Vasa labeling shows later oocytes with irregular mesenchymal like morphology with pointy edges (pink arrows), in *cep290^-/-^* ovaries, in addition to cyst phenotypes shown in Fig. 4C. n=6 ovaries. Scale bars are 10 μm.

**Supplementary Figure 12.**
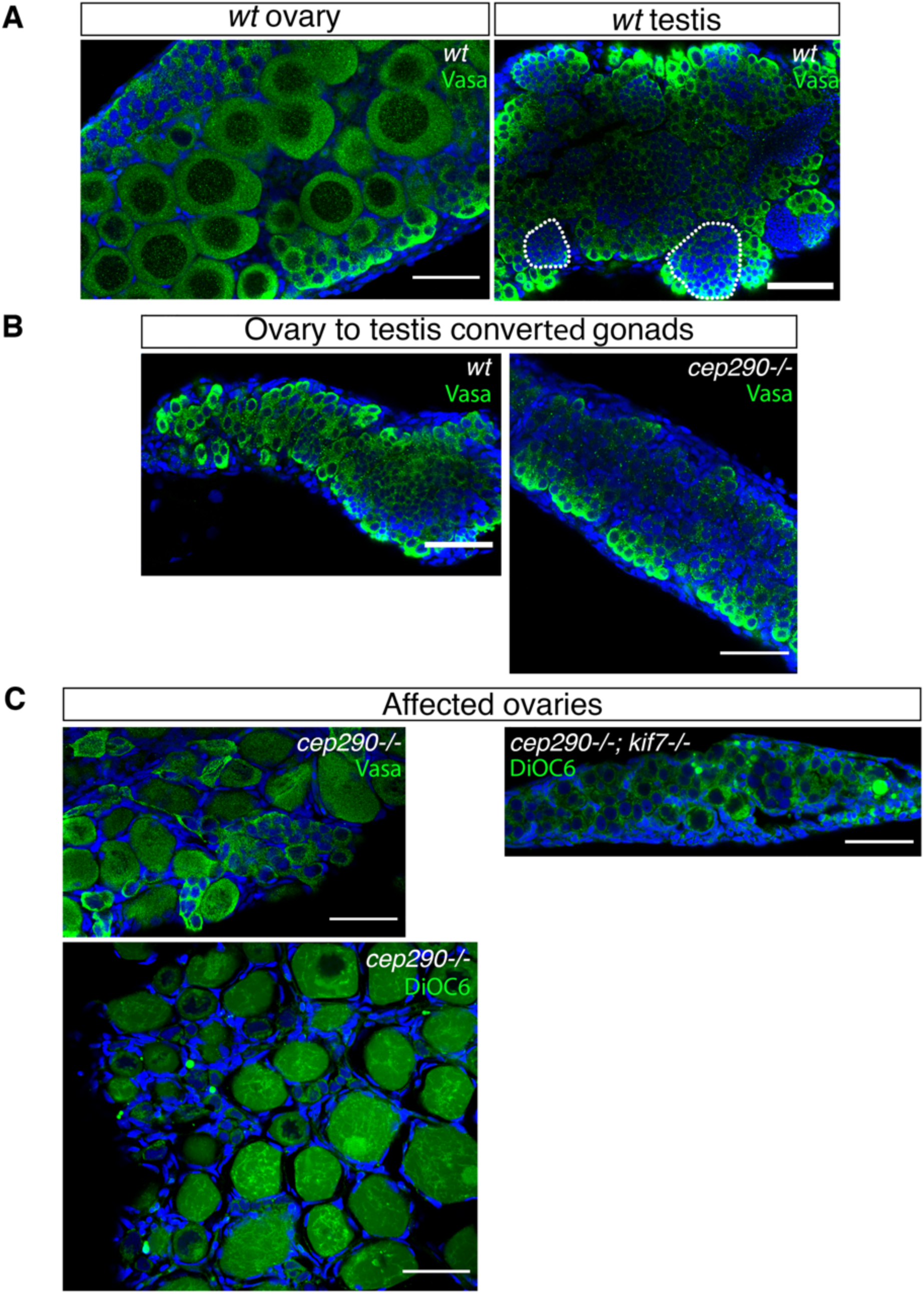
Examples of normal, converted and affected gonads. **A.** Normal *wt* ovary (left) and testis (right). In normal developing ovaries, oocytes of varying stages and sizes are clearly detected. In developing testes, spermatogonial cells are dense throughout the gonad, and their organization into early seminiferous tubules (white outline) is apparent. **B.** Ovaries that converted to testes are much thinner than normal developing ovaries and are similar to very young testes. They do not contain oocytes, and only spermatogonial-like cells with still no apparent tubule organization. *Wt* (left) and *cep290^-/-^* (right) converted gonads are shown. **C.** Affected ovaries clearly contain oocytes of varying stages and sizes, and not spermatogonial-like cells. These ovaries can be smaller and less organized in general, and exhibit the phenotypes described in this work. Gonads in **A-C** are representative for each category from various experiments in this work, and labeled with DAPI and either Vasa or DiOC6 as indicated. All gonads are shown to scale. Scale bars are 50 μm.

**Supplementary Figure 13.**
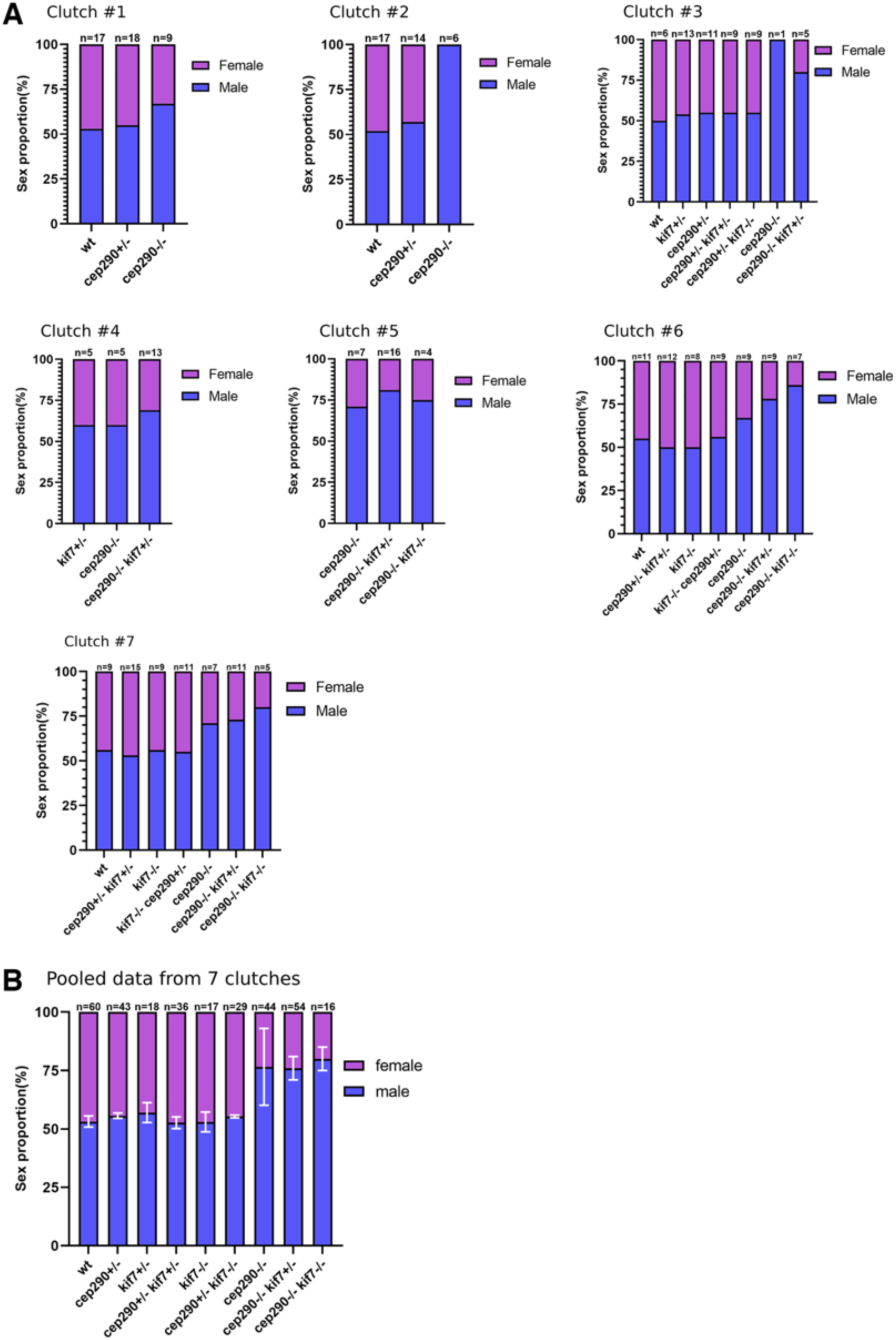
Adult male bias in mutants. **A.** Sex ratios of adult fish from five different clutches. n=number of fish. *cep290^+/-^;kif7^-/-^* and *cep290^-/-^;kif7^-/-^* fish are produced in mendelian ratios of 12.5% and 6.25% respectively, are used for experiments at juvenile stages, and mostly do not survive to adulthood. We were able to raise these in a few clutches for analysis. **B.** Sex ratios of adult fish pooled from all clutches in A. n=number of fish. White bars are SD.

**Supplementary Figure 14.**
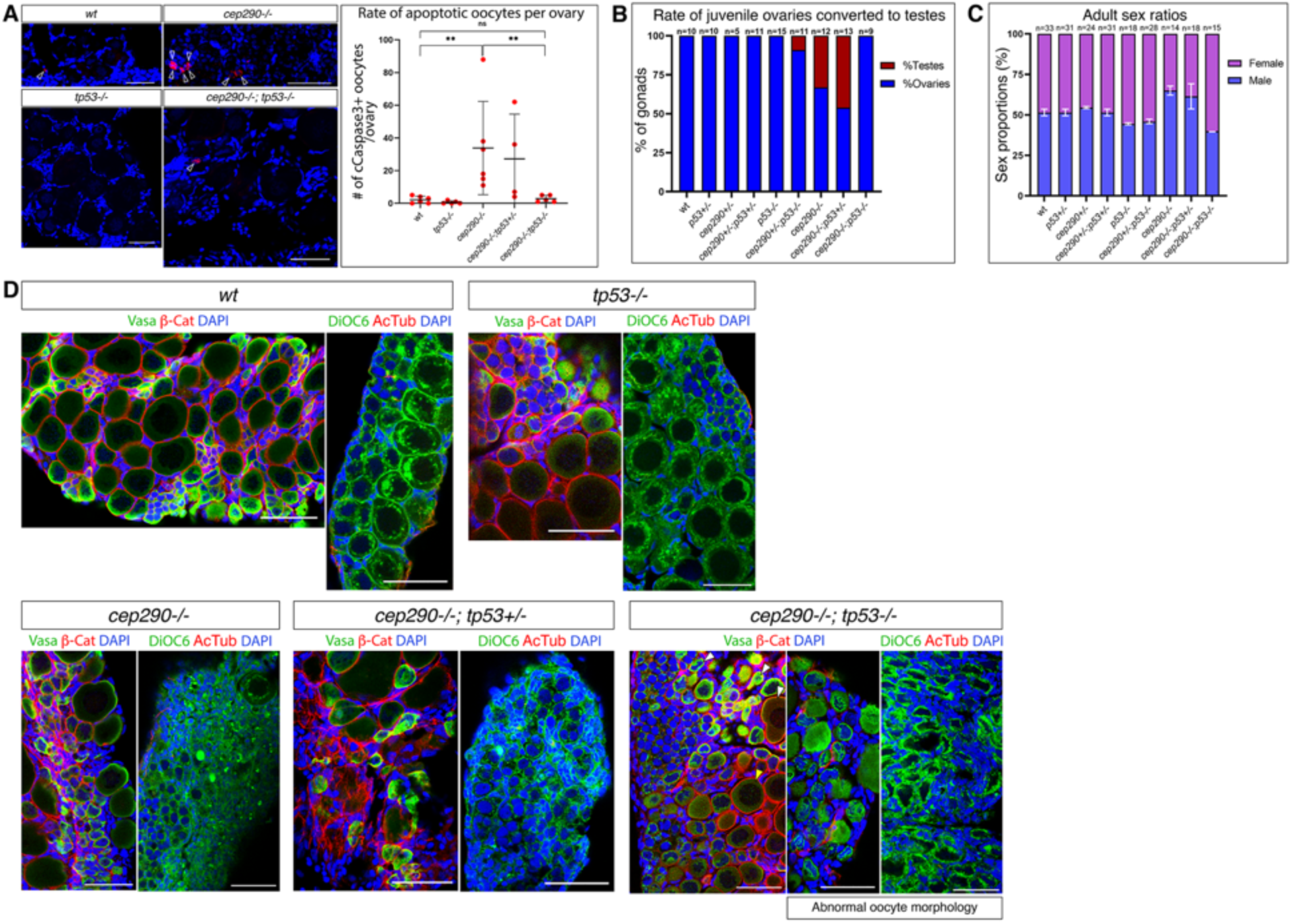
Loss of *tp53* rescues *cep290* mutant oocyte apoptosis, juvenile gonad conversion, and adult sex bias. **A.** cCaspase3 apoptosis labeling (red, white arrowheads) in juvenile ovaries. Left panels: representative images of wt, *tp53^-/-^*, *cep290^-/-^* and *cep290^-/-^;tp53^-/-^* ovaries. Scale bars are 50 μm. Right panel: number of cCaspase3-positive oocytes per gonad for all genotypes. Each dot represents a gonad, n=4-6 gonads per genotype. Bars are mean ± SD. **B.** Rates of juvenile gonad conversion per genotype. n=number of gonads. Example images of normal and converted gonads are shown in Fig. S12. **C.** Sex ratios of adult fish. n=number of fish. White bars are SD. **D.** Representative images of ovaries from *wt* and *cep290;tp53* genotypes labeled either for Vasa (germ cells, green), β-cat (cytoplasmic membrane, red), and DAPI (blue) in left panels, or DiOC6 (cytoplasm, green), AcTub (cilia, red), and DAPI (blue) in right panels. *wt* and *tp53^-/-^* ovaries appear normal and contain normal cysts and typically spherical oocytes. *cep290^-/-^* and *cep290^-/-^;tp53^+/-^* ovaries exhibit typical *cep290^-/-^* phenotypes with abnormal cysts and few spherical later stage oocytes. In contrast, *cep290^-/-^;tp53^-/-^* exhibit spherical oocytes (yellow arrowhead), but also many oocytes with defected abnormal morphology (white arrowheads and two right panels). These ovaries are not converted (B-D) and demonstrate that oocytes survive apoptosis, but fail to develop normally. n=5-7 ovaries in each genotype. Scale bars are 50 μm.

**Supplementary Figure 15.**
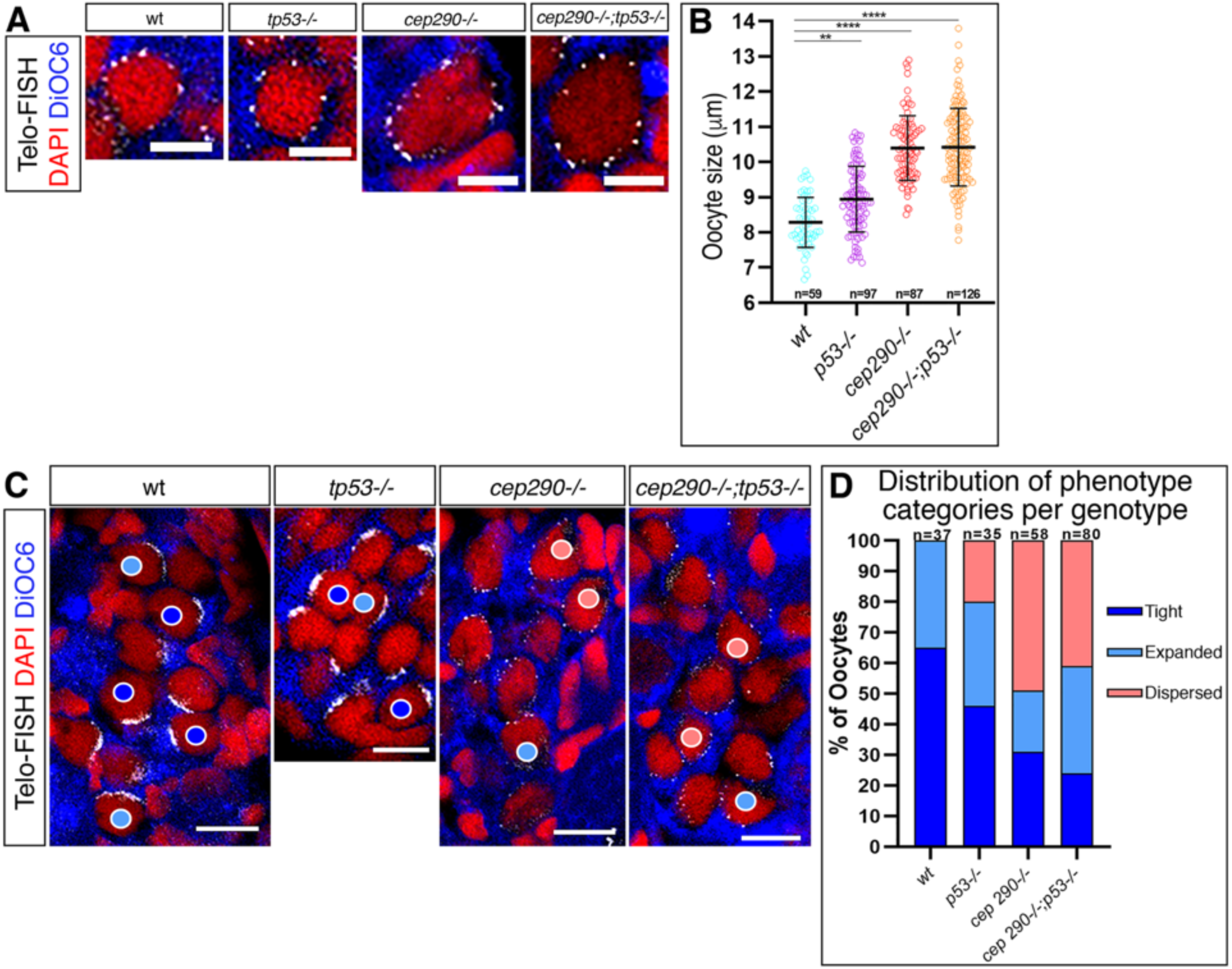
*cep290* ciliary loss bouquet defects are not rescued by loss of *tp53* in *cep290^-/-^;tp53^-/-^* ovaries. **A.** Representative images of “leptotene-zygotene-like” oocytes in *wt* and *cep290;tp53* genotypes. Scale bars are 5 μm. **B.** “Leptotene-zygotene-like” oocyte sizes per genotype. n=number of oocytes. Bars are mean ± SD. **C.** Representative images of mid- bouquet stage oocytes with Tight (dark blue circles), Expanded (light blue circles), and Dispersed (red circles) telomeres (categories shown in Fig. 2G). Scale bars are 10 μm. D. The percentage of each category in all genotypes. n=number of oocytes. *Chi-square* test *p*-values are <0.0001 between wt and all genotypes.

**Supplementary Figure 16.**
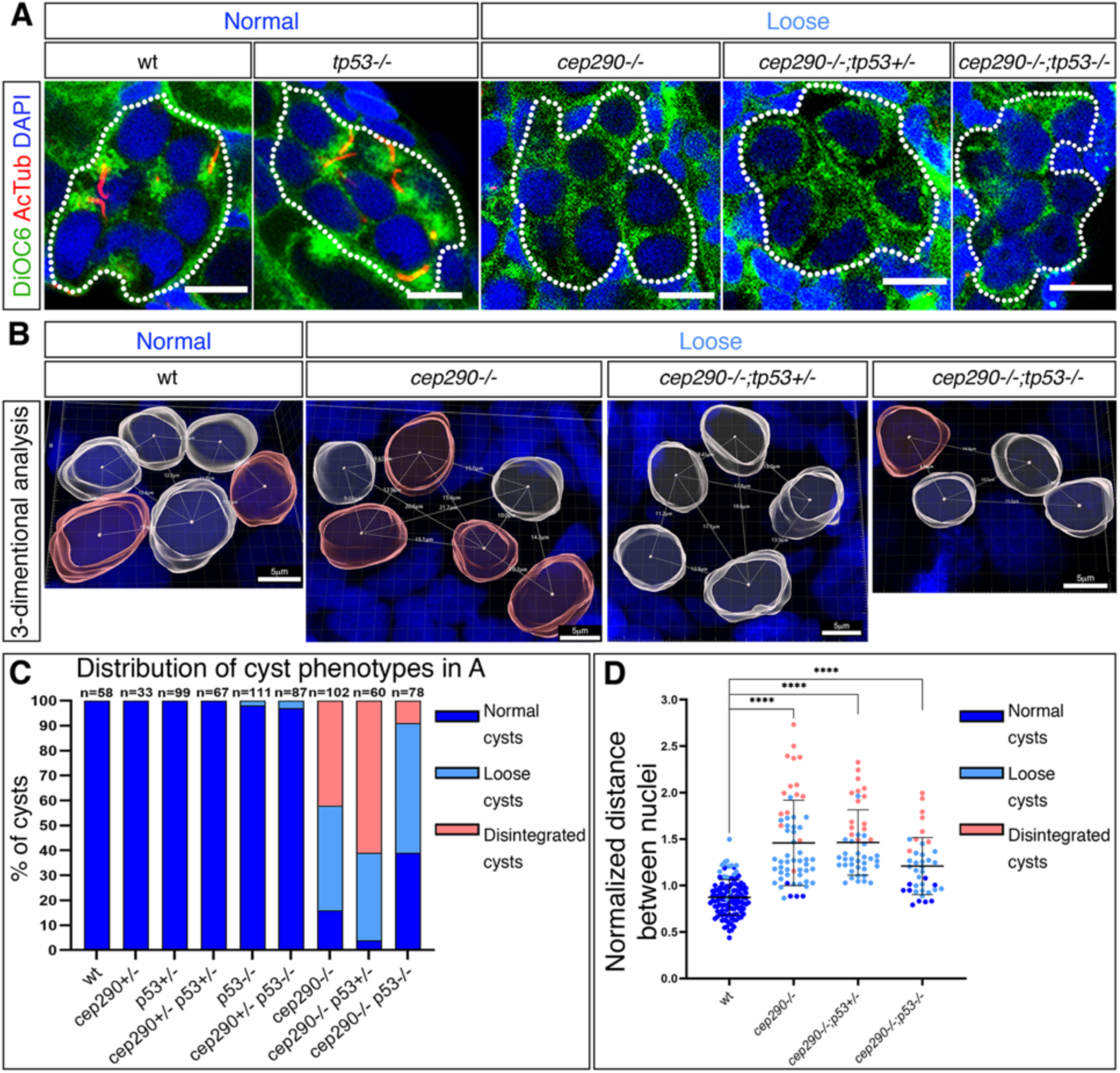
*cep290* ciliary loss cyst defects are not rescued by loss of *tp53* in *cep290^-/-^;tp53^-/-^* ovaries. **A.** Representative images of “normal” and “loose” cyst (white outlines) in ovaries from in *wt* and *cep290;tp53* genotypes labeled for the cytoplasmic marker DiOC6 and AcTub, as in Fig. 4A. Scale bars are 10 μm. The percentage of cyst phenotype categories in all genotypes is shown in **C** (n=number of cysts; *Chi-square* test *p*-values are <0.0001 between wt and *cep290^-/-^*, *cep290^-/-^;tp53^+/-^*, and *cep290^-/-^;tp53^-/-^*, and n.s for all other genotypes). **B.** Representative images of 3-D cyst morphology analysis per category, as in Fig. 4B, showing the distances between neighboring nuclei. Normalized distances are plotted in **D**. n=5-8 cysts from 3 ovaries per genotype. Bars are mean ± SD.

**Supplementary Figure 17.**
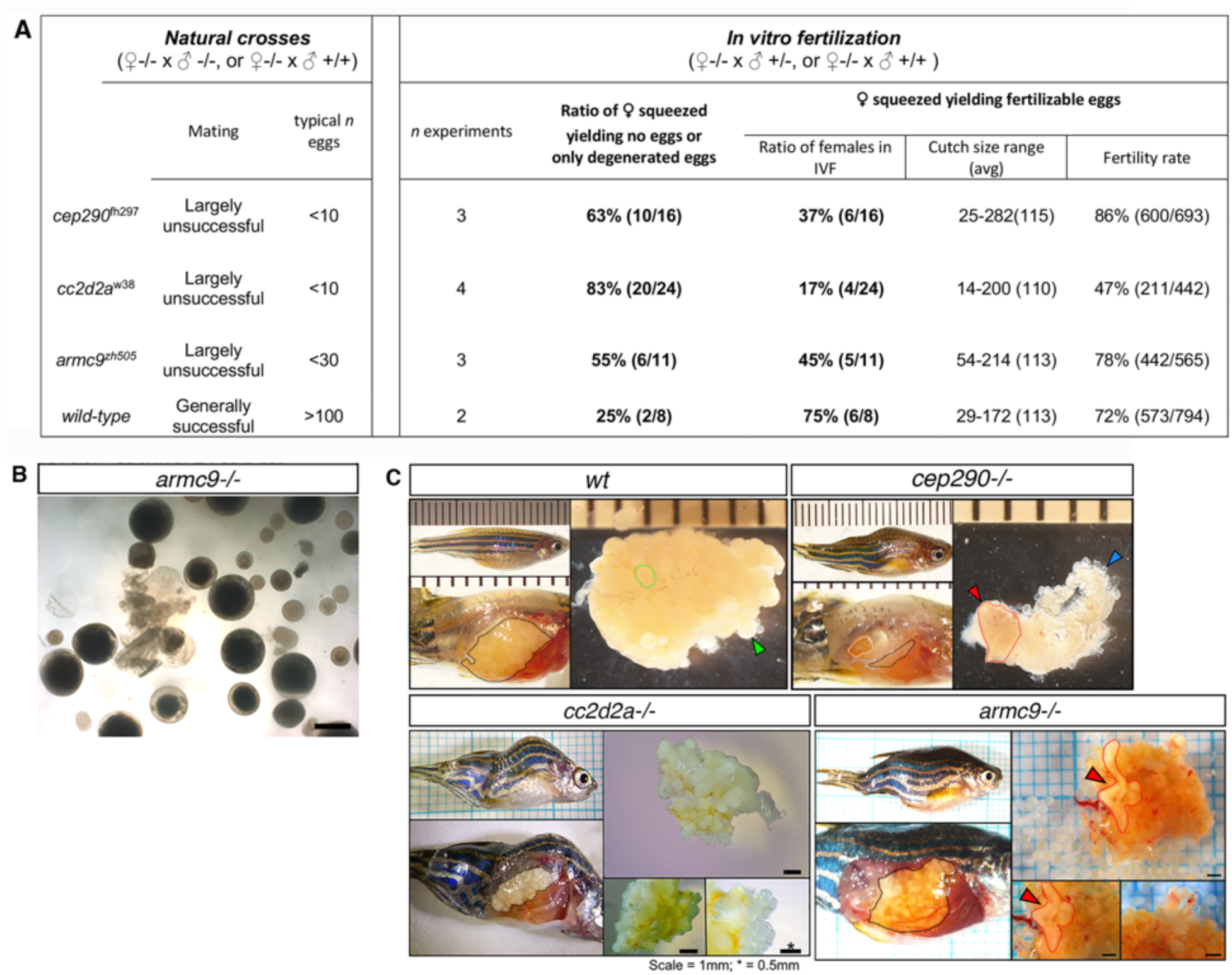
Fertility deficiencies in multiple ciliary mutants. **A.** A table summarizing fertility assessment of various ciliopathy mutant zebrafish models. In natural crosses, mutants were virtually all infertile. In *in vitro* fertilization (IVF) experiments between wt females and wt males, the majority of *wt* females yielded healthy looking fertilizable eggs (Fig. 5C). In contrast, in IVF experiments between mutant females and wt or heterozygous males, most mutant females yielded no eggs upon squeezing, or only degenerated eggs (Fig. 5C). In the fewer cases where healthy looking eggs could be collected, the fertility rate was overall similar to wt. n=number, avg=average, NA=not available. **B.** A representative image of eggs obtained by squeezing for IVF experiments from *armc9^-/-^* females, as in Fig. 5C. Scale bar=1mm. **C.** Additional examples of wt, *cep290^-/-^*, and *cc2d2a^-/-^* adult females to Fig. 5D, as well as a similar image of *armc9^-/-^* female, showing the ovaries within the peritoneal cavity (left panels), and dissected ovaries (right panels). Mutant ovaries are smaller and contain degenerated tissue (red outline and arrowhead). Ruler marks and scale bars are 1 mm (except when indicated by *, 0.5 mm). Mutant females exhibit scoliosis, a typical ciliary phenotype.

**Supplementary Figure 18.**
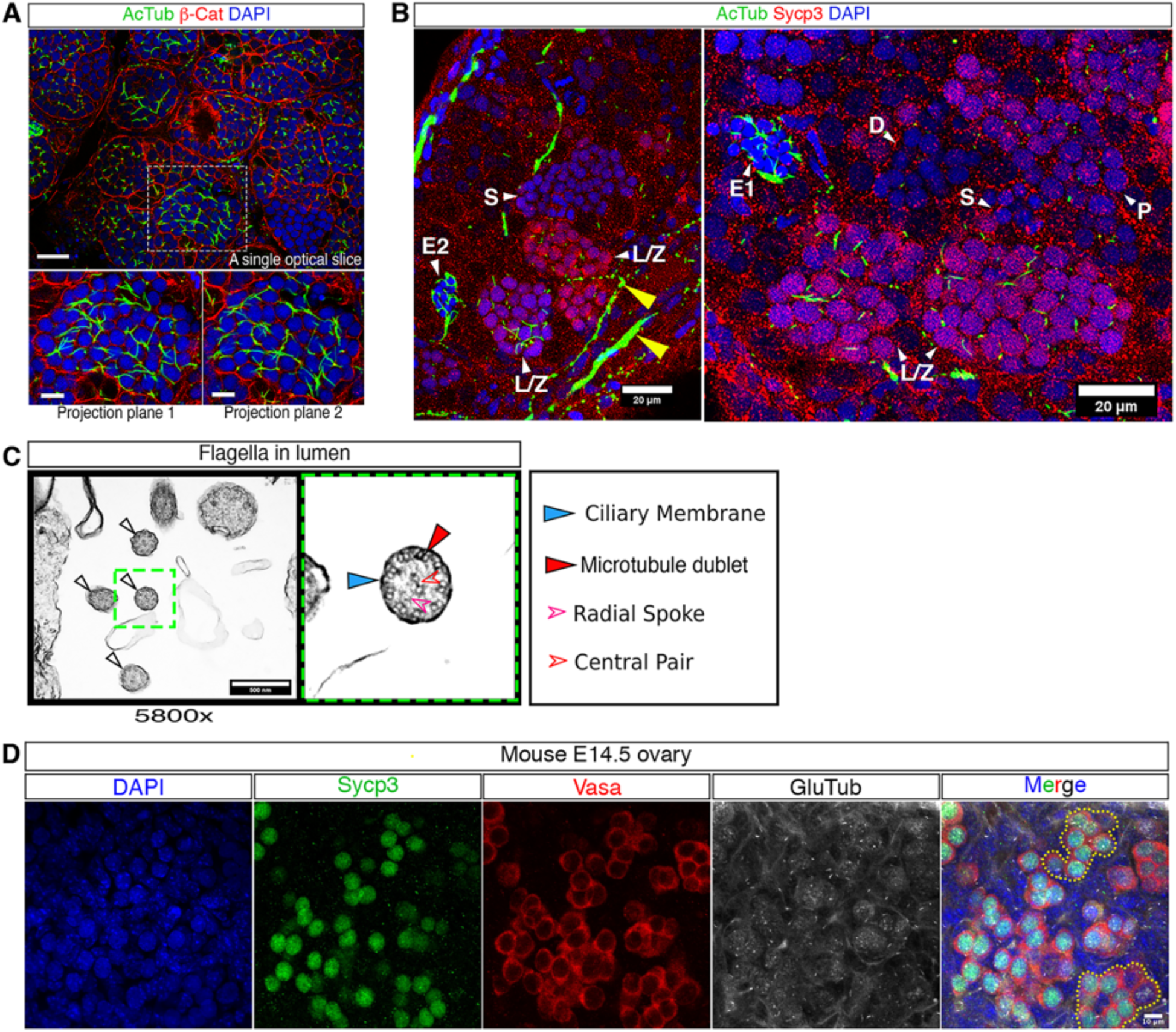
The zygotene cilia in juvenile testes, in TEM in adult testes, and in mouse oocytes. **A.** Juvenile testes labeled with AcTub (green), β-catenin (cytoplasmic membranes, red), and DPAI (blue), show cilia in young seminiferous tubules. Top panel shows a single optical section, the bottom panels show partial Sum z-projections of the white boxed region in the top panel, at two Z positions. Scale bars are 25 μm in top panel, and 10 mm in bottom panels. n=17 seminiferous tubules in 2 testes **B.** Juvenile testes labeled with AcTub (green), Sycp3 (red) and DAPI (blue), show flagella in early forming spermatids, and confirm the AcTub labeled cilia in Sycp3-positive zygotene spermatocytes. Two example images are shown. Yellow arrowheads indicate AcTub in gonad axons. n=37 seminiferous tubules in 2 testes. Scale bars are 20 μm. **C.** TEM images of adult testis showing cross sections of flagella (empty arrows) within the lumen of seminiferous tubules. Right panel is a zoom-in image of the green box in the left panel. Axonemal membrane (blue arrowhead), microtubule doublets (red arrowhead), a central microtubule pair (empty red arrowhead), and radial spokes (empty magenta arrowhead) are shown. Magnifications and scale bars are as indicated. n=2 testes. **D.** Mouse E14.5 ovaries co-labeled with Vasa, Sycp3, GluTub, and DAPI. A representative ovarian image is shown (four channels individually and their merge on the right). Cyst images in Fig. 6K were taken from this and other such frames (n=2 ovaries). Outlined cysts in the merged panel correspond to the cysts shown in the left two panels in Fig. 6K. Scale bar is 10 μm.

**Supplementary Figure 19.**
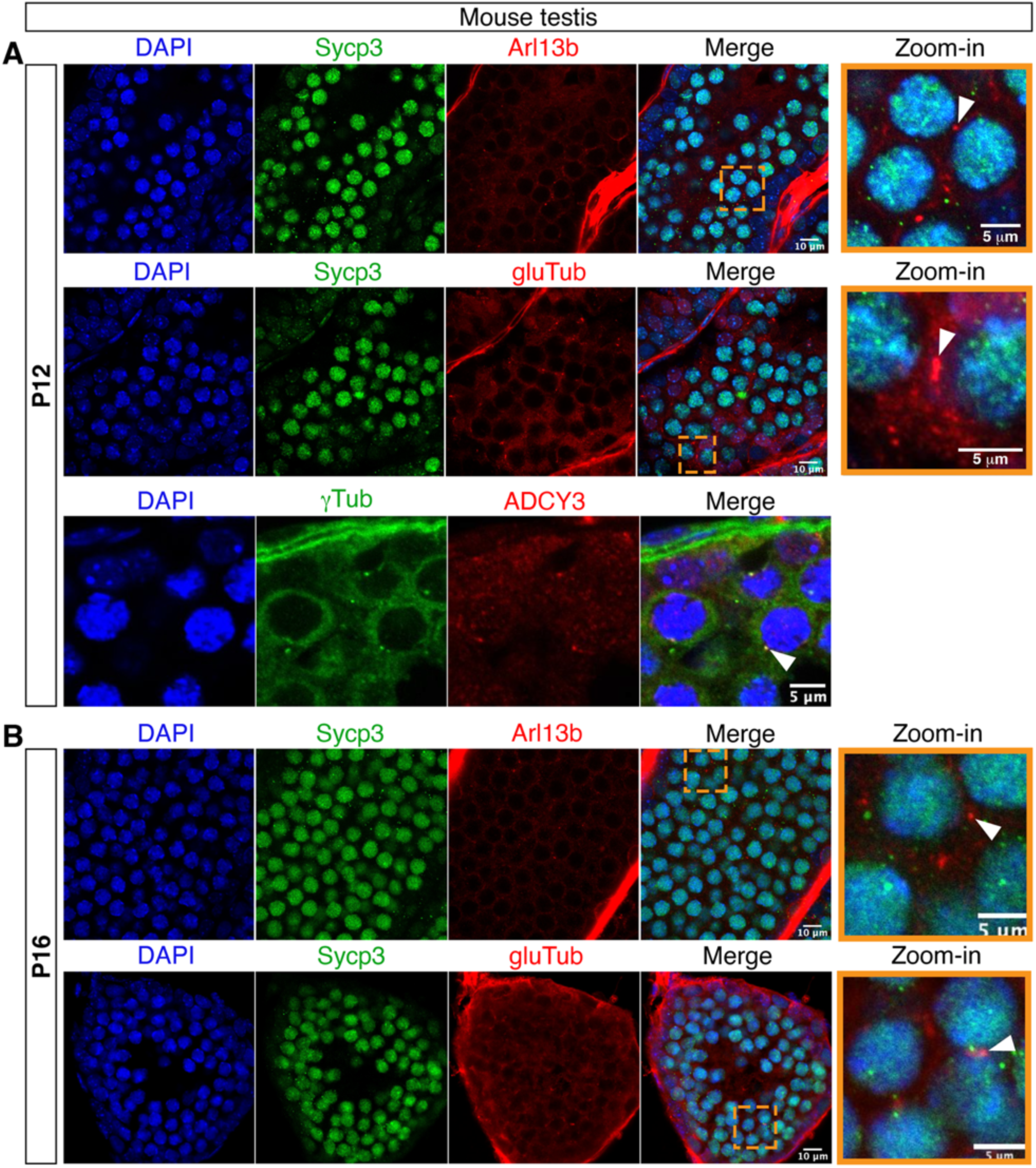
Ciliary structures in mouse spermatocytes. Thick (20 μm) cryo-sections of testes from P12 (A) and P16 (B) mice, co-labeled for Sycp3 and ciliary markers Arl13b and gluTub. P12 testes co-labeled for γTub and ADCY3 (bottom panels in A) shows localization of the ADCY3 ciliary protein to the centrosome basal body. Left panels are individual channels and their merge images. Orange boxed right panels are zoom-in images of the orange dash-boxed regions in their corresponding merge panel. n= 2 P12 and 2 P16 testes.

## Captions for Supplemental Movies

**Movie S1.** Live time-lapse recording of chromosomal dynamics (*Tg(h2a:h2a-gfp)*, grey) in a representative bouquet rotation oocyte (bottom), as well as oogonial- (middle) and somatic pre-granulosa (top) cells as controls (from Fig. S2A). All cells were co-recorded simultaneously in the same ovary and video. Each cell is shown without (left) and with tracks overlay (right; Methods). n=12 bouquet rotation oocytes, 13 oogonial cells, and 15 follicle cells from 4 ovaries. Scale bars are 5 μm.

**Movie S2.** Live time-lapse images of bouquet chromosomal rotations (Hoechst, red) and the centrosome (*Tg(bact:cetn2-GFP)*, green). n=5 ovaries. Scale bar is 10 μm.

**Movie S3.** 3D Rendering of a zygotene cyst SBF-SEM data, showing oocyte cytoplasm (color coded), nuclei (white), cilia (maroon) and centrosome (green).

**Movie S4-5.** Tracking of zygotene cilia in Z-stacks of SBF-SEM sections. Each cilium and its associated centriole and basal body is color-coded, numbered, and tracked throughout the z-stack. Different example are shown in Movies S4 and S5.

**Movie S6.** 3D Rendering of a zygotene cyst SBF-SEM data as in Movie S3, but stripped from cytoplasm, to show the cilia (maroon) tangling throughout the cyst. Nuclei (transparent grey), and centrosome (green) are also shown.

**Movie S7.** 3-D reconstruction of a zygotene bouquet oocyte labeled with DAPI, AcTub, and Telo-FISH. Fig. 1H is a snapshot from this video. n=3 ovaries. Scale bar is indicated.

**Movie S8-9.** 3-D reconstruction of zygotene bouquet oocytes labeled with EMTB-GFP (microtubules) and DAPI, showing perinuclear microtubule cables. Arrowhead in Movie S8 indicate the oocyte of which snapshot is shown in Fig. 1J. Additional examples is shown in Movie S9. n=7 ovaries. Scale bars are indicated.

**Movie S10.** Live time-lapse recording of the zygotene cilium [*Tg(β−act:arl13b-gfp),* grey], during bouquet chromosomal rotations (*Tg(h2a:h2a-gfp)*, grey) in a representative zygotene cyst (magenta outline). Oocyte cortex is visualized by actin [*Tg(β−act:lifeact-gfp),* grey]. Video images are focused on the optical sections of the annotated oocyte in a partial SUM projection. n=2. Scale bar is 5 μm.

**Movie S11.** Live time-lapse recording of the zygotene cilium [*Tg(β−act:arl13b-gfp),* grey], during bouquet chromosomal rotations (*Tg(h2a:h2a-gfp)*, grey) in a representative zygotene cyst (magenta outline). Video image is a larger SUM projection showing multiple cilia through the depth of the cyst. Cilia wiggle but do not exhibit beating or propelling movements. n=3 ovaries. Scale bar is 10 μm.

**Movie S12.** Live time-lapse recording of chromosomal dynamics (*H2A-GFP*, grey) in representative oocytes from *wt Tg(h2a:h2a-gfp);cep290^+/+^* (top) and sibling *Tg(h2a:h2a- gfp);cep290^-/-^* (middle and bottom) ovaries (from Fig. 3A). Each cell is shown without (left) and with tracks overlay (right). Two *cep290^-/-^* oocytes are shown: a representative oocyte that exhibited average track velocities (middle panel) and a low outlier oocyte with track velocities below the lower SD, which still exhibit rotation movements. wt n=21 oocytes from 4 ovaries, and *cep290^-/-^* n=73 oocytes from 4 ovaries. Scale bars are 5 μm.

**Movie S13-14.** Laser ablation of the zygotene cilium in live time-lapse imaging of a double transgenic line labeling cilia and centrosomes [*Tg(βAct:Cetn2-GFP);Tg(βAct:Arl13b-GFP)*]. Teal arrowhead indicates the ablation targeted cilium before ablation (pre-ablation), and its associated centrosome is circled in red. Yellow rectangle indicates the ablated region at the base of the targeted cilium. Centrosomes associated with other non-ablated cilia are circled in green. Time is indicated in mili-seconds. Note the ciliary-ablated red circled centrosome is stationary and then immediately dislocates upon its ciliary ablation, as it moves in the XY axes and in the Z (“centrosome out of focus”), while non-ciliary ablated centrosomes remain stationary. Movie S13 and S14 each shows a representative example. n=17 cilia (oocytes), from 11 ovaries.

**Movie S15.** Ciliary Laser ablation as in Movies S13-14, except high laser power is used and leads to cells death, showing that only higher laser power than what used in Movies S13-14 is toxic. n=11 oocytes from 9 ovaries.

**Movie S16.** Laser ablation as in Movies S13-14, except high laser power is used and targeted to the cytoplasmic membrane and leads to cells death, showing that laser ablations in Movies S13-14 precisely and specifically excise the zygotene cilium without damaging the cell. n=5 oocytes from 9 ovaries.

**Movie S17.** Quantitative analysis of cyst phenotype categories. A video demonstrating the 3D measurements of distances between neighboring nuclei in germline cysts, as described in the Methods section. Briefly, ROIs of cyst labeled with DiOC6 (cytoplasm) and DAPI are constructed in 3D and their nuclei rendered using IMARIS, identifying their centers of masses. Distances between centers of masses of all neighboring nuclei are measured and exported for further normalization to oocyte sizes and for statistical analysis.

